# Hiding in Plain Sight: CD8+FOXP3+Tregs sequester CD25 and are enriched in human tissues

**DOI:** 10.1101/2023.12.24.573232

**Authors:** Lorna B. Jarvis, Sarah K. Howlett, Daniel B. Rainbow, Valerie Coppard, Ricardo Ferreira, Ed Needham, Aaditya Prabhu, Sarah Alkwai, Lou Ellis, Zoya Georgieva, Ondrej Suchanek, Hani Mousa, Krishnaa Mahbubani, Kourosh Saeb-Parsy, Linda S. Wicker, Joanne L. Jones

## Abstract

For decades, regulatory T cell (Treg) research has focussed on CD4+FOXP3+ Tregs, while characterisation of CD8+FOXP3+ Tregs has been limited due to their scarcity in blood. Here, by analysing 95 tissue samples from 26 deceased transplant organ donors we show that, despite representing less than 5% of circulating Tregs, CD8+ Tregs are enriched in human tissue, particularly in non-lymphoid tissues and bone marrow. We further show that they are fully demethylated at the FOXP3 TSDR, indicating lineage stability, and demonstrate their presence in human thymic tissue and cord blood. Transcriptomic profiling revealed strong similarities to CD4+ Tregs, however at the protein level, they reside in tissue as surface CD25^lo^/-CD8+CD69+CD103+TLR9+HELIOS+FOXP3+ cells, expressing CD25 intracellularly. Surface CD25 was rapidly regained *ex-vivo*, allowing us to sort and expand them, and to subsequently demonstrate their therapeutic potential in a humanised mouse model of graft-vs-host disease. Additionally we report increased circulating CD8+Tregs in individuals with SLE and patients early following traumatic brain injury (TBI), underscoring their functional importance. We conclude that these under-studied cells likely play an essential but previously unappreciated role in maintaining peripheral tolerance.

**One Sentence Summary:** FOXP3+CD8+ Tregs, expressing tissue residency markers and intracellular CD25, are enriched in human non-lymphoid tissues.

## Main Text

Naturally occurring CD4+ Tregs are a specialised subset of CD4+ T cells that play a vital role in regulating the immune system. They are selected in the thymus to self-antigen expressed on MHC-Class II molecules and are capable of suppressing harmful immune responses to self in the periphery, where they play a key role in preventing autoimmunity and maintaining peripheral tolerance (reviewed (*1*)). Tregs can be distinguished from transiently activated T effector cells expressing FOXP3 in humans by the methylation status of the TSDR (Treg-specific demethylated region) located in intron 1 of the *FOXP3* gene, which is demethylated in Tregs promoting stable FOXP3 expression(*2*). CD4+ Tregs have been extensively studied over the past decades and their therapeutic potential is now being explored - to date over 50 clinical trials of Treg therapy have been completed or are ongoing exploring their use in the treatment of autoimmune disease and transplantation (according to www.clinicaltrials.gov clinical trials, reviewed in (*3*)).

Multiple CD8+ T cells with suppressive properties have also been described, both in mice and humans, including CD8+ cells that are CD2S^lo^, CD2Slo/CD57+(*4*), CD122+(*5*), CD45RC^lo^(*6*), KIR+ (in human)(*7*), Ly49+ (in mice)(*8*) and CD56+/NKT-like (in mice)(*9*). Whilst the CD2Slo and CD45RC^lo^ populations have been shown to contain a small subpopulation of FOXP3 expressing cells, the majority of these reported suppressive CDS+ T cells do not express FOXP3 and are thought to suppress via different mechanisms including cytotoxicity (reviewed in (*10*)). There have also been descriptions of CD8+FOXP3+ Tregs, primarily in the context of human tumour microenvironments where increased numbers correlate with worse outcomes *(11, 12)*. They are also known to be highly responsive to IL-2 (demonstrated in both humans and mice *in vivo)* (*13, 14)* and human CD8+FOXP3+ Tregs can be expanded *in vitro* using DC(*15*) or polyclonal stimulation(*6*). Yet due to their very low frequency in peripheral blood, few in-depth studies have been performed on *ex vivo* CDS+FOXP3+ Tregs from healthy individuals and their origin and role in human immunity remains poorly understood (reviewed in(*16*)).

In this study we undertook a detailed analysis of the CD8+FOXP3+ T cell compartment in humans. By integrating high-parameter flow cytometry, bulk and single-cell transcriptomic profiling, TSDR methylation analysis, in vitro and in vivo suppression assays, and by studying tissues from deceased transplant organ donors we reveal, for the first time, that human CD8+FOXP3+ cells are highly suppressive, TSDR demethylated regulatory cells that are enriched within the Treg compartment of human tissues compared to peripheral blood, particularly in non-lymphoid organs such as gut and liver. Furthermore, we identify increased proportions of circulating CD8+ Tregs in individuals with systemic lupus erythematosus (SLE) and in patients shortly following traumatic brain injury (TBI), underscoring their functional relevance.

Collectively our findings suggest that CD8+FOXP3+ regulatory cells play a previously unappreciated role in maintaining tissue health in humans, and we propose that these cells hold promise for tissue specific Treg cellular therapy.

## Results

### CD8+FOXP3+ Tregs are naturally occurring FOXP3 TSDR-demethylated counterparts of CD4+FOXP3+ Tregs

To understand the nature of FOXP3 expressing CD8+ T cells in relation to previously reported CD8+ suppressive populations, we undertook deep phenotyping of CD8+FOXP3+ T cells in human peripheral blood, analysing both classical markers of naturally occurring CD4+FOXP3+ Tregs and markers associated with other reported suppressive CD8+ populations.

In agreement with previous studies, human peripheral blood CD8+FOXP3+ T cells were TCRαβ+ (Fig. 1a). They had a similar expression profile as their CD4+FOXP3+ counterparts (CD25hi, CD127lo, HELIOS+, CTLA-4+) but lacked markers of unconventional cytotoxic-like CD8+ Tregs (KIRs, CD56, CD122). They expressed low levels of perforin and granzyme B compared with non-FOXP3+ CD8+ T effectors and were CD28+ (Fig.1a). Whilst our data confirmed reports that most CD8+FOXP3+ T cells in human blood are CD45RC^lo^(*6*) (as are CD4+FOXP3+ Tregs), CD45RC status was not specific for CD8+FOXP3+ cells, but rather was a general feature of all memory cells (Fig. 1a and fig.S1).

**Figure 1.**
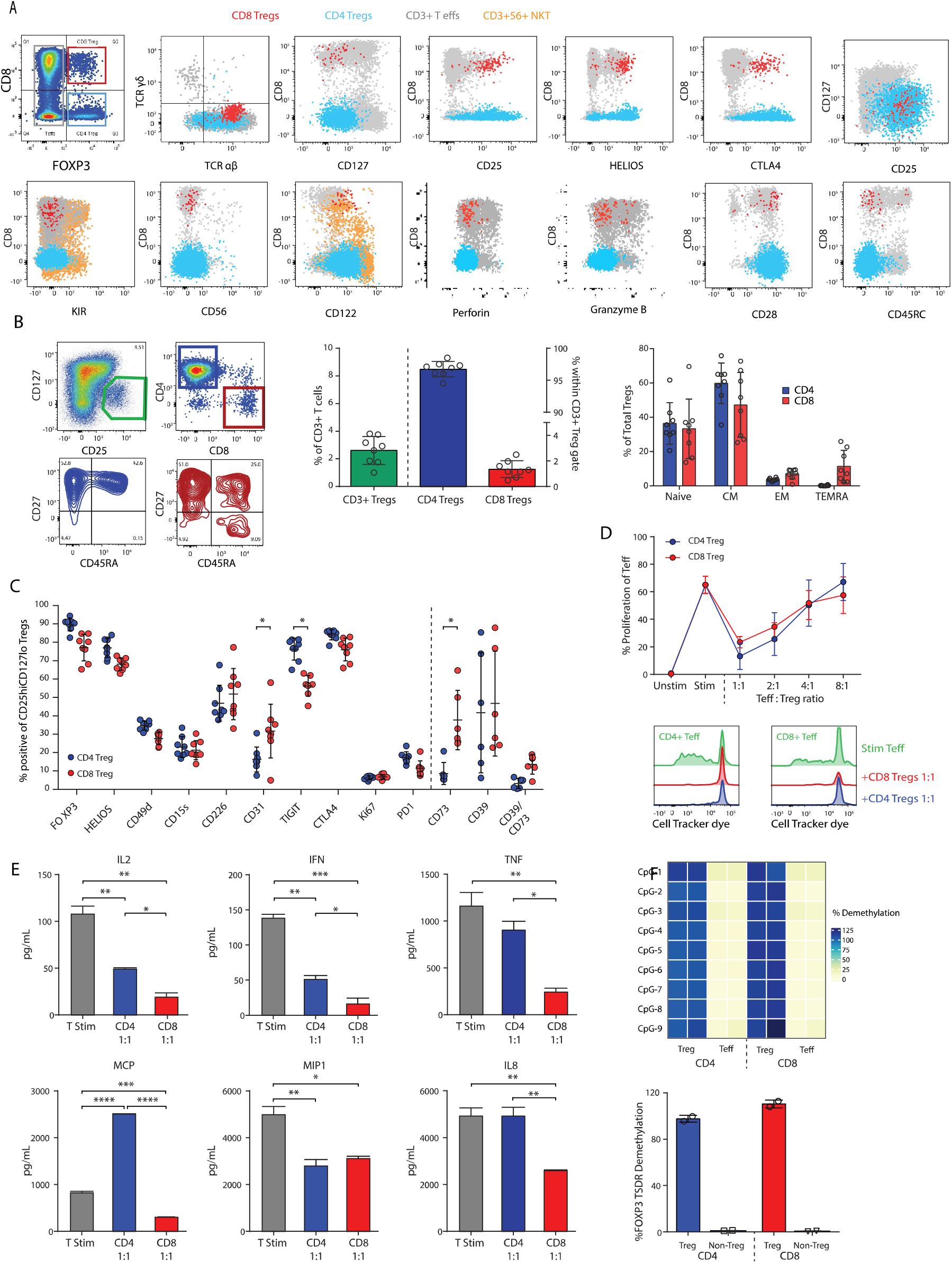
CD8+FOXP3+ Tregs in human blood are functionally suppressive and TSDR demethylated. A: Canonical and unconventional CD8+ regulatory cell markers by flow cytometry. Dotplot overlays show live - CD3+CD4+FOXP3+ Tregs (blue); CD3+CD8+FOXP3+ Tregs (red), FOXP3-CD3+ T cells (grey) and CD56+CD3+ NKT cells (orange). n = 3 donors B: The proportion of CD3+CD25hiCD127lo CD4+ (blue) and CD8+ cells (red) in human PBMC and the distribution of naive and memory subsets within those compartments based on CO2? and CD45RA expression. Representative gating and summary plots of n = 8 donors (mean +/− SD) shown C: Comparison of Treg canonical marker expression between CD4+ and CD8+ Tregs (gated on CD3+CD25hiCD127lo Tregs) in human PBMC. Percentage positive of total CD4+ (blue) or CD8+ (red) Tregs are presented forn = 8 donors. Data shows mean +/− SD and statistical differences were assessed using multiple T-tests with Holm-sidak correction method. D: Proliferation of stimulated and tracker dye labelled pan T (CD3+) effector T cells cultured with and without CD4+CD25hiCD127lo (blue) or CD8+CD25hiCD127lo (red) Tregs, FAG-sorted from human PBMC. Summary of 3 donors showing mean % proliferation +/− SD and representative histograms showing suppression of both CD4+ and CD8+ T effectors E: Cytokine production of stimulated human pan =(CD3+) T effector cells (grey) cultured with and without CD4+ Tregs (blue) or CD8+ Tregs (red). Data shows mean +/− SD, n=2 Comparisons were made using one way ANOVA with Tukey’s multiple comparisons test. F:TSDR demethylation of human *ex vivo* CD3+CD4+CD25hiCD127I0 and CD3+CD8+CD25hiCD127I0 Tregs sorted from PBMC (n = 4 donors).

As with CD4+ Tregs, CD25hi and CD127lo expression was the best surface phenotype for identifying CD8+FOXP3+ Tregs in blood. Around 72% of all circulating CD8+FOXP3+ cells could be identified using this gating strategy vs. 90% for CD4+FOXP3+ cells (cells outside this gate were predominantly CD25lo/negative - Fig. 1a and fig. S1) and 77% vs. 89% of CD8+ and CD4+ cells within the CD25hiCD127lo gate were FOXP3+. Within blood, approximately 2% of CD25hiCD127lo CD3 Tregs were CD8+ (Fig. 1b). This aligns with a previous study in psoriasis(*17*). Similar to CD4+CD25hiCD127lo Tregs, most circulating CD8+CD25hiCD127lo Tregs were either central memory (CD45RA-CD27+) or naive (CD45RA+CD27+), although, in keeping with their CD8 lineage, a greater proportion had a terminally differentiated effector RA (TEMRA) phenotype (Fig.1a, b). Overall, CD8+CD25hiCD127lo Tregs expressed similar levels of canonical Treg markers to their CD4+ counterparts (Fig.1c), with the exception of lower expression of TIGIT and higher expression of CD31 and CD73 (p <0.05 Fig.1c). Similar expression patterns were observed when gating on HELIOS+FOXP3+ Tregs (fig. S2).

Next, based on surface CD25hi/CD127lo expression, CD4+ and CD8+ Tregs were sorted from human blood for targeted transcriptomic analysis. These data demonstrated a high degree of similarity with differential gene expression largely confined to genes associated with CD4 vs. CD8 co-receptor expression (fig.S3).

Having established that CD8+CD25hiCD127loFOXP3+ cells likely represent naturally occurring CD8+ counterparts to CD4+ Tregs, we went on to test their suppressive function *ex vivo* using a miniaturised Treg suppression assay - required because of their low frequency in blood (Fig.1d). Surface-sorted circulating CD8+ Tregs were found to be highly suppressive, capable of suppressing CD4+ and CD8+ effector proliferation to the same degree as CD4+ Tregs (Fig.1d and fig.S4) but being significantly more effective in inhibiting pro-inflammatory cytokines production (Fig. 1e). Suppression was not mediated through cytotoxicity, as no increase in effector T-cell death was observed when co-cultured with CD8+ Tregs compared to CD4+ Tregs (fig. S4). Finally, we confirmed that *ex-vivo* blood-derived human CD127loCD25hiCD8+Tregs are TSDR demethylated, indicating that they are a stable regulatory population (Fig. 1f).

### TSDR demethylated and CD8+ lineage committed Tregs are found in human thymus

To investigate the presence of CD8+FOXP3+ Tregs in human thymus, we analysed thymic tissue from 4 adult deceased transplant organ donors. In addition to CD4+ Tregs, and developing CD4+CD8+ double positive (DP) Tregs, we detected a distinct population of CD8+ single positive Tregs comprising 6-8% of all TCRαβ+FOXP3+ thymocytes (Fig. 2a/b). Deep phenotyping revealed that these CD8+FOXP3+ Tregs were HELIOS+CD5hiCD69+TCRab+ mature cells and that they expressed high levels of the transcription factor RUNX3 and low ThPOK, consistent with thymocytes positively selected on MHC class I/peptide complexes (reviewed (*18*)) and thus committed to the CD8 lineage (Fig.2c).

**Figure 2.**
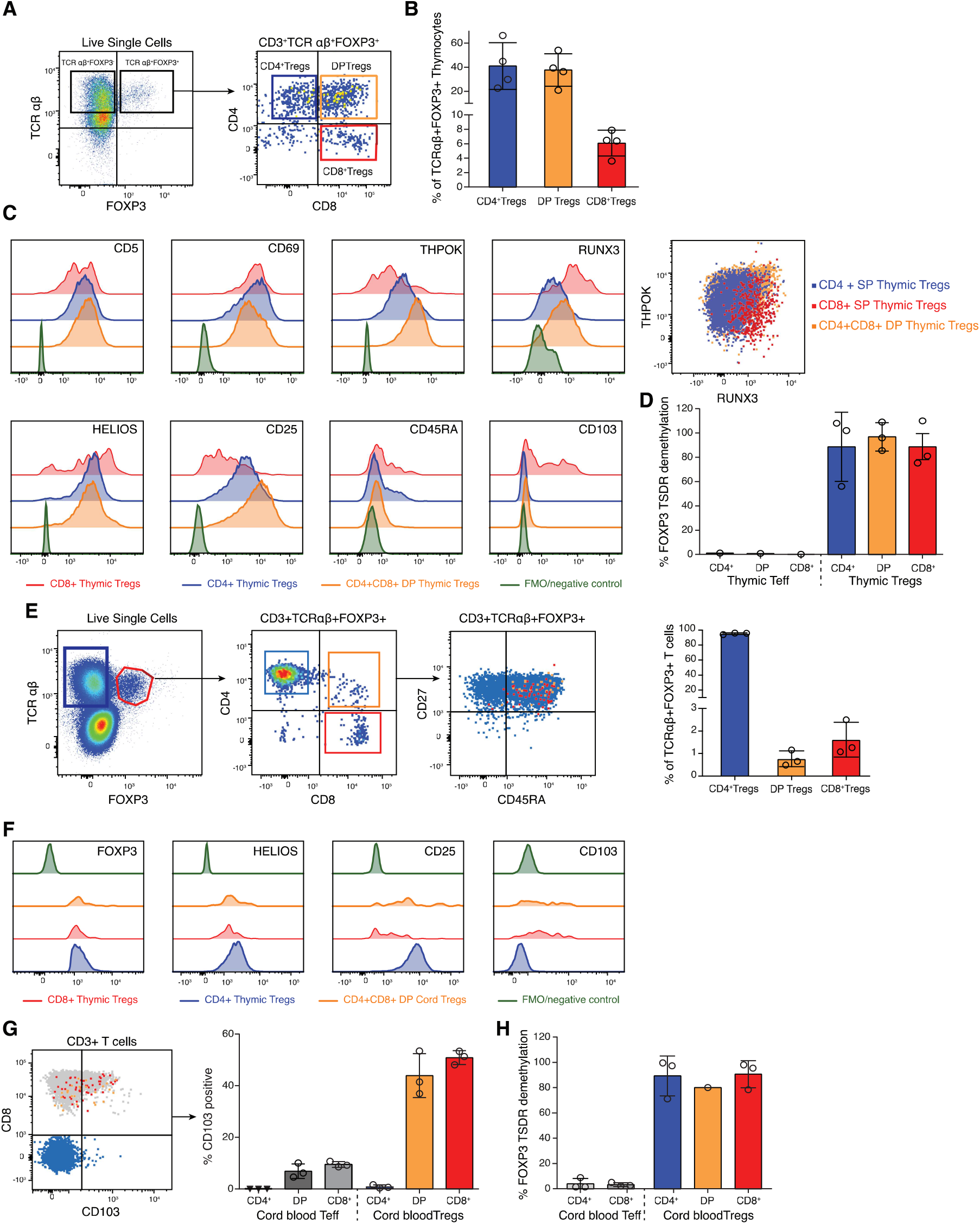
Fully demethylated CD8+FOXP3+ Tregs a exist in thymus and in early life in humans. A: Representative flow cytometric staining of TCRαβ vs FOXP3 on live lymphocytes and identification of CD4+, CD8+ and DP TCRαβ+FOXP3+ T cells in human thymus. B: Percentage of CD4+ SP (blue), CD4+CD8+ DP (orange) and CD8+SP (red) FOXP3+ cells in human thymus (n = 3 donors). Bars show mean +/− SD. C: Flow staining of canonical Treg and mature thymocyte markers and transcription factors THPOK and RUNX3 on CD4+ (blue), CD8+ (red) and DP+ (orange) FOXP3+ thymocytes compared with unstained or FMO controls (green). Representative example of n=3 donors. D: Histogram showing percentage FOXP3 TSDR demethylation in flow sorted TCRαβ+ FOXP3+ CD4+(blue), CD8+ (red) and DP (orange) thymocytes compared with FOXP3-Teffectors (mean values + SD are shown, summary of 3 donors). E: Representative dot plots showing CO27 vs. CD45RA expression on CD4+ SP (blue), CD4+CD8+ DP (yellow) and CDS+SP (red) TCRαβ+ FOXP3+ cells in human cord blood. Histogram show the percent of CD4+, CD8+ and F: Flow cytometric phenotyping of cord blood TCRαβ+FOXP3+ Tregs comparing CD4+ (blue), CD8+ (red) and DP+ (orange) with unstained or FMO controls (green). Representative example of 3 donors. G: Dotplot overlay shows expression of CD8 vs CD103 on CD4+FOXP3+ TCRαβ+ Tregs (blue), CD8+FOXP3+ TCRαβ+ Tregs (red) and CD4+CD8+ (DP) FOXP3+TCRαβ+ Tregs (orange), representative example of 3 donors. Histogram shows summary data for percentage CD103 expression within the Treg and Teff gates (n = 3 donors). H: Histogram showing % FOXP3 TSDR demethylation of CD4+(blue), CD8+ (red) and DP (orange) TCRαβ+ FOXP3+ Tregs, sorted from human cord blood, and their CD4+ and CD8+ effector counterparts (grey)mean +/− SD.; n=3

Of note, thymic CD8+FOXP3+ Tregs expressed lower surface CD25 than their CD4+ counterparts. They also expressed higher levels of the αE integrin CD103, compared to both CD4+FOXP3+ Tregs and CD8+ Teffectors (Fig.2c and fig. S6). Within the CD8+FOXP3+ population no correlation was observed between CD25 and CD103 expression (fig. S7) and CD103+Tregs within the thymus were found to be either CD8 single positive FOXP3+ Tregs, or CD4+CD8+ double positive (DP) Tregs that expressed RUNX3/ low ThPOK suggesting that DP CD103+ Tregs may be CD8+ lineage committed, and precursors of the CD8+ SP Treg (fig. S7).

To assess regulatory lineage stability, we flow-sorted CD4+, DP and CD8+FOXP3 positive and negative TCRαβ+ thymocytes and analysed them for TSDR demethylation - all FOXP3+ populations were highly demethylated (Fig. 2d). A small percentage of both CD4+ and CD8+ single positive Tregs were observed to express CD45RA (Fig. 2c), likely representing thymocytes closest to thymic egress(*19*).

Next we analysed umbilical cord blood (n=3), where we observed a small population of naive (CD27+CD45RA+) CD8+FOXP3+ Tregs (and also a very small DP FOXP3+ population) (Fig.2e). These cells expressed HELIOS, but in comparison to cord blood CD4+Tregs, had low CD25 expression and higher CD103 expression (Fig. 2f). CD103 was highly expressed on both CD8+ and DP FOXP3+Tregs compared to CD4+FOXP3+ Tregs and CD4+ and CD8+ Teffectors (Fig. 2g and fig. S8.). All three cord blood-derived FOXP3+ Treg subsets were highly demethylated at the TSDR, confirming their identity as stable regulatory cells (Fig. 2h).

Together with the T-cell activation marker CD69, expression of the αE integrin CD103 defines a recently identified subtype of T cells called “tissue-resident memory (TRM) cells”, found particularly in mucosal and barrier sites*(20, 21)*. We therefore questioned whether CD8+FOXP3+ Tregs are destined to home to tissues, and hypothesised that this may explain their paucity in blood. Supporting this, several genes associated with tissue homing, migration and residency were among the most differentially expressed between CD8+ Tregs and CD4+ Tregs, and between CD8+ Tregs and CD8+ effectors measured by Nanostring (despite them being sorted from blood, figs. S3 and S9). These included ITGAE (CD103), SIPR1, CD44, CD97 and other “effector” Treg (eTreg) markers associated with antigen experienced Tregs with tissue migratory potential as demonstrated in CD4+ Tregs in mice(22, 23) as well as a number of genes associated with tissue residency and tissue repair programmes in CD4+ Tregs (DUSP4, BATF(24–27)). We therefore sought to determine if CDS+FOXP3+ Tregs are enriched in tissues.

### CD8+FOXP3 TSDR-demethylated Tregs are enriched in human tissues

To explore the tissue distribution of regulatory T cells, we analysed 95 tissue samples from 26 deceased human transplant organ donors including: thymus, peripheral blood, thoracic and mesenteric lymph node, lung, spleen, fat, kidney, bone marrow (BM), liver, and gut (ileum and intraepithelial (IEL) layer and lamina propria (LP) of the jejunum) (Fig. 3a). Using multiparameter flow cytometry we found that, as a proportion of the CD127loFOXP3+ Treg pool, CD8+ Tregs are significantly enriched in tissues compared to peripheral blood (Fig. 3 b,c and fig. S10). In particular, liver, gut, kidney, and BM were highly enriched for CD8+ Tregs, where on average they accounted for 50%, 76%, 24.6% and 35.3% of all CD127loFOXP3+ cells compared to 4.4% in blood. In contrast, CD8+Tregs were not significantly enriched within the Treg pool in thymus, lymph nodes (LN), spleen, fat or lung, although variability across donors was seen.

**Figure 3.**
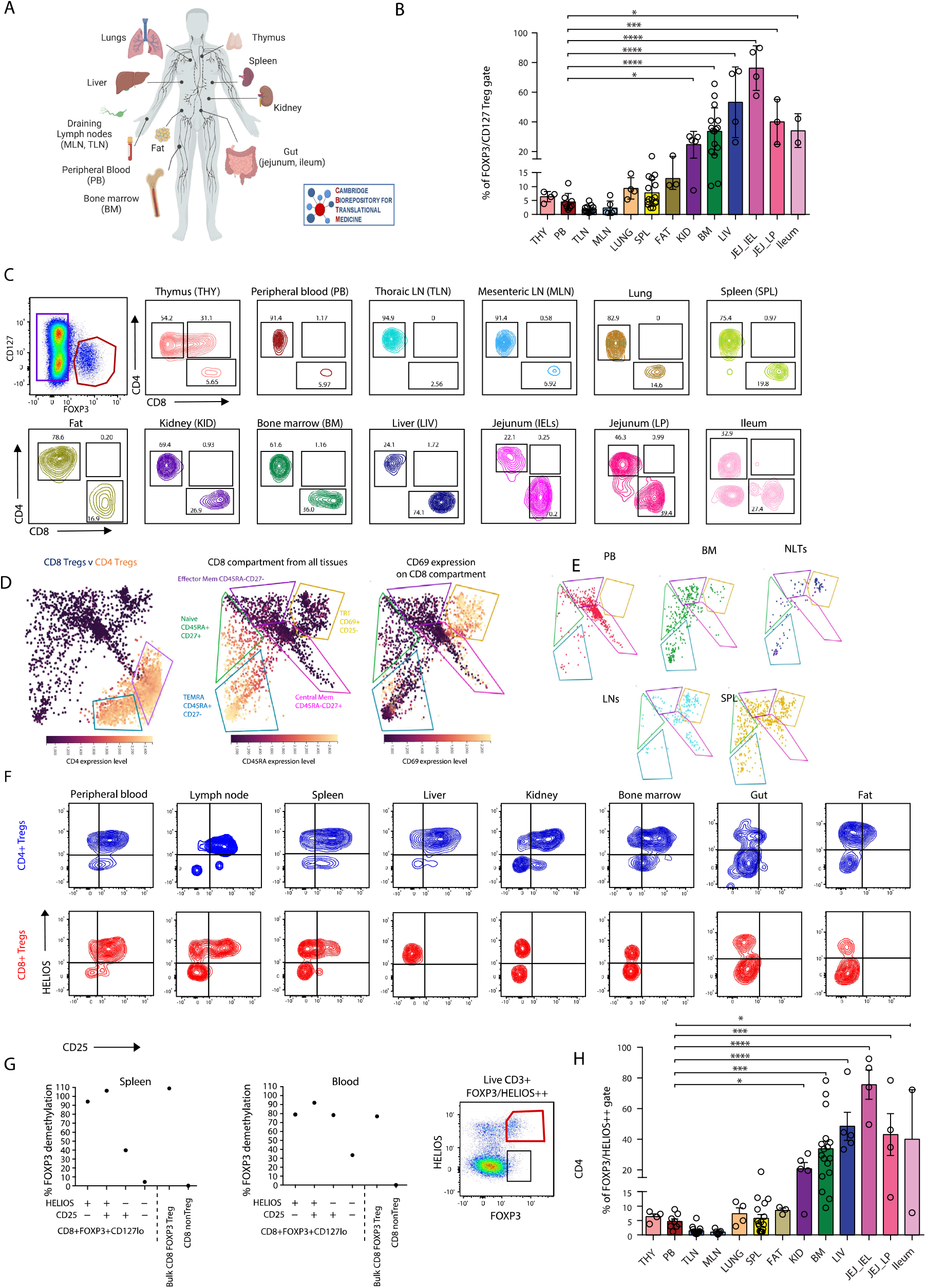
CD8+CD127loFOXP3+ Tregs are enriched in Human tissues and are TSDR demethylated. A: Schematic showing tissues collected by the Cambridge Biorepository for Translational Medicine (Cambridge) from 25 deceased organ transplant donors (image produced using Biorender). B: Frequency of CD8+ cells within the live, CD3+FOXP3+CD127lo Treg gate across multiple human tissues - (n = 4 Thymi (THY); 8 PBMC; 14 thoracic LN (TLN); 12 mesenteric LN (MLN); 4 Lungs; 16 spleens (SPL); 3 Fat samples; 5 Kidneys (KID); 17 bone marrow samples (BM); 5 livers (LIV); 4 Jejunum IELs (JEJ-JEL); 4 Jejunum Lamina propria (JEJ_LP) and 2 ileums. Bar charts and error bars show mean +/− SD with individual data points shown. Statistical differences were assessed using One Way ANOVA with Tukey’s multiple comparisons test. C: Example of CD127 vs FOXP3 staining on live CD3+ T cells and contour plots showing the proportion of CD4+ vs CD8+ Tregs across multiple tissues. D: FlowAtlas analysis of the T cell compartment of 40 tissues from 20 donors. Heatmap of CD4 expression level is shown (left plot) The middle heatmap shows CD45RA expression on the CD8+FOXP3+CD127lo compartment and the identification of 5 subpopulations within the CD8+ Treg pool. The right hand heatmap shows CD69 expression. E: CD8+ Treg subsets across human tissues by organ (peripheral blood = red; BM = green; non-lymphoid tissues (lung, kidney, liver) = purple; lymph nodes = blue and spleen = yellow) F: Contour plots showing CD25 v HELIOS expression in CD4+FOXP3+CD127lo (blue) and CD8+FOXP3+CD127lo (red) Tregs across human tissues (representative plots of at least 3 donors per tissue). G: Percent TSDR demethylation of CD3+CD8+FOXP3+CD127lo Treg subpopulations based on expression of CD25 and HELIOS sorted from human spleen and peripheral blood. H: Representative dotpot showing FOXP3 vs HELIOS staining of live CD3+ T cells and histogram showing percent of CD8+ cells within CD3+FOXP3+HELIOS+ Treg gate across multiple human tissues One Way ANOVA with Tukey’s multiple comparisons test.

These findings in part reflect the relative dominance of CD8+ T cells in certain tissues in our dataset (fig. S11), as has been reported previously *(28, 29)*. However, whereas CD4+ Tregs (as a percent of all CD4+ T-cells) are highest in blood (as well as thymus and lung), CD8+ Tregs are particularly underrepresented in blood - despite the reversal of the CD4:CDS ratio in the blood of our deceased organ transplant donors (Fig. 3 b,c and fig S11). No difference was seen in CD8+ Treg frequency between males and females. A trend towards increased CD8+ Tregs with age was observed in bone marrow and spleen, although this did not reach statistical significance (fig. S12).

Next we used FlowAtlas*(30)*, an interactive data explorer for dimensionality-reduced high-parameter flow cytometric analysis developed in-house, to assess Treg heterogeneity based on the Treg markers FOXP3, CD25, HELIOS, CD127, TIGIT, CTLA-4, and tissue resident markers CD69 and CD103 (figs. S13 and S14). Gut-derived samples were not included in this dataset.

Within the CD4+ Treg compartment two main subpopulations could be seen - naive (CD45RA+CD27+) and central memory Tregs (CD45RA-CD27+) (fig. S13). Broader heterogeneity was seen in the CDS+FOXP3+CD127I0 compartment, which could be segregated into five distinct subsets: (i) naive CD45RA+CD27+ cells, (ii) CM CD45RA-CD27+ cells, (iii) EM CD45RA-CD27-cells, (iv) TEMRA CD45RA+CD27-cells and (v) a CD45RA-population that expressed high levels of CD69 and increased levels of CD103 (Fig. 3d-e and fig. S14). This tissue resident population was absent from blood, rare in bone marrow, but seen in spleen and LN and was the dominant population in non-lymphoid tissues (nLT, including liver, lung and kidney) and thymus (Fig.3e and fig. S14). Blood from the deceased organ donors contained mainly CM cells; bone marrow contained TEMRAs, EM and naive cells; LN contained CM cells and tissue resident Tregs (TRTs) and spleen contained a mixture of all five populations (Fig. 3e). Whereas the CM population present in blood expressed CD25 and was mainly HELIOS positive, the other subpopulations showed variable HELIOS and low/absent CD25 expression. To better understand this, we further analysed HELIOS and CD25 expression in CD4+ and CD8+ FOXP3+CD127I0 cells across a range of human tissues (Fig. 3f). This showed that, whilst both CD4+ and CD8+ FOXP3+CD127lo cells in blood were predominantly HELIOS and CD25 positive, within non-lymphoid tissues (liver, gut, kidney, BM and fat) CD25 expression was low/absent on the CD8+ population, including on FOXP3+CD127I0 cells that co-expressed HELIOS. This was also the case for the CD4+FOXP3+CD127lo cells obtained from human gut and fat. Loss of CD25 was not caused by enzymatic tissue dissociation, since CD25 was retained on CD4+ Tregs and PBMCs treated with and without liberase and collagenase had comparable CD25 levels (fig. S15).

To understand the nature and origin of these populations, we next sorted FOXP3+CD127I0 CD4+ and CD8+ cells from blood, spleen and LN tissue from 3 different donors (where we had sufficient cells) by HELIOS and CD25 for TSDR methylation analysis. This demonstrated that regardless of CD25 expression, a high proportion of HELIOS+FOXP3+CD8+ cells are TSDR-demethylated, and are therefore, stable regulatory cells (Fig. 3g). This is in keeping with data previously published for CD4+ Tregs(*31*). The HELI0S-ve CD25-ve DN fraction of CD8+F0XP3+ cells from the spleen and blood contained a low proportion (20-40%) of TSDR-fully demethylated cells (Fig. 3g) compared to spleen- and blood-derived CD4 DN F0XP3+Tregs (fig. S16) where approximately 50% of the cells were fully demethylated at the TSDR. These populations are likely therefore, to contain a mixture of stable Tregs, peripherally induced Tregs and contaminating effector T cells. Given there is no consensus on how to distinguish these populations, we elected to re-calculate the fraction of CD8+ Tregs, as a proportion of total Tregs, across our donor tissues based on HELIOS and FOXP3 co-expression (fig. S17). Data showing the proportion of FOXP3+ CD4+ and CD8+ Tregs expressing HELIOS is also shown in fig. S18. Using this conservative definition of a Treg - CDS+FOXP3+HELIOS+, Tregs were still found to be highly enriched as a proportion of total Tregs in tissue, particularly non-lymphoid tissues (Fig. 3h).

### Further analysis of CD8+Tregs in single-cell and spatial datasets

To better characterise these cells, we performed 10X 5’ single-cell RNA sequencing on CD4+ and CD8+ Treg populations sorted from blood, spleen, kidney and liver tissues (n=9) and integrated the data generated with published immune single cell RNAseq datasets, both from our own group and from others *(17, 28, 32)*. CD8+ and CD4+ Treg enrichment was performed as these cell populations are usually poorly represented - even in large single-cell datasets.

In line with our flow cytometry analysis, we focussed on cells positively expressing either CD4+ or CDSA/CD8B+ and both FOXP3 and IZFK2 (HELIOS) as a stringent identifier of stable regulatory T cells. All other cells, not positively expressing these markers, but annotated as regulatory cells by CellTypist, are labelled as “other Regulatory T cells” and shown for comparison (Fig. 4a).

**Figure 4.**
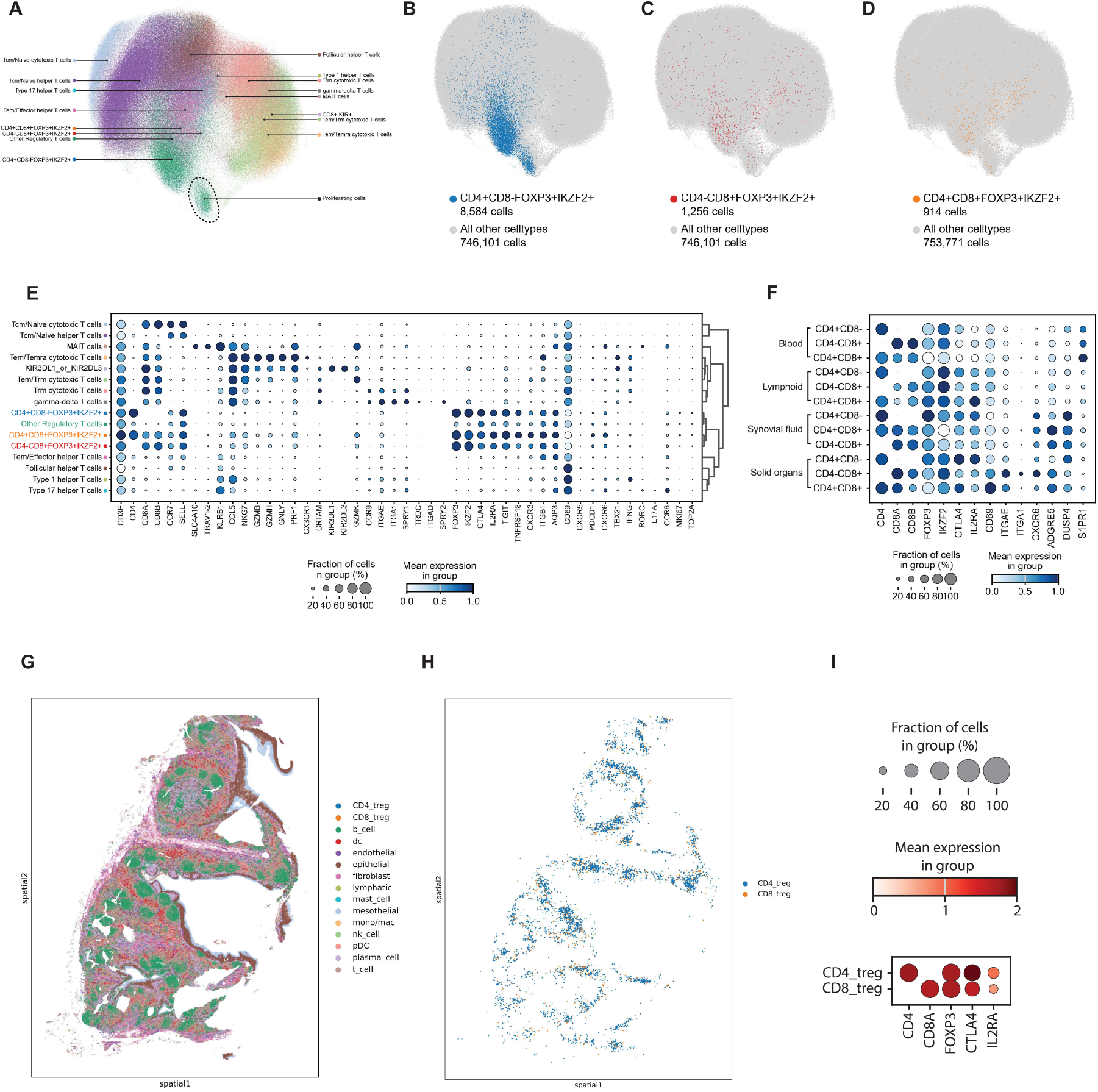
Identification of CD8+FOXP3+ and CD4+FOXP3+ Tregs in single Cell RNA sequencing data of Human tissues. T cells from four single cell RNA sequencing datasets generated from human tissues were combined to create a dataset of 754, 685 T cells(*17, 28, 32*) A-D: A combination of CellTypist and manual annotation was used to identify FOXP3+ IKZF2+ CD4+, CD8+ or CD4+CD8+ Tregs, along with the expected T cell subsets. The location of CD4+ (B), CD8+ (C) and CD4+ CD8+ (D) Tregs are shown on their own UMAP to aid visualisation. E: The expression of genes that defined each T cell population are shown as a dotplot, and are primarily based on the panel of genes used by Dominguez Conde (*28*), with the addition of some key genes identified in this study. F: The expression of tissue residency genes are shown as a dotplot for CD4+, CD8+ and CD4+CD8+ Tregs which are grouped tissue of origin. As the number of cells from some tissues is low, tissues were grouped as either lymphoid (lymph node, spleen and bone marrow) or solid organ Uejunum, liver and kidney). G: A Tonsil follicular lymphoid hyperplasia spatial transcriptomic data set from 10X Genomics was analysed for the presence of CD8+ Tregs. The data was annotated using the tonsil atlas from (*66*), and the annotations are projected back onto the tissue section. H: To aid visualisation, both CD4+ and CD8+ Tregs are plotted onto the section without any of the other celltypes. I: CD4+ and CD8+ Tregs were identified as expressing CD4+CD8A-FOXP3+ or CD4-CD8+FOXP3+ respectively, and these cells were enriched in the classical Treg genes CTLA4 and IL2RA (CD25), and their gene expression is shown as a dotplot.

CD4-CD8+FOXP3+IKZF2+ and CD4+CDS+FOXP3+IKZF2+ (CD8+ and DP Tregs) clustered together with regulatory T cells and tissue resident memory T cells (TRM) (Fig 4 a-d), suggesting their transcriptomic similarity. In keeping with our previous data, FOXP3+IKZF2+CD8+ Tregs transcriptomically closely mirrored FOXP3+IKZF2+CD4+ Tregs, and this was consistent across multiple human tissues (Fig 4e-f and fig, S19). *CD4* and *CDBA/CDBB* were the main differentially expressed genes between CD4+ and CD8+ Tregs (fig. S19–20). In addition, CD8+ Tregs expressed higher levels of *NKG7* and *CCL5*,consistent with their CD8 identity and long lived memory phenotype (*33, 34*). They also showed higher expression of cytotoxic genes, in particular granzymes, compared with CD4+CD8-FOXP3+IKZF2+ cells, although levels remained substantially lower than those seen in CD8+ cytotoxic effector populations (Fig 4e) and in keeping with our previous flow cytometry data (Fig 1). They had a distinct transcriptomic profile compared with recently described KIR+ tissue resident CD8+ Tregs(*7*), which as reported lacked FOXP3 and CTLA4 but expressed IKZF2 and cytotoxic genes. (Fig. 4e). Furthermore, and again consistent with our prior transcriptomic data (figs. S3 and S9), CD8+ Tregs in tissue and synovial fluid expressed high levels of CD69, ITGAE (CD103), DUSP4, ADGRE5 (CD97) and also CXCR6 and ITGA1 (canonical markers of tissue resident memory cells (*21, 35*)) (Fig 4f). SIPR1 was highly expressed in blood CD8+ Tregs (as observed in our earlier Nanostring data) but not in tissue, consistent with the dynamic roles of SIPR1 and CD69 in tissue migration and retention respectively (Fig. 4e) (*23, 36*).

To further investigate spatial localisation, we analysed publicly available 1OX Genomics Xenium spatial transcriptomic data from human tonsil tissue (*37*). This revealed both CD4+ and CD8+ FOXP3+ cells throughout the tissue with a similar distribution - clustering within follicles, (Fig 4g-i and fig. S20). Collectively these datasets confirm the existence of CD8+ Tregs within tissues and their transcriptomic and spatial similarities with CD4+ Tregs.

### CD8+FOXP3+ T cells in tissues express tissue residency markers, CD25 intracellularly and TLR9

Next we validated the expression of tissue residency markers at the protein level on CD4+ and CD8+ Tregs in our human tissue samples across multiple donors. This confirmed that CD8+FOXP3+HELIOS+ Tregs express higher levels of CD69, CXCR6 and CD103 compared to their CD4+ counterparts (Fig 5a-b and fig. S21). We also confirmed reduced perforin and granzyme expression relative to CD8+FOXP3-Teffectors (fig S.22).

**Figure 5:**
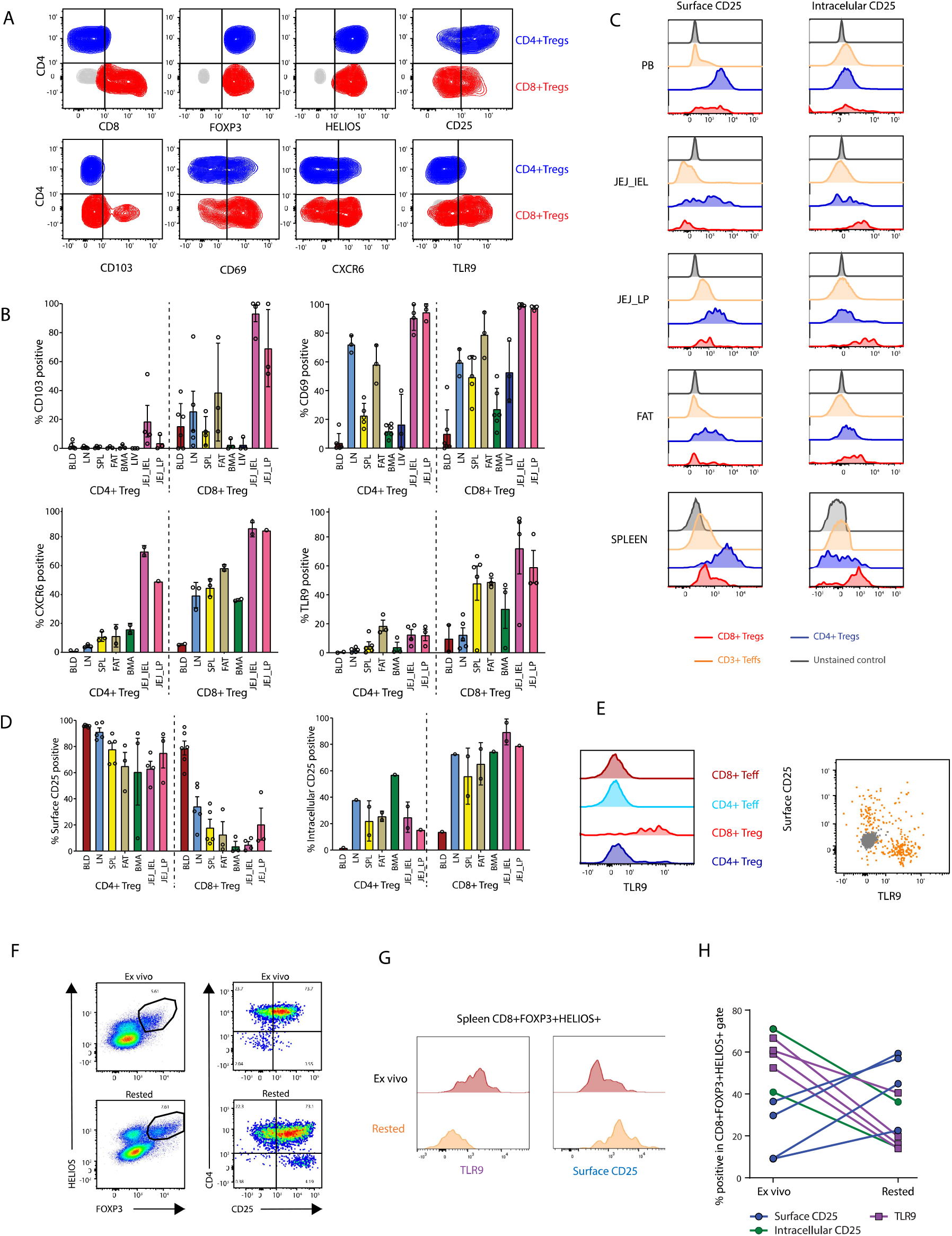
Tissue CD8+ Regulatory T cells express markers consistent with tissue residency, lack surface CD25 and co-express TLR9. A: Expression of tissue resident markers by CD4+ (blue) and CD8+ (red) CD3+FOXP3+HELIOS+ gated Tregs in human tissues. Contour plots show representative CD4 v canonical Treg and tissue resident markers. B: Bar chart showing percent CXCR6, CD69, CD103 and intracellular TLR9 positive cells within the CD4+ or CD8+ CD3+FOXP3+HELIOS+ gates (Bars show mean+-SD, for 2-8 donors per tissue) C: Flow histogram showing surface (left) vs Intracellular (right) CD25 staining of gated CD4+ (blue) and CD8+ (red) CD3+FOXP3+HELIOS+ Tregs across multiple human tissues. Grey histograms show unstained control and orange histograms show Teffector (CD3+Non-FOXP3+) cells. D: Summary of surface and intracellular CD25 expression of CD4+ and CD8+ CD3+FOXP3+HELIOS+ Tregs across multiple human tissues (24 tissues from 5 donors for Surface_CD25 and 10 tissues from 2 donors for intracellular_CD25). Bars show mean values+/− SD. E: Intracellular TLR9 expression by flow cytometry of gated CD3+CD4+ (blue) and CD8+ (red) FOXP3+HELIOS+ Tregs and FOXP3-Teffectors and an overlay dot plot showing TLR9 vs surface CD25 expression in CD3+FOXP3+HELIOS+ gated bulk Tregs (yellow) in a representative jejunum sample. FMO in grey. **F:** Dotplot showing surface CD25 expression on human splenic bulk CD3+FOXP3+HELIOS+ Tregs ex-vivo and after resting G: TLR9 and surface CD25 expression on gated CD3+FOXP3+HELIOS+CD8+ Tregs in human splenic MNCs stained directly *ex vivo* and after resting for 24hrs at t 37oC H: Summary data for surface CD25 (blue), intracellular CD25 (green) and TLR9 (pink) expression pre and post-resting for (n=2-4 donors).

We were intrigued by the absence of CD25 on the surface of FOXP3+HELIOS+CD8+ Tregs in tissues, despite its expression at the transcript level, and hypothesised that CD25 might be expressed intracellularly. To test this, we surface-stained cells extracted from a range of healthy human tissues with saturating concentrations of two anti-CD25 antibodies conjugated to one fluorochrome (B8515, clones MA2.1/2A3), then counterstained the cells post permeabilization with the same anti-CD25 clones conjugated to a different fluorochrome (APC), allowing us to distinguish surface and intracellularly expressed CD25. We observed that the majority of surface CD25 negative CD8+FOXP3+HELIOS+ Tregs in tissues express CD25 intracellularly. Of note, CD25 surface negative CD4+Tregs in tissue were also found to express CD25 intracellularly. CD25 intracellular expression was not observed in the effector population (Fig. 5c-d).

As lack of CD25 on the surface of tissue CD8+FOXP3+HELIOS+ CD8+ Tregs was preventing their isolation for downstream functional studies, we explored our transcriptomic datasets to try and identify differentially expressed genes between CD8+ Tregs and CD8+ Teffs, that might encode discriminating surface molecules. Candidates were then tested by flow cytometry (fig S23). Given that IRF4 and TRAF1 - both interferon response genes downstream of TLR signalling - were upregulated in CD8+ Tregs compared to CD8+ effectors, and given that TLR expression has been described in human Tregs, with TLR agonists shown to promote Treg expansion *in vitro* (*38*), we also screened for TLR expression.

Among the candidates tested, only TLR9 distinguished surface CD25-FOXP3+HELIOS+CD8+ Tregs from CD8+ Teffs (fig. S23), with expression levels comparable to that seen in CD79b+B-cells - where it is constitutively expressed (fig, S24), TLR9 was expressed in CD8+ Tregs across multiple human tissues (Fig 5B) and in gut we also detected in a minor population of CD4+FOXP3+HELIOS+ Tregs that lacked surface CD25 expression, and the reciprocal relationship between TLR9 and surface CD25 expression remained striking (Fig. 5E). These data suggest that TLR9 marks a subset of FOXP3+HELIOS+ Tregs lacking surface CD25. Since CD25-Tregs have been associated with recent activation we also assessed Ki67 expression, which was absent in this population (fig. S25).

TLR9 was not among the genes identified in our single cell data as being differentially expressed between CD4+ and CD8+ Tregs, however when expression across all T-cell subsets was assessed, low TLR9 expression was detectable in Tregs, consistent with other published reports((*38*)), but not in other T-cell populations (fig S20). Treg gene expression was, however, much lower than seen in B cells and pDC.

### Tissue resident CD8+ Tregs are TSDR demethylated, can be expanded ex-vivo and are functionally suppressive

Given that long-lived memory CD8+ T cells retain the ability to re-express key effector molecules(*39, 40*), we considered that CD8+ tissue Tregs might similarly regain surface CD25 expression upon removal from their tissue environment. To test this, splenocytes were isolated and rested *ex-vivo* for 24h at 37°C. As hypothesised, surface CD25 expression was regained. Notably, this was accompanied by a decrease in intracellular CD25 and a concomitant loss of TLR9 expression, supporting their reciprocal relationship (Fig.5f-h).

The recovery of surface CD25 expression ex-vivo also enabled the isolation of CD8+ Tregs from human tissues for TSDR demethylation analysis. CD4+ and CD8+ CD127loCD25hi Tregs were sorted from MNCs isolated from human spleen, LN and gut after resting overnight at 37oC (Fig. 6a-b). Consistent with our previous data, CD4+ and CD8+ Tregs recovered from spleen exhibited complete TSDR demethylated, as did cells from LN (Fig. 6c). In contrast, <50% of CD127loCD25hi CD8+ Tregs isolated from the gut were fully demethylated at the TSDR locus, a pattern similarly observed in gut-derived CD4+ Tregs (Fig. 6c). Of note both CD4+ and CD8+ FOXP3+ cells in the gut contain a large proportion of HELIOS negative cells, more than other tissues. Like their HELIOS+ counterparts these cells are negative for surface CD25 but are intracellular CD25+ and TLR9+ (Fig. 6d; and fig. S26). It is possible that these represent peripherally induced Tregs (pTregs). Given the very low number of cells obtained from the gut it was not possible to split the recovered CD25+ Tregs by HELIOS expression for TSDR analysis. Additionally, although gut contained the highest proportion of CDSaa FOX3+ cells of any tissue assayed (~13%), the vast majority co-expressed CDSa and CDSb (fig. S27).

**Figure 6:**
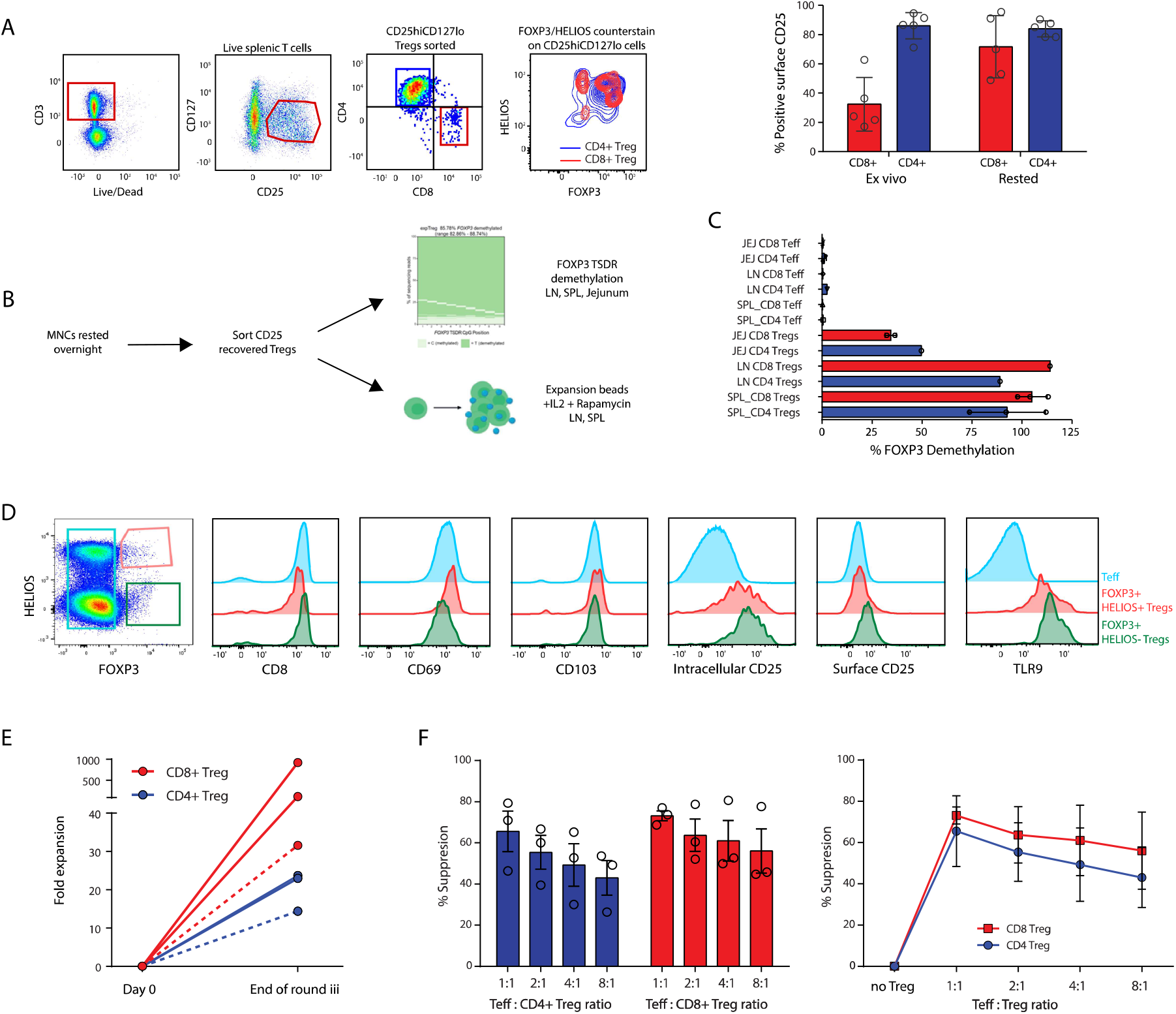
Tissue resident CD8+ Tregs (TRTs) regain surface CD25 expression and are functionally suppressive and TSDR demethylated. A: Example flow plots showing FAG-sorting of live CD3+_CD27lo_recovered surface CD25+ CD4+ and CD8+ Tregs after resting of splenic MNCs overnight at 37oC. The right hand contour plot shows intracellular FOXP3/HELIOS counterstaining of the sorted CD8+ Tregs (red) and CD4+ Tregs (blue) populations The bar chart shows summary data for the recovery of surface CD25 on 5 human spleens (bars show mean % surface CD25 expression +/SD. B: Schematic of protocol for isolating CD4+ and CD8+ Tregs from human tissues. Sorted Tregs were taken forward for TSDR demethylation analysis and, where cell number allowed, expansion in vitro. C: Post-surface CD25 recovery surface sorted CD3+CD25hiCD127lo CD4+ and CD8+ Tregs and CD3+CD25-CD4+ and CD8+ Teffector cells were assessed for TSDR demethylation. % TSDR demethylation in Jejunum (2 donors); LN (1 donor) and spleen (3 donors). Bars show mean +/− SD. D: Comparison of gut CD3+ FOXP3-Teffectors (blue), FOXP3+HELIOS+ Tregs (red) and FOXP3+HELIOS Tregs (green) for CD69, CD103, surface and intracellular CD25 and TLR9. Representative example of 2 donors E: Expansion of CD4+ (blue) and CD8+ (red) Tregs from 3 donors after 3 rounds of expansion either as flow sorted bulk CD3+CD25hiCD127lo human splenocytes (solid lines) or as separated CD4+ and CD8+. CD3+CD25hiCD 1271o populations (dotted lines). F: Suppression of proliferation of CT_efluor450 labelled and anti-CD3/28 stimulated pan T cells in co-culture with either CD4+ or CD8+ spleen-derived expanded Tregs. Bars represent mean +/− SD; n=3

**Figure 7:**
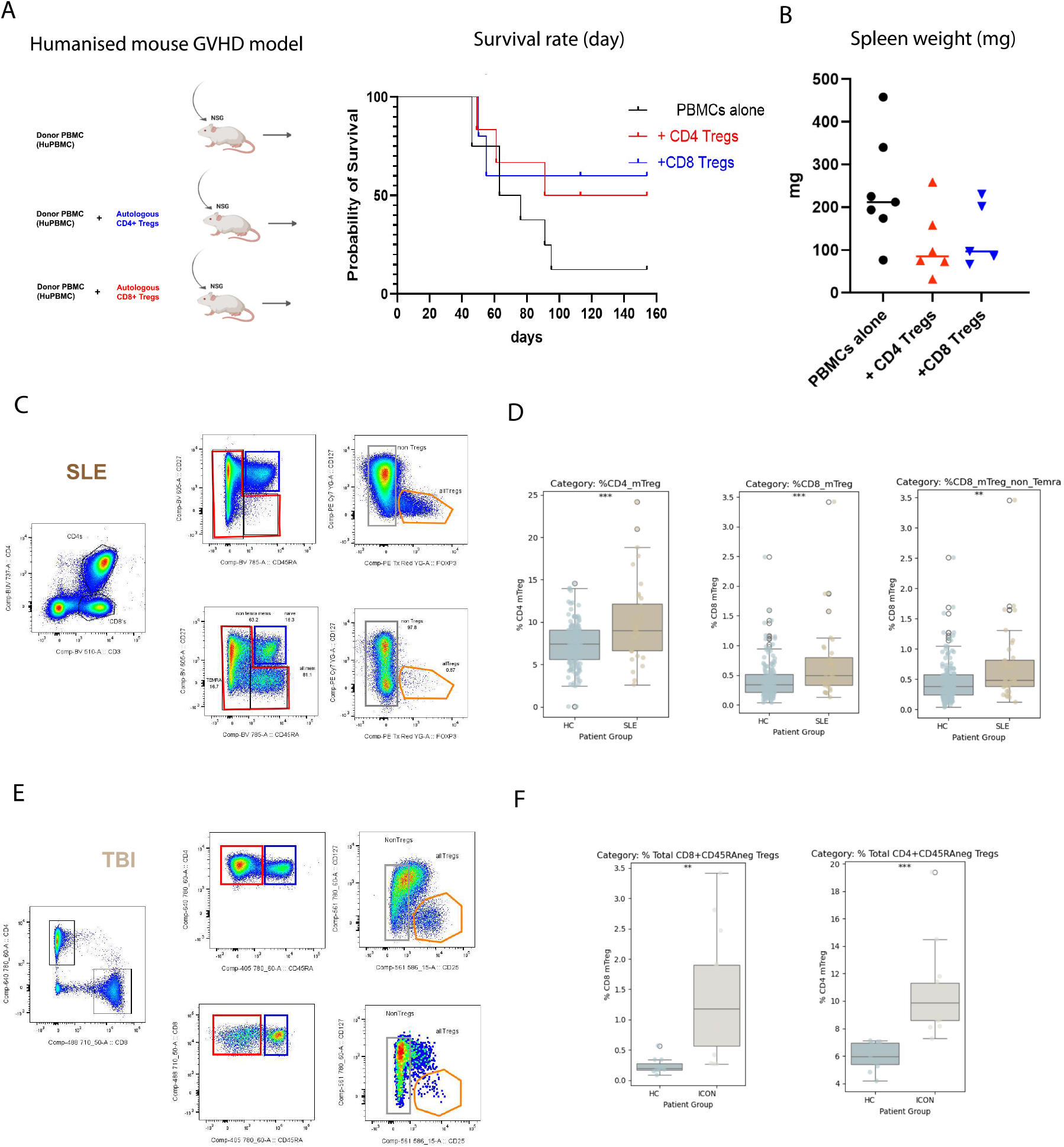
Clinical potential and relevance of CD8+ Regulatory T cells. A: Shows the experimental set up for humanising mice with total PBMCs (~ 5million per mouse) with or without expanded CD4+ Tregs or CO8+ Tregs (each ~ 5million) and the survival curves of the GVHD for each of the 3 sets of mice; PBMCs alone (black line; n=9)), with additional CD4+ Tregs (red; n=6) or CD8+ Tregs (blue; n=5) B: Shows the spleen weights at the time of sacrifice (mg). PBMCs alone (black; n=9)), with additional CD4+ Tregs (red; n=6) or CD8+ Tregs (blue; n=5). C: The gating strategy used to identify CD4+ and CD8+ memory Tregs from a previously published flow cytometric analysis of an SLE cohort(*31*). D: The frequency of CD4+ FOXP3+ or CD8+ FOXP3+ Tregs in either the healthy control (HC) or SLE cohort was plotted as a proportion of their parent memory population. CD8+ FOXP3+ Tregs were plotted as a proportion of total memory CD8+ T cells, or out of CD8+ memory T cells excluding TEMRA cells. E: The gating strategy used to identify CD4+ and CD8+ memory Tregs from a flow cytometric analysis of a traumatic brain injury cohort (TBI). The antibody panel used did not contain a marker to separate the naive/central memory population from effector/TEMRA population, therefore memory cells were gated solely as CD45RA-. FOXP3 was not on the antibody panel and Tregs were gated as CD127-CD25+. F: The frequency of CD4+ or CD8+ Tregs in either the healthy control (HC) or TBI cohort was plotted as a proportion of their parent memory population.

Next, CD127loCD3+ splenic Tregs (containing a mixture of CD4+ and CD8+ Tregs) with recovered CD25 expression were expanded using a standard expansion protocol involving TCR polyclonal stimulation in the presence of IL-2, and their expansion rate, phenotype and suppressive capacity after three rounds compared and contrasted. Despite comprising only a small fraction of the starting population, CD8+ Tregs expanded more rapidly than CD4+ Tregs in bulk cultures (Fig. 6e). They also maintained a stable suppressive phenotype, including surface expression of CD25 (HELIOS+FOXP3+CD127lowCD25hiTLR9−) (fig. S28). While CD39 expression was similar between expanded splenic CD4+ and CD8+ Tregs, CD73 expression - and consequently CD39/CD73 co-expression - was significantly higher on CD8+ Tregs (fig. S28). We have previously reported that co-expression of the ectonucleotidases CD39/CD73, which together convert extracellular ATP to immunosuppressive adenosine, is a key mechanism by which therapeutically expanded Tregs suppress(*41*)). After 3 rounds of expansion, CD4+ and CD8+ splenic Tregs were re-isolated from the bulk cultures and tested in a standard in vitro Treg suppression assay. Splenic CD25-recovered CD127loCD8+Tregs demonstrated high suppressive function, exceeding that of splenic CD4+ Tregs from the same donor (Fig 6f). Suppression was not observed with bulk donor-matched CD8+ T effector cells (fig S29).

### The clinical importance of CD8+ Tegs

Having established their suppressive capacity in vitro, we next evaluated their therapeutic potential by testing CD8+ Tregs in a humanised graft-versus-host-disease (huGVHO) mouse model. Co-transfer of either CD4+ or CD8+ Tregs, from the same donor, significantly delayed GVHO onset (Fig 7a). Additionally, spleen weights in Treg treated mice were significantly reduced compared to controls, consistent with attenuated T effector expansion in presence of either regulatory cell subset (Fig 7b).

Next we reanalysed two in-house flow-cytometry datasets originally generated to investigate CD4+Tregs in patients with systemic lupus erythematosus (SLE) (*31*) and early following traumatic brain injury (TBI), both of which had shown increased frequencies of circulating CD4+Tregs compared to age and sex-matched controls. By re-gating the data to extract equivalent data on Tregs within the CD8+ T cell compartment, in both cohorts we observed a clear increase in the frequency of circulatory CD8+ Tregs (defined as CD8+FOXP3+CD127loCD45RA+/ in the SLE cohort and as CD3+CD8+CD25hiCD127lo in the TBI cohort, fig S30 and S31) compared with healthy donors, which paralleled the CD4+ Treg response (Fig 7c-d), suggesting that CD8+ Tregs follow similar immunoregulatory dynamics during inflammation.

## Discussion

The existence of CD8+FOXP3+ Tregs has been known for some time but their relative abundance in non-lymphoid organs and their potential role in tissue immunity has not been described before and is an important new finding. Lack of awareness of these cells is likely due to an historical focus on CD4+ Tregs given the rarity of the CD8+ Treg population in blood, the assumption that all Tregs express high levels of surface CD25 and the inherent difficulty of accessing healthy human tissues. An increasing emphasis on tissue immunology, improved access to tissues and the advent of single cell technologies allowing unbiased cell profiling, have enabled discovery of an underappreciated role for this regulatory population. The fact that tissue resident cells tend to accumulate with age (so reducing the likelihood of detecting these cells in young mice housed in sterile environments) and due to the.

Our early experiments demonstrated that circulating CD4+ and CD8+ Tregs express comparable levels of canonical Treg-associated molecules - transcriptomically and at the protein level. We also demonstrated that, like CD4+ Tregs, most CD8+ Tregs (sorted either on CD25 or FOXP3 and low CD127 expression) are demethylated at the TSDR, confirming lineage stability. In human thymus and cord blood, we identified a population of naive CD8+FOXP3+ Tregs, providing further evidence of their early-life development. Notably, these cells exhibited high CD103 expression, not observed on CD4+ Tregs or T effector populations, suggesting a predisposition for tissue homing. *ITGAE* (CD103) and several other genes associated with tissue homing and residency, including *DUSP4* and *BATF*(*24, 25*), were also observed to be differentially expressed in our peripheral blood nanostring dataset supporting this hypothesis.

To test the hypothesis that CD8+ Tregs home to tissue, we performed multi-parameter flow cytometry on 95 tissues obtained from 26 deceased organ donors. This extensive and rare dataset revealed that CD8+ Tregs (whether defined as FOX3+CD8+ cells or FOXP3+HELIOS+CD8+ cells) are relatively enriched in tissue - particularly non-lymphoid tissue and especially in the gut - where they exhibit broader memory phenotypes and greater expression of tissue residency markers, such as CD69, CD103 and CXCR6. Integration of our in-house flow-sorted Treg data with publicly available cross-tissue scRNAseq datasets (from our own group and others) confirmed the presence of CD8+Tregs in tissues and highlighted their close transcriptional resemblance to CD4+Tregs. Differentially expressed genes largely reflect their lineage (*CDBA, COBB)* or effector memory phenotype (*CCL5, NKG7*) and tissue residency phenotype (*ITGAE, ITGA1, CXCR6 and CD69*). Granzyme and perforin genes were also noted to be expressed at higher levels in CD8+Tregs compared to CD4+Tregs - however gene and protein expression was significantly lower than that observed in CD8+ effector populations, in keeping with their non-cytotoxic behaviour in in-vitro suppression assays.

Although found in all tissues, HELIOS negative cells accounted for ~50% of all CD127loFOXP3+ CD8+ and CD4+ cells in the gut, where <=50% of Tregs (sorted on the basis of CD127loCD25hi and CD8 or CD4 expression) were found to be TSDR demethylated. Whilst we were unable to split the recovered Treg population by HELIOS expression (due to very low cell number) these data support prior studies suggesting that HELIOS denotes a stable regulatory phenotype (*42, 43*). The FOXP3+HELIOS-population may contain a mixture of activated FOXP3 expressing effectors and peripherally induced Tregs, however recent work from Maja Susko and Ethan Shevach - claiming that FOXP3 is not acquired by human conventional T-cells activated *in vivo*(*44*) would support the latter, and would be consistent with the literature reporting the gut to be an important site of Treg induction, in mice and humans (reviewed in (*45, 46*)).

We also identified a striking loss of surface CD25 on CD8+FOXP3+ Tregs in tissue (unrelated to enzymatic dissociation). Why this should be is unclear, although other CD25-Treg populations, including functionally suppressive CD4+CD25-T-follicular regulatory cells (CXCR5/PD1hi) have been described in mice and humans(*47*). TSDR demethylated CD25lo CD4+HELIOS+FOXP3+Tregs have also been reported to accumulate in autoimmunity, where their numbers correlate with Treg cell cycle and activation(*31*). Here, we show that FOXP3+CD8+ Tregs retain CD25 expression intracellularly, in keeping with retained *CD25* expression, and that surface expression is gained following their removal from the tissue microenvironment. This was not explored in previous studies.

Additionally, we identified an inverse relationship between intracellular TLR9 expression and surface CD25 expression in both CD4+ and CD8+ Tregs. While the mechanistic basis for this remains unclear, it is possible that sensing microbial or self-DNA during the course of infection or tissue injury might be an important part of their function in preventing aberrant immune responses to self antigen (TLR9 has been shown to be able to sense self-DNA/DAMPS in the context of sterile tissue injury(*48–50*)).

In general little is known about TLR expression by human Tregs. Several studies have reported that TLR9 signalling in APCs has a limiting effect on peripherally induced Tregs(*51, 52*), but very little has been published on the role of TLR9 expression in Tregs themselves. In one study TLR9 was reported to be expressed by a subset of human IL-10 secreting CD4+ pTregs (induced in the presence of dexamethasone and Vlt03/Calcitriol), and signalling through TLR9 via CpGs inhibited their function by inhibiting IL-10 secretion(*53*). However, in another study, TLR9 expression was identified in *ex vivo* nTregs undergoing therapeutic expansion and TLR9 agonists were shown to promote their function(*38*). *In vivo* evidence from TLR9−/− mice supports a role for TLR9 in promoting Treg function, tissue migration and retention (*54*). Additionally, cobitolimod (which activates TLR9) was recently shown to promote immune regulation in a human colitis IBD trial with improved resolution of colitis(*55, 56*).

While our findings suggest that CD8+Tregs are a tissue-resident regulatory population akin to CD8+ TRMs, our data do not speak to the longevity of these cells in tissues.

To assess the potential clinical relevance of CD8+ Tregs, we analysed two in-house flow cytometry datasets derived from peripheral blood samples of individuals with systemic lupus erythematosus and patients one week following traumatic brain injury. In both cohorts, CD8+ Tregs were proportionally increased within the circulating CD8+ memory T-cell compartment. This expansion mirrored that of CD4+ Tregs, suggesting that CD8+ Tregs are subject to similar dynamics during inflammation. Additionally, multiple studies have reported increased frequencies of CD8+FOXP3^+^ Tregs within tumour microenvironments where their presence often associates with poor outcome. Together, these findings suggest that CD8+ Tregs likely contribute to the regulation of human immune responses in both autoimmune and malignant settings, and demonstrate the need for further investigation into their role across tissues and diseases.

Given their potent suppressive function, tissue tropism, and their potential to respond to antigen presented on MHC class I - expressed by nearly all cells - we suggest that CD8+FOXP3^+^. Tregs also represent a highly promising target for therapeutic development. To this end, we have shown that they can be (i) isolated from bone marrow (a clinically accessible site), (ii) expanded robustly *in vitro* while retaining their suppressive phenotype, and (iii) are effective in preventing graft-versus-host disease in a humanised mouse model. And, notably, despite their low starting frequency, CD8+ Tregs expand more rapidly in vitro compared to their CD4 counterparts. Although to date, Treg therapy has almost exclusively focused on CD4+Tregs there is a growing interest in harnessing CD8+ Tregs therapeutically. For example Guillonneau and colleagues have shown that human CD8+ Tregs expanded from CD45RClow CD8+ T cells have therapeutic potential despite containing a relatively low proportion of FOXP3+CD8+ T cells(*6*). This group and others have also demonstrated that engineering Tregs with chimeric antigen receptors that specifically recognise MHC Class I can promote immune tolerance of transplanted organs.(*57, 58*)

In conclusion - we report for the first time that highly suppressive, stable CD8+ FOXP3+ Tregs are enriched in human tissue, where they exist as the likely regulatory counterpart of TRMs. This novel finding has significant implications both for the development of novel therapies, but also for our basic understanding of human tissue health and disease.

## Acknowledgements

We thank the deceased organ donors, donor families, and the extended Cambridge Biorepository for Translational Medicine (CBTM) team for access to the tissue samples. We also thank David Menon, Edward Needham, Menna Clatworthy and Alasdair Coles for access to additional blood and tissue samples. We acknowledge and were grateful for the support received from the Cambridge NIHR BRC cell phenotyping hub (Dept. Medicine, University of Cambridge) for this project.

## Supplementary Materials

Materials and Methods

Supp Figs. S1 to S31

Table S1

Table S2

## Materials and Methods

### Ethics Statement

All work was completed under ethically approved studies. Human PBMCs were isolated from healthy volunteers after obtaining fully informed consent under CAMSAFE (REC-11/33/0007) or CAMREG (21/NS/0011) and from leukocyte cones or cord blood of healthy donors (NHSBT, UK) or from in-patients under REC 97/290. TBI samples were gathered as part of the ICON-TBI study, REC 13/EE/0119, SLE patients as previously described(*31*) under REC Ref: 07/H0718/49. Human tissues were obtained from deceased transplant organ donors via the Cambridge Biorepository for Translational Medicine (CBTM) (REC 15/EE/0152 REC: East of England-Central Cambridge Research Committee). Additional kidneys were provided under REC12/EE/0446.

### Cell isolation and Magnetic separation

Human PBMCs were isolated from heparinized blood and tissue derived mononuclear cells were isolated from mechanically dissociated and filtered human tissue using Ficoll density gradient centrifugation (Ficoll PaquePlus; GE Healthcare, Amersham) or 33% Percell (GE Healthcare) as previously described(*28*). Solid tissues were mechanically dissociated following enzymatic treatment with either Liberase TL (Roche) or collagenase IV (Sigma) where appropriate. For a detailed protocol please refer to protocols.io(*59*). Prior to flow sorting Tregs, untouched Pan T cells were enriched by negative selection using T cell isolation kit II (Miltenyi biotec). All donor tissue information is included in Table S1.

### Flow phenotyping and flow sorting

Cells were stained using a range of antibodies (BD, eBioscience, Biolegend) at 1:50 dilution unless otherwise stated, and blocked using 3% FCS or mouse serum in Faes buffer and/or Tru block monocyte blocker (Biolegend) as required. Intracellular staining was performed using the FOXP3 permeabilization staining kit (lnvitrogen) for 45 minutes at room temperature followed by staining of intracellular markers in 1x perm wash buffer for 45 minutes at room temperature. Oual-fluorochrome CO25 staining was performed by staining surface CO25 using two CO25 antibodies (clones MA2.1/2A3) on BB515 at saturating concentration, washing cells x 3 with Faes buffer to remove excess surface antibody and then staining intracellular CO25 post perm/fix using antibodies of the same two clones on a second fluorochrome (APC). Cell sorting was performed on either a FACS Aria (BD) or Influx (BD) cell sorter. Flow acquisition for phenotyping was performed on the Canto II or Fortessa LSR (BD Biosciences) and analysed using FlowJo v7.6.5-v10.0 (Tree Star Inc) or FlowAtlas(*30*). All antibodies and flow panels are detailed in Table S2.

For analysis of patient datasets, two in-house datasets underwent a robust and blinded reanalysis for both CD4+ and CD8+ Tregs using gating strategies shown in figs S30 and S31 (Flowjo) and data was subsequently exported and QC/ thresholding performed prior to unblinding.

### *Ex vivo* Treg isolation and expansion

Human *ex vivo* CD3+CD127^lo^CD25^+^CD4^+^. CD3+CD127^lo^CD25^+^CD8+ or bulk CD3+ CD127^lo^CD25+ were flow sorted from human blood, spleen, LN or gut MNCs rested overnight in 6 well plates at 37oC in X-vivo 15 continuing 1% human AB serum (or for 48 hour with 5U/ml IL2 added for the last 24 hours to support Treg viability), using a BD FACSAria cell sorter (BD Biosciences). Cells were collected for TSDR demethylation analysis and stored at −80 as cell pellets or expanded with 500-1000 U/mL human rlL2 (Miltenyi Biotec/Chiron) +/− 100 nM rapamycin (Miltenyi Biotec) and human Treg expansion kit anti-CD3/CD28 beads (Miltenyi Biotec/lnvitrogen), using x-vivo 15 cell medium (Lonza) tissue culture medium supplemented with L-glutamine, penicillin-streptomycin (both PAA Laboratories, UK), HEPES (Gibco, UK) and 1-3% human AB pooled serum and media exchanged with fresh IL-2 every 3-5 days. For additional expansion periods Grex cell culture flasks were used (Wilson Wolf). Beads were removed using MACSibead magnet (Milteny Biotec) prior to phenotyping or use in functional assays.

### Measurement of *FOXP3* TSDR methylation

The methylation status of the *FOXP3* TSDR was measured using the protocol previously published *(41)*. In brief, Tregs or control Teffector cells, were processed with the Qiagen Epitect Fast Bisulfite kit, which lyses the cells and performs the bisulfite conversion in a single step. First round PCR targeted 9 CpG sites within the *FOXP3* TSDR and a second round PCR added index sequences and illumina compatible ends allowing for sequencing on an lllumina MiSeq (2×300 bp reads). The percent FOXP3 TSDR demethylation was calculated as the proportion of sequencing reads containing 8 or 9 of the CpG sites within the TSDR being demethylated compared to the total number of sequencing reads. For female donors TSDR demethylation values were multiplied by two to account for the fact that one X chromosome is fully methylated due to X inactivation as FOXP3 is located on the X chromosome. Since there is a known bias in PCR efficiency of demethylated DNA(*60, 61*) this can lead to values of over 100% TSDR demethylation being measured in cells from some female donors. For Treg samples with less than 10,000 cells, flow sorted FOXP3-B cells were added to bring the total cell count to 10,000 (i.e. within the optimal cell number range for the methylation assay). The dilution factor was used to adjust the TSDR methylation value obtained.

### *In vitro* stimulation and suppression assays

For short term activation of sorted Tregs and Teffs, cells were cultured in RPMI complete media containing 10% FCS, Penicillin-streptomycin and Hepes +/− anti-CD3/CD28 stimulation (lnvitrogen Dynabeads) at a cell:bead ratio of 4:1. *In vitro* suppression assays were performed as previously described(*41, 62*), briefly: efluor670 (lnvitrogen) labelled Tregs (ex *vivo* or expanded) and cell tracker efluor450 (lnvitrogen) labelled MACS sorted Pan T cells (Miltenyi T cell isolation kit II) were co-cultured in 96V bottom plates (Greiner). Pan Teffs were seeded at 2×10^4^ cells per well in RPMI+ 3% human AB serum in the absence or presence of equal and titrated numbers of Tregs and stimulated using Miltenyi Tregs suppression inspector beads according to the manufacturer’s instructions. Stocks of allogeneic pan Teffs pre-labelled with efluor450 cell tracker dye were frozen in liquid nitrogen and used as standard Teffs for these assays. Proliferation was measured on day 4-5 using BD Fortessa HTS plate reader. V670^+^ Tregs were gated out to enable analysis of the effector population (CD4^+^efluor450^+^efluor670^−^ cells) and suppression was calculated by taking the ratio between the proliferating and non-proliferating populations as previously described(*41, 62*)

### In vivo Functional experiments

Treg function was tested *in vivo* using a graft versus host disease (GVHD) model in NODCg-Prkdcscidll2rgtm1Wjl/SzJ (NSG) (purchased from Charles River UK) humanised mice (*63*). Briefly, NSG mice were humanised (by intra peritoneal injection into un-irradiated mice) with 5 million human PBMC with and without co-administration of equal numbers of autologous CD4+ or CD8+ Tregs expanded from human peripheral blood. Mice were bled at regular intervals to monitor successful humanisation and length of survival determined for all conditions. Spleen weights were also determined at the end of the experiment and final measure of humanisation assessed by Flow Cytometry.

### Transcriptomic profiling by Nanostring

15,000 freshly-sorted *ex vivo* CD8+ Tregs, CD4+ Tregs and DP Tregs and equivalent effector cell populations were harvested and washed twice in PBS, cells pelleted and lysed in RLT buffer (Qiagen). RNA was extracted using Qiagen RNeasy micro kit and their transcriptome measured using the nanostring Human Immunology V2 characterisation panel (Nanostring technologies) using an nCounter prep station and digital analyser.

### Transcriptomic profiling by Single Cell RNA-sequencing

As CD8+ Tregs are rare, we utilised two single cell immune atlases we generated(*28, 32*) as well as a single cell dataset from AS and PsA (*17*). Additionally, we performed a number of in house sorts to enrich for both CD4+ and CD8+ Tregs from human tissue (blood (n=2), spleen (n=5), liver (n=1) and kidney (n=1)). Since CD8+ Tregs in tissues lacked surface CD25 expression, we allowed MNCs to rest overnight at 37oC and then sorted Tregs then next day on the basis of low CD127 and surface CD25 expression on the rested cells. This enabled CD8+ Tregs to regain surface CD25 expression for sorting.

To integrate the single cell data from these different sources, the Scanpy package (*64)* was first used to combine the cells from each dataset, and basic QC performed to remove cells expressing less than 600 genes and 800 transcripts. Integration was performed using Harmonypy, with the batch key being set as the donor+tissue. Following integration, CellTypist was used to annotate the cells by majority voting. To identify CD4+and CD8+ Tregs, cells were positively selected as expressing CD4+ CD8A-COBB-FOXP3+ IKZF2+ or CD4-CD8A+ or CO8B+ FOXP3+ IKZF2+, respectively, and named CD4 Treg or CD8 Treg in Fig. 4. Cells identified as expressing both CD4 and CD8 (CD4+ CD8A+ or COBB+ FOXP3+ IKZF2+) were also identified in the single cell datasets. However, these cells only represent a small proportion of all the cells identified as Regulatory T cells by CellTypist, as CellTypist uses majority voting to combine cell annotations, and does not rely on each individual cell expressing the Regulatory T cell markers. Therefore, in Fig. 4 cells have been annotated as CD4+ or CD8+ Treg by positive selection of markers, and the remaining cells annotated as Regulatory T cells.

### Spatial transcriptomic profiling

To assess the spatial location of CD4+ and CD8+ Tregs within tissue, we used a human tonsil follicular lymphoid hyperplasia FFPE section that has been analysed using the 10X Genomics Xenium analyser with a 377 gene multi-tissue and cancer panel. The data is available from 10X Genomics (https://www.10xgenomicscom/datasets/human-tonsil-data-xenium-human-multi-tissue-and-cancer-panel-1-standard). The tonsil section was segmented based on DAPI nuclear stain with nuclear expansion, and resulted in 864,388 cells with a median of 34 genes and 60 transcripts detected per cell. This data was analysed using the scanpy and squidpy packages *(64, 65*). The data was first quality controlled to remove lower quality cells, by applying a filter to remove cells with less than 17 unique genes detected and 30 transcripts - filtered at 50% of the median values, and resulted in removal of 17.5% of the cells. Next the standard scanpy pipeline was applied to the data, with the data being normalised to 200 counts (sc.pp.normalize_total(adata, target_sum=200) and log transformed (sc.pp.log1p(adata) and subsequently scaled (sc.pp.scale(adata, max_value=10)). PCA was calculated using 30 principal components, neighbors calculated using n_neighbors = 30 and UMAP embedding visualised. Cell type annotation was performed based on the Azimuth tonsil celltype gene sets which is based on the atlas from Massoni-Badosa et al(*66*). To identify CD4+ and CD8+ Tregs, cells were positively selected as expressing CD4+ CD8A-FOXP3+ or CD4-CD8A+ FOXP3+, respectively. IKZF2 (HELIOS) was not on the Xenium panel, but the cells identified as CD4+ or CD8+ Tregs were enriched for the expression of CTLA4 and IL2RA (CD25). Annotations were mapped back to the section using squidpy to allow the spatial visualisation of cell type annotations.

### Cytokine Analysis

Cytokines were measured in cell culture supernatants using luminex analysis (Biorad human 17-plex Bioplex kit) according to the manufacturer’s instructions. Briefly, cell culture supernatants were harvested either from expanding CD4+ and CD8+ Tregs or from suppression assay supernatants at day 5, and stored in −80 until used. Upon thawing, supernatants were used either neat or diluted 1:2, and plated in duplicates. 17 human cytokines were analysed simultaneously using the kit and run on the Luminex analyser (BIORAD), standard curves for each cytokine created and per cytokine concentrations calculated using the BioRad software.

### Statistics and Reproducibility

All flow cytometry data were analysed using FlowJo (version 10). Statistical tests were performed using GraphPad Prism 6.0 software (GraphPad Software Inc, California). Comparisons between groups of three or more were compared using a one way ANOVA with Tukey’s or Dunnet’s post-hoc multiple comparisons tests, as indicated. Comparisons of 2 groups were tested with either one-tailed or two-tailed Student’s t-tests, a two-way ANOVA with Bonferroni post-hoc multiple comparisons test, or multiple T-tests with Holm-Sidak post-hoc correction as appropriate. The number of biological replicates for each data set are given in figure Nanostring data were normalised using nSolver 4.0 software (Nanostring), background thresholding was performed at the mean+2SD of the negative probe values for each sample (range 15.2 – 36.4). Only probes with counts greater than the background threshold in at least 2 of the 3 samples were included in the analysis. Volcano plots were generated using the EnhancedVolcano R package(*67*) and heatmaps were visualised with the Bioinfokit Python package(*68*).

### Data Availability

All flow cytometry panels and donor information are available in Tables S1 and S2. All other raw data are available from the corresponding author upon reasonable request.

---

**Supplementary Fig 1.**
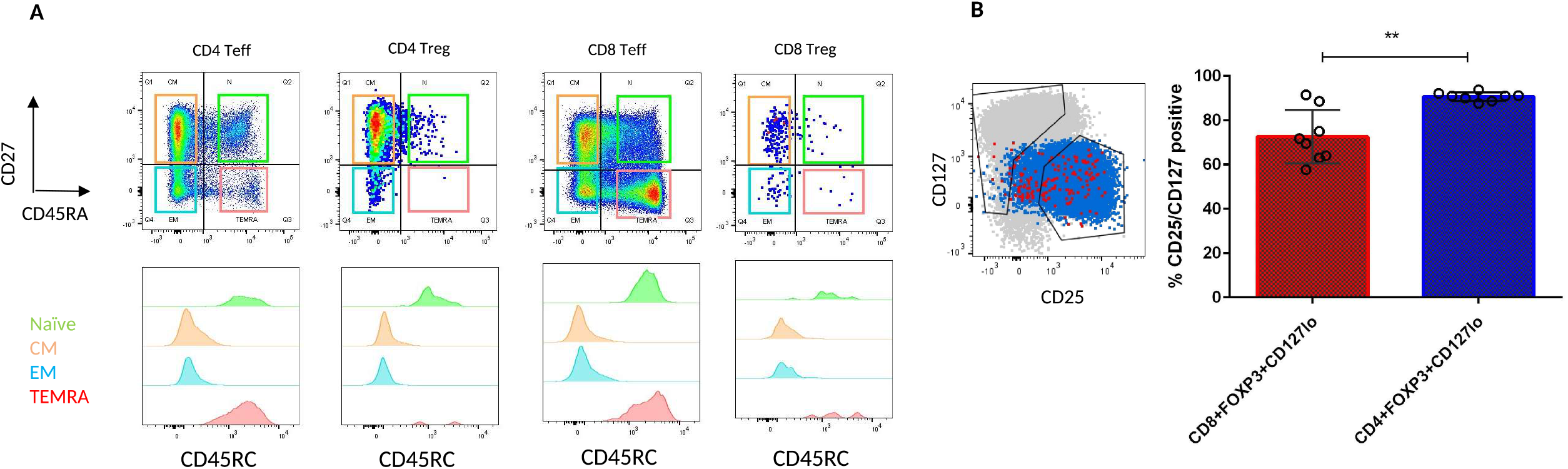
CD45RC v CD45RA and surface CD25 expression in blood. A: CD27 vs CD45RA staining on CD4 and CD8 CD3+FOXP3-T effectors and CD3+FOXP3+ Tregs (top panel) from human blood. Histogram overlays showing CD45RA expression levels in naïve (green), central memory (CM orange), effector memory (EM, blue) and TEMRA (red) cells. CD45RC expression was expressed with CD45RA on naïve and TEMRA T effectors and Tregs B: CD3+CD8+FOXP3+ cells (red) and CD3+CD4+FOXP3+ cells (blue) overlaid onto CD127 vs CD25 plot shown against CD3+ Teffector (grey); representative flow plot of 8 donors (left) and % of Total CD3+CD8+FOXP3+ (red) and CD3+CD4+FOXP3+ (blue) cells that are captured by the CD127loCD25hi gate in human PBMC: Summary data ofn = 8 donors showing mean +/− SD

**Supplementary Fig 2.**
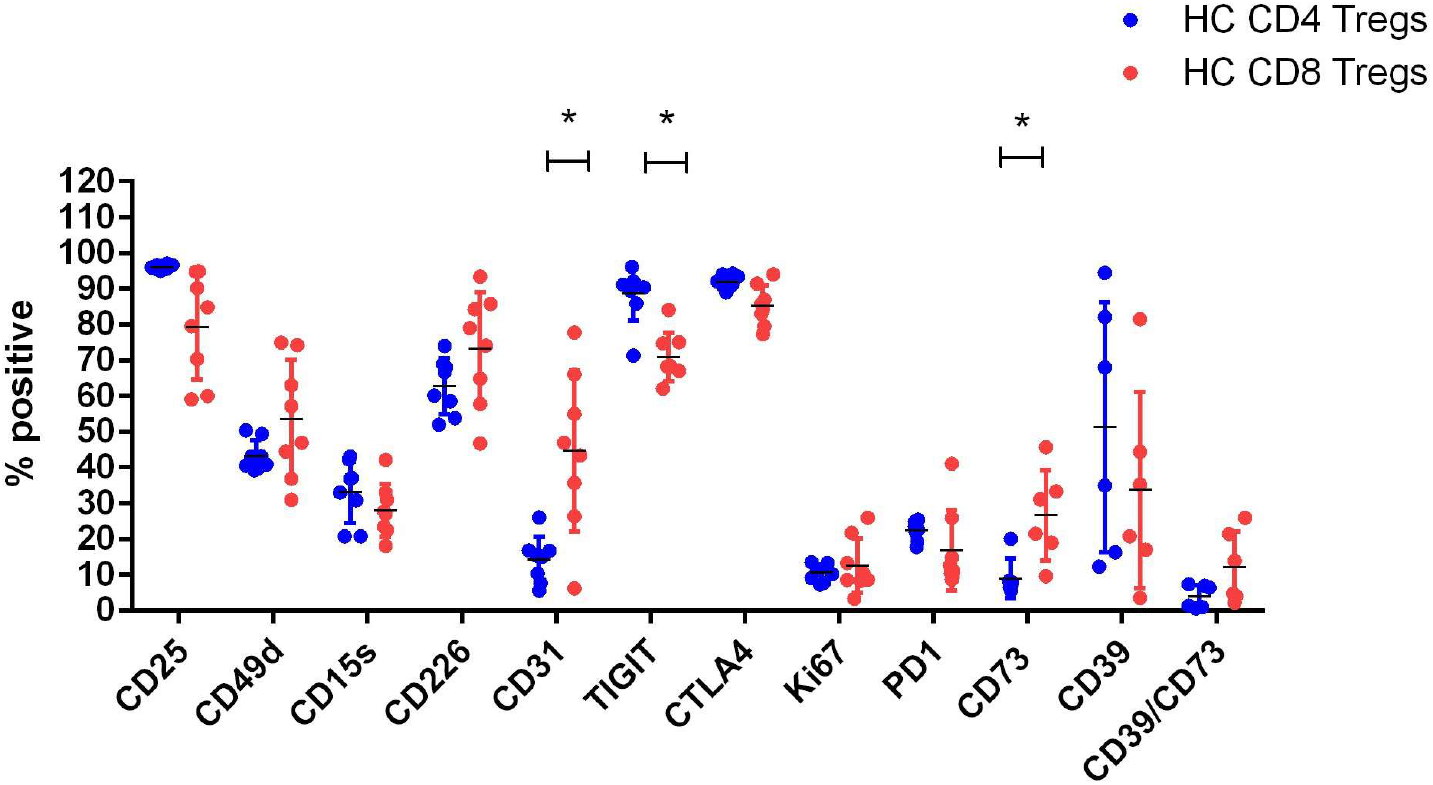
Expression of Treg markers on CD3+FOXP3+HELIOS+ CD4+ and CD8+ cells in human peripheral blood. Expression of Canonical Treg markers on CD3+CD127loFOXP3+HELIOS+ CD4 (blue) and CD8 (red) cells. Data for n = 8 healthy donors is shown (mean +/− SD). Statistical significance determined using multiple paired T-tests with the Holm-Sidak method for multiple testing correction.

**Supplementary Fig 3.**
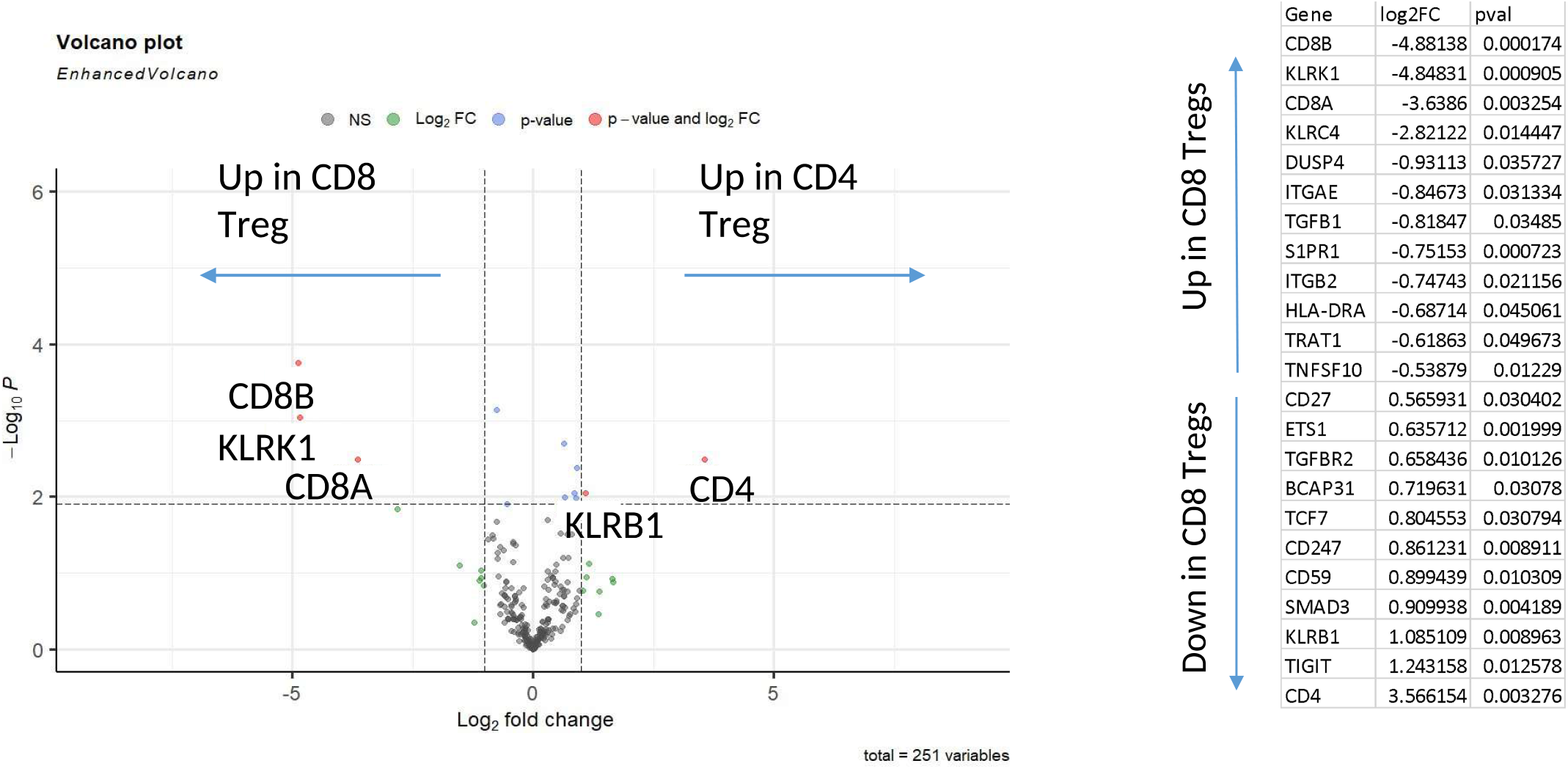
Nanostring gene expression data. Nanostring transcriptomic analysis comparing differential gene expression of surface sorted (CD3+CD25hiCD127lo) CD8+ and CD4+ Tregs from peripheral blood (human immunology panel_v2; n = 3 donors).

**Supplementary Fig 4.**
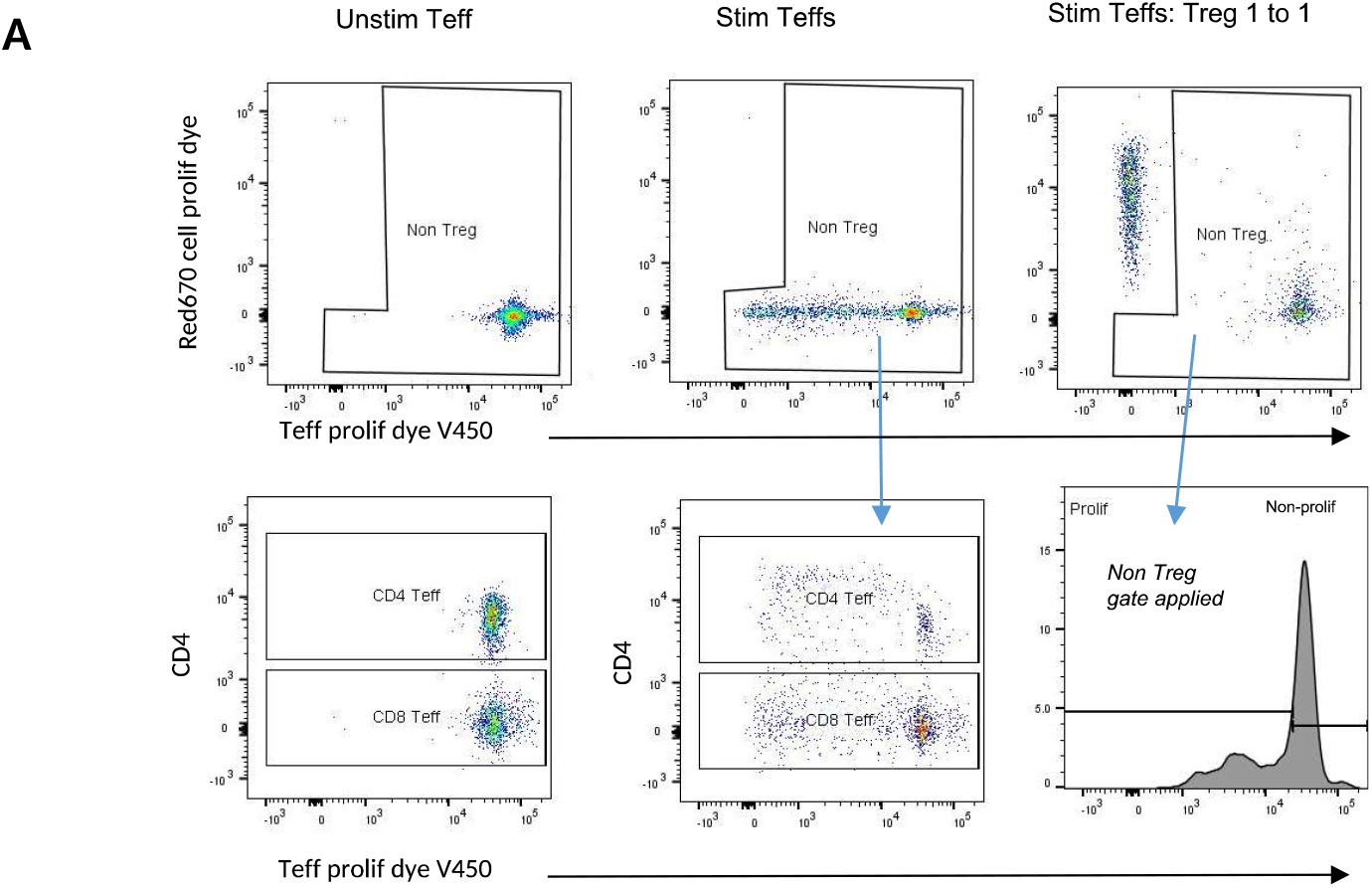
Suppression assay gating strategy. CD3+ Pan T cells labelled with V450 cell proliferation dye unstim (left plots) and stimulated with anti-CD3/28 beads (middle plots) in the absence and presence (top right) of R670 cell tracker labelled Tregs. R670+ Tregs are gated out and Non-Treg cells analysed for proliferation, setting regions for non proliferating and proliferating T effectors on histogram (bottom right).

**Supplementary Fig 5.**
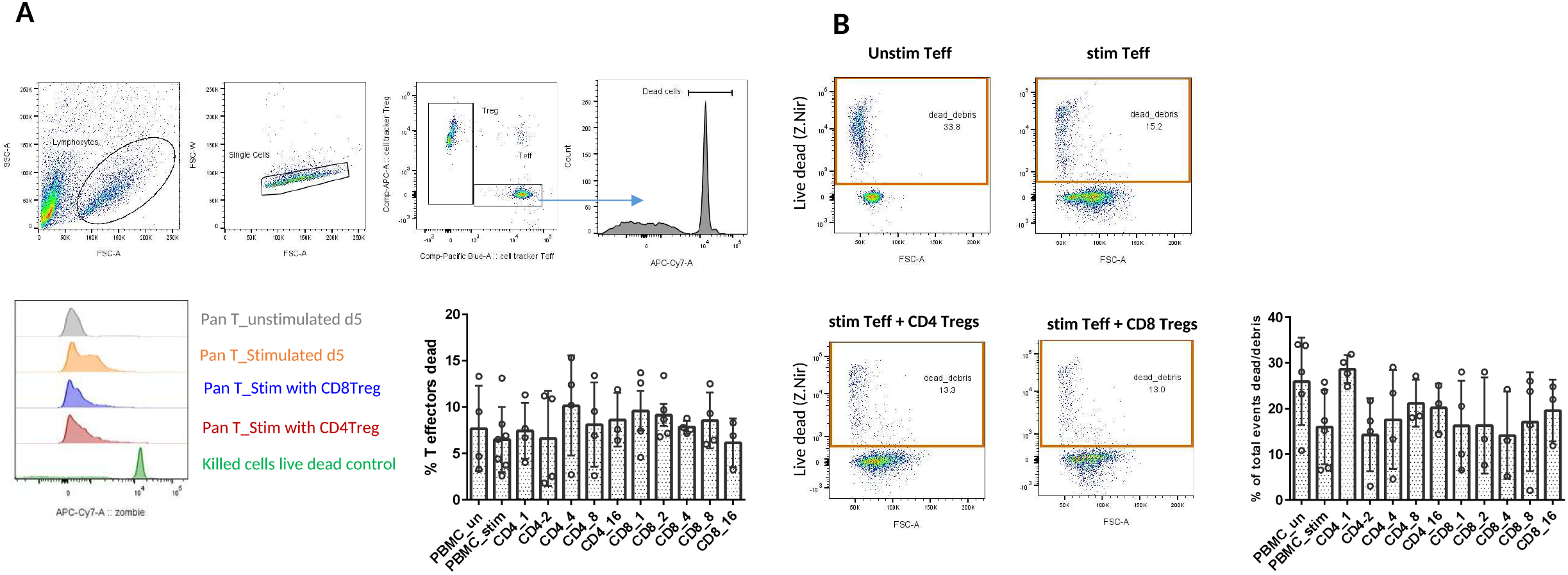
Cell death in suppression assay in presence of CD8+ Tregs. A: V450+ Pan T cell effectors were gated before live/dead exclusion and % dead cells gated for all conditions. The % cell death in Teffector population was not significantly different between those cultured in presence of CD4+Tregs or CD8+ Tregs suggesting limited cytotoxic function of CD8+ Tregs. B: Total dead cells and debris were gated for each condition and proportions compared. No statistical difference in cell death was detected in different assay conditions or between Teffectors cultured with CD8+ Tregs compared to Teffectors cultured with CD4+ Tregs (one way ANOVA with Tukey’s multiple comparison correction)

**Supplementary Fig 6.**
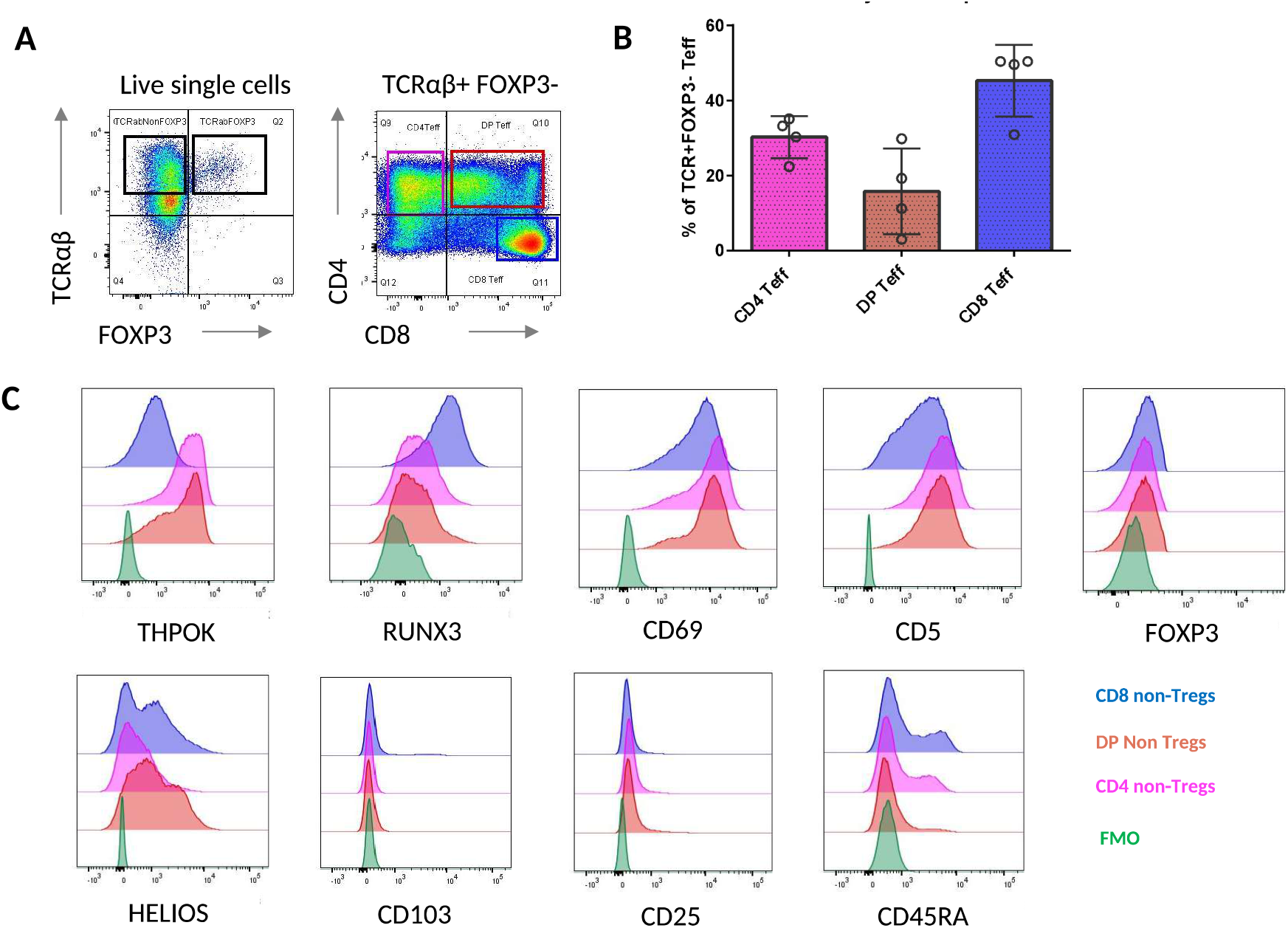
Effector T cells in Thymus. A: TCRαβ vs FOXP3 staining on live lymphocytes (left) and CD4 vs CD8 staining on gated TCRαβ+FOXP3-T effector cells in human thymus (representative plot) B: Percentage ofCD4+ SP (pink), CD4+CD8+ DP (orange) and CD8+SP (blue) FOXP3-cells in human thymus (n =3 donors) C: Flow staining of canonical Treg and mature thymocyte markers and transcription factors THPOK and RUNX3 CD4+ (pink), CD8+ (blue) and DP+ (orange) FOXP3-thymocytes compared with unstained or FMO controls (green). Representative example of n = 3 donors.

**Supplementary Fig 7.**
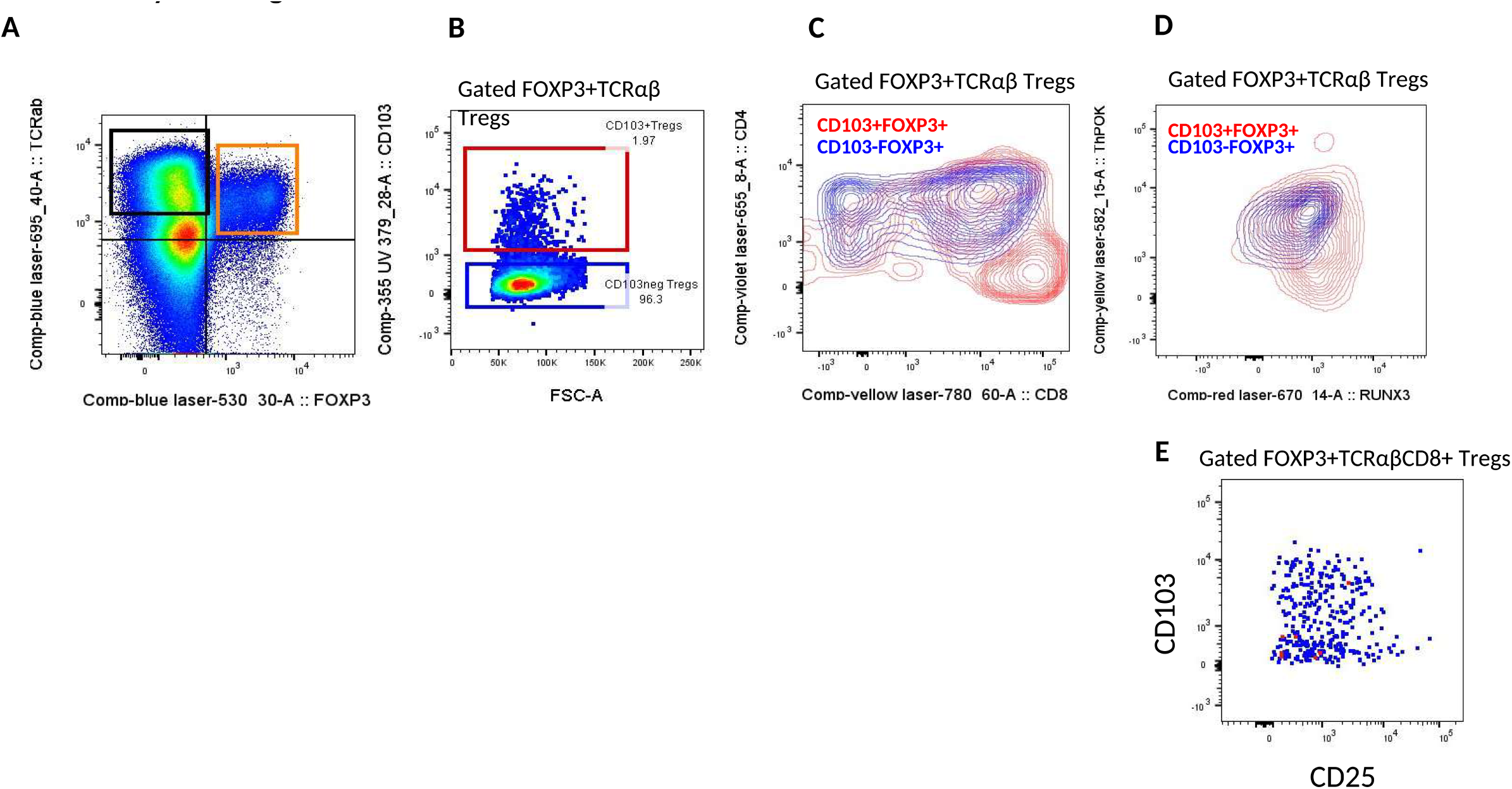
CD103 in Thymus Tregs A. A: TCRαβ vs FOXP3 staining on live lymphocytes (left), representative plot of n = 3 human thymi B: Gating of CD103+ (red) and CD103-(blue) Tregs within the FOXP3+ gate, representative plot ofn = 3 human thymi C: CD4 vs. CD8 staining within CD103+FOXP3+ Tregs (red) and CD103-FOXP3+ Tregs (blue) D: THPOK vs RUNX3 staining within CD103+FOXP3+ Tregs (red) and CD103-FOXP3+ Tregs (blue) E: CD103 vs CD25 on human CD3+CD8+FOXP3+TCRab+ Tregs, representative plot

**Supplementary Fig 8.**
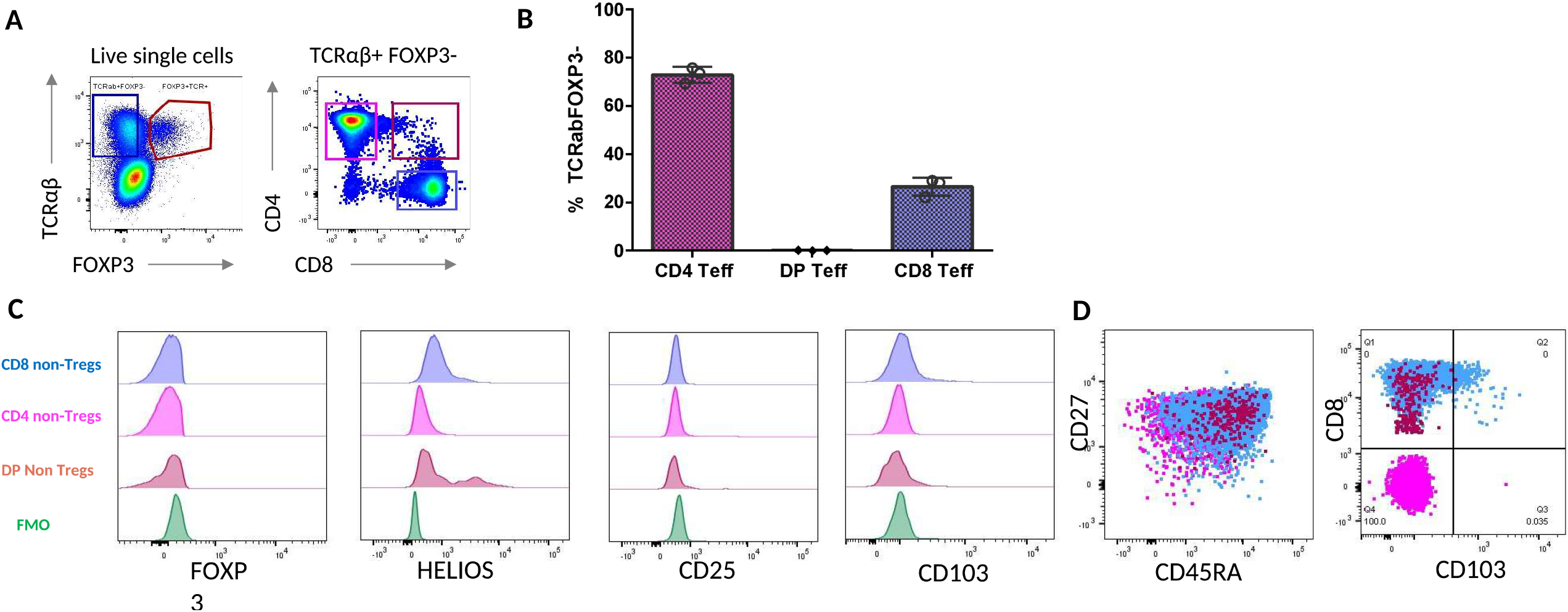
Effector T cells in cord bloods. A: TCRαβ vs FOXP3 staining on live lymphocytes (left) and CD4 vs CD8 staining on gated TCRαβ+FOXP3-T cells in human cord blood (representative plot) B: Percentage of CD4+ SP (pink), CD4+CD8+ DP (red) and CD8+SP (blue) FOXP3-cells in human cord blood (n = 3 donors) C: :Flow cytometric phenotyping of cord blood TCRαβ+FOXP3-Tregs comparing CD4+ (pink), CD8+ (blue) and DP+ (red) with unstained or FMO controls (green). Overlaid histograms representative example of n = 3 donors. D: Dotplot overlays showing expression ofCD27 vs CD45RA and CD8 vs CD103 in CD4+FOXP3-TCRαβ+ T effector (Teff) cells (pink), CD8+FOXP3-TCRαβ+ Teffs (blue) and CD4+CD8+ (DP) +FOXP3-TCRαβ+Teffs (red), representative example ofn = 3 donors.

**Supplementary Fig 9.**
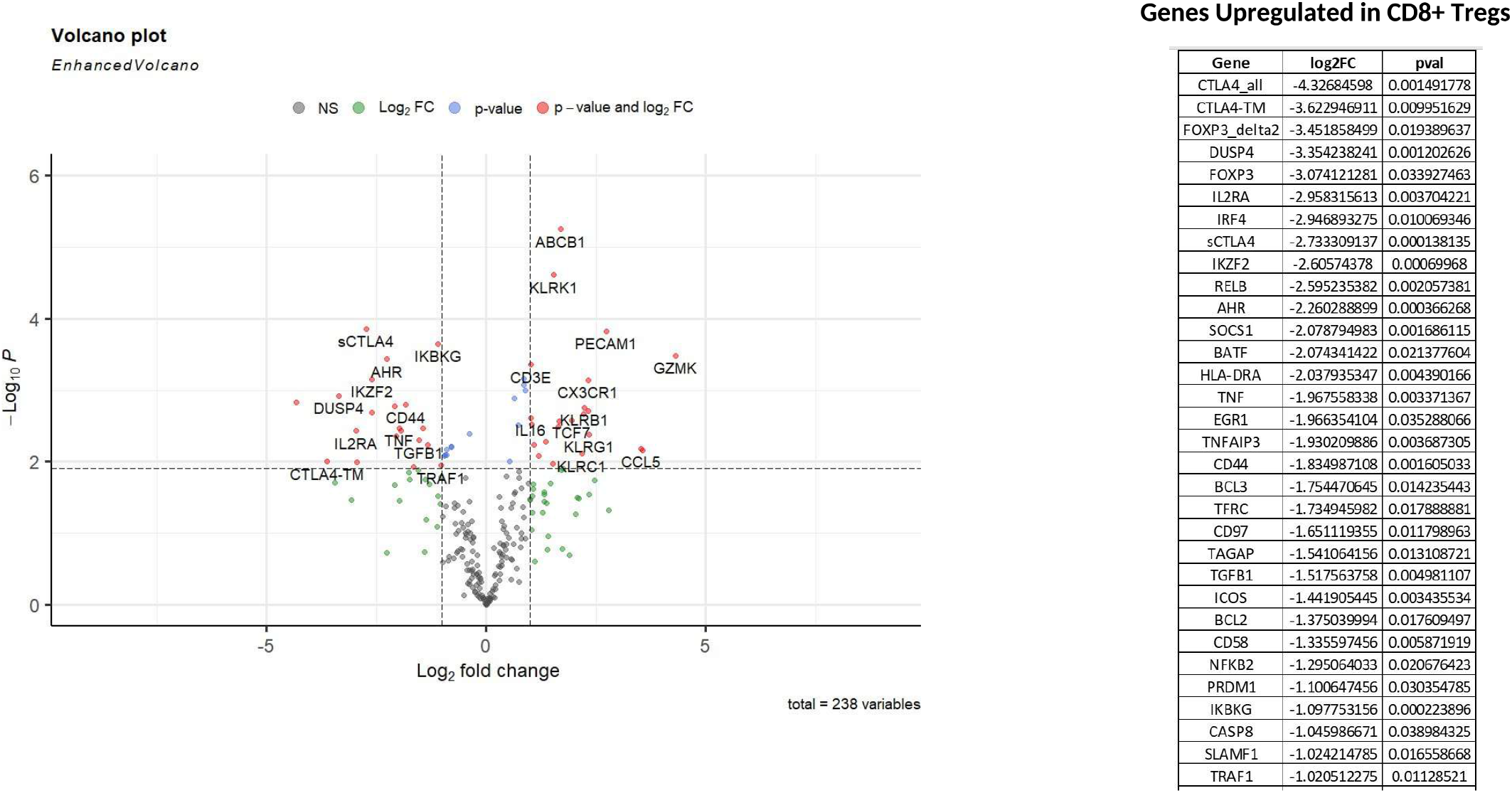
Differential Expression of Nanostring genes between CD8 Tregs and CD8 T effectors. DE of genes between CD3+CD25hiCD127lo CD8+ Tregs and CD3+CD25-CD8 T effectors (left) sorted from human peripheral blood, by Nanostring and top 30 upregulated genes in CD8+ Tregs (right)

**Supplementary Fig 10.**
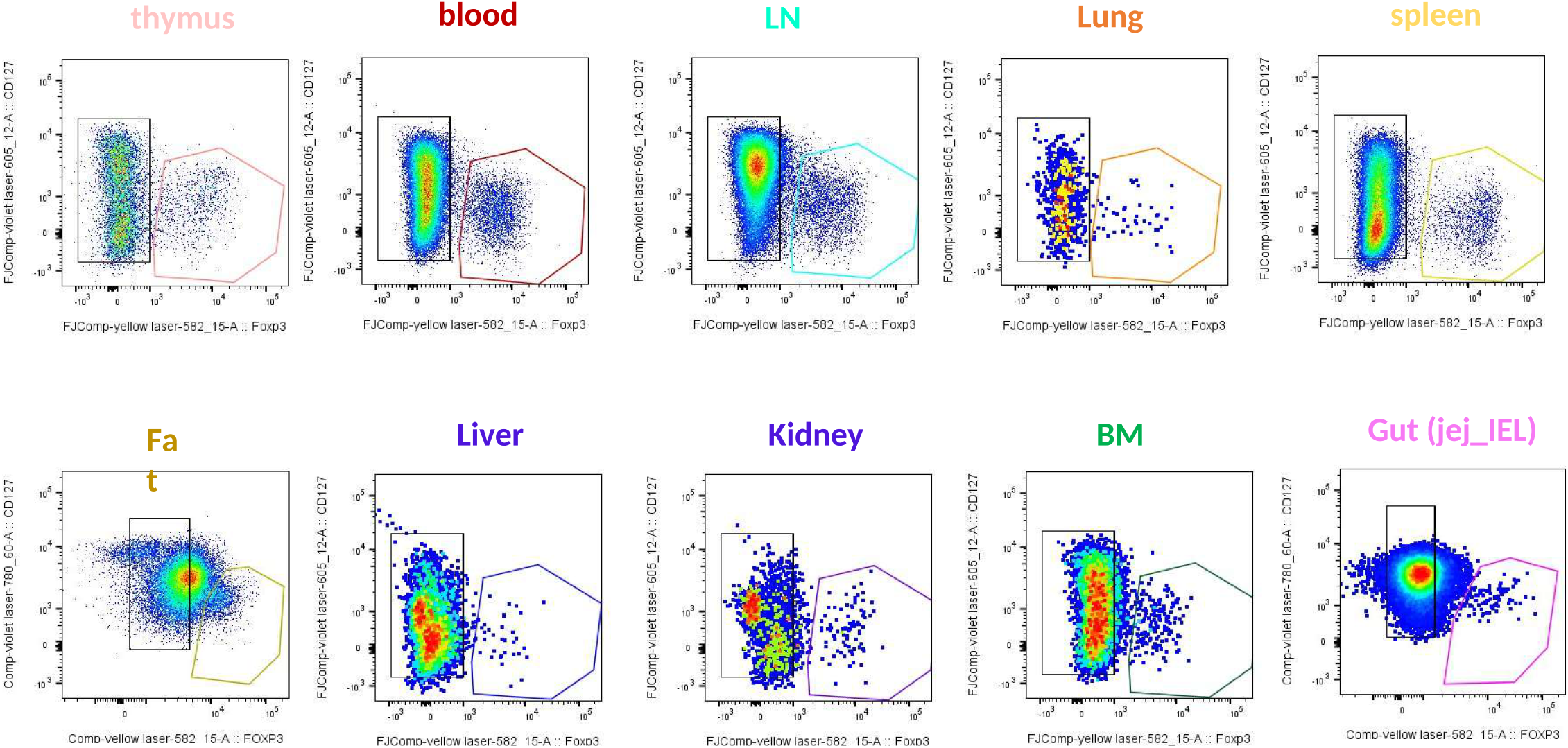
Gating strategy example of 127 vs FOXP3 by tissue.

**Supplementary Fig 11.**
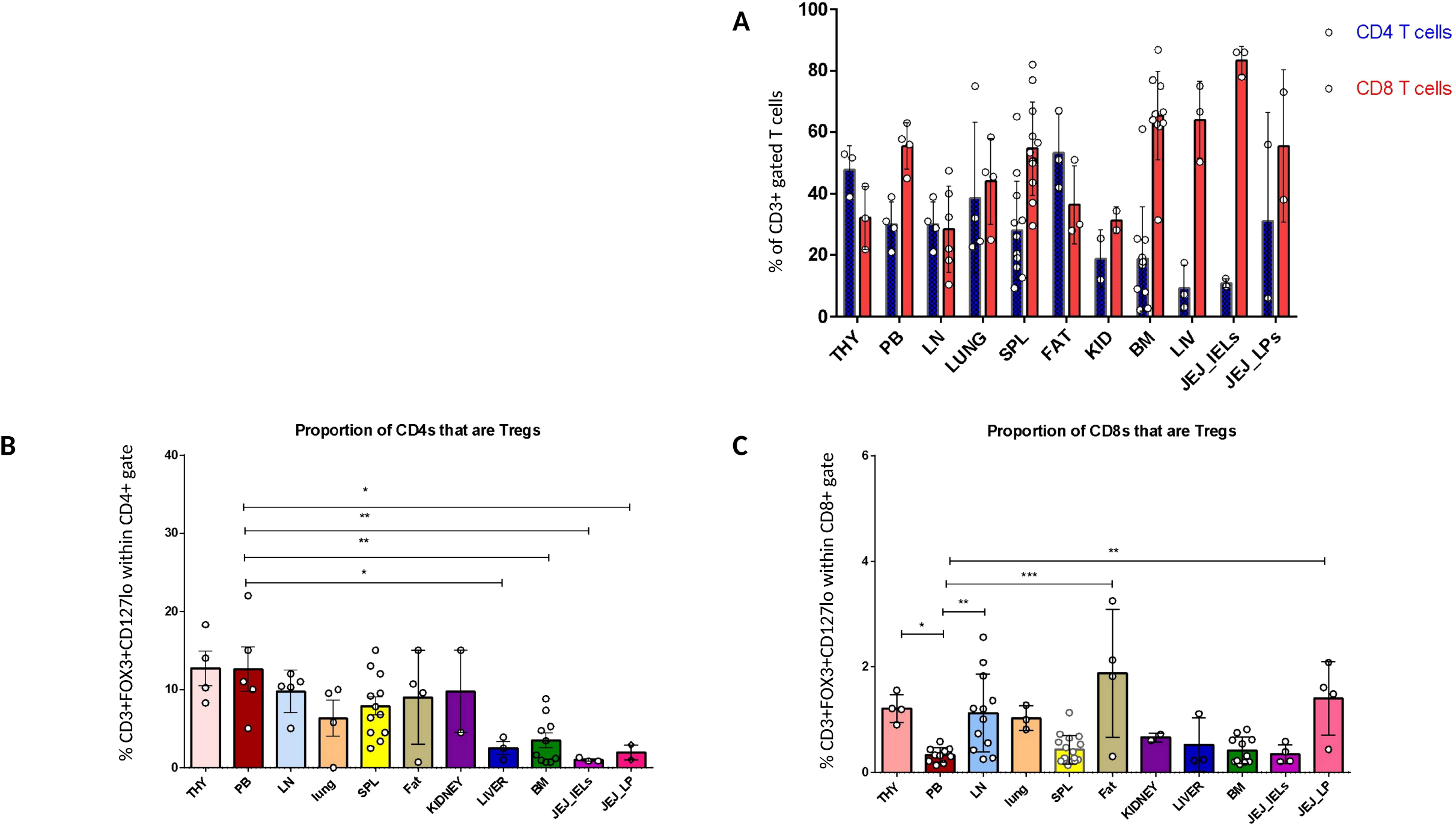
Proportions of CD4 and CD8 Teff and Tregs across tissues. A: Summary of % CD4+ T cells (blue bars) and CD8+ T cells (red bars) within live CD3+ gated human T cells across multiple tissues B: % of CD3+CD4+ T cells that are CD4+CD1271oFOXP3+ across multiple human tissues. One way ANOVA comparisons to PB with Dunnett’s multiple testing correction. C: % of CD3+CD8+ T cells that are CD8+CD1271oFOXP3+ across multiple human tissues. One way ANOVA comparisons to PB with Dunnett’s multiple testing correction.

**Supplementary Fig 12.**
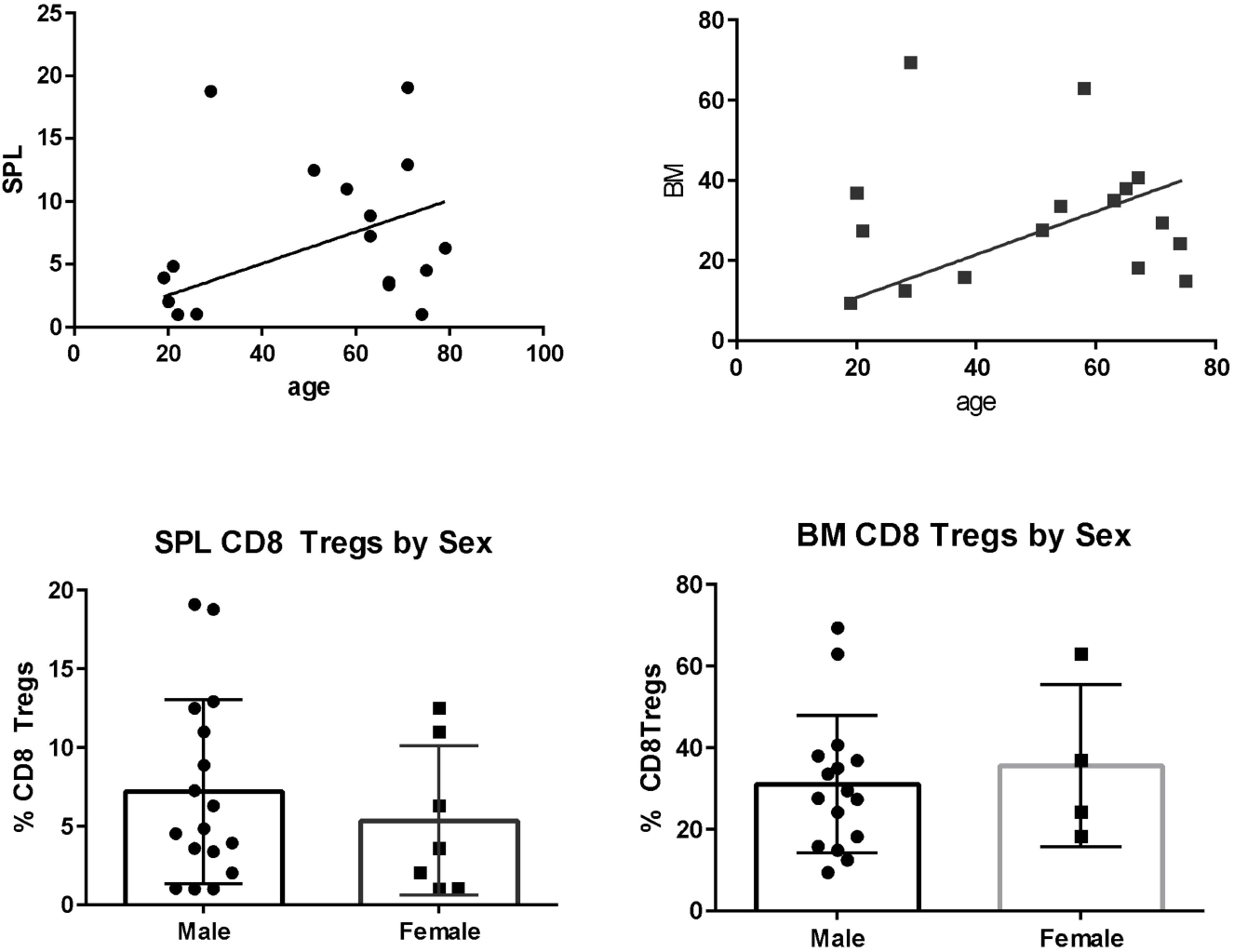
% of FOXP3+CD127lo Tregs that are CD8+ across age and gender. A: Percentage of CD8+ Tregs within total CD3+FOXP3+HELIOS+ Treg gate in spleen (SPL) samples, plotted against age B: Percentage of CD8+ Tregs within total CD3+FOXP3+HELIOS+ Treg gate in bone marrow (BM) samples, plotted against age. Trend against ageing did not reach statistically significant correlation C: Percentage of CD8+ Tregs within total CD3+FOXP3+HELIOS+ Treg gate in spleen (SPL) samples in male and female donors in spleen. D: Percentage of CD8+ Tregs within total CD3+FOXP3+HELIOS+ Treg gate in spleen (SPL) samples in male and female donors in BM.

**Supplementary Fig 13:**
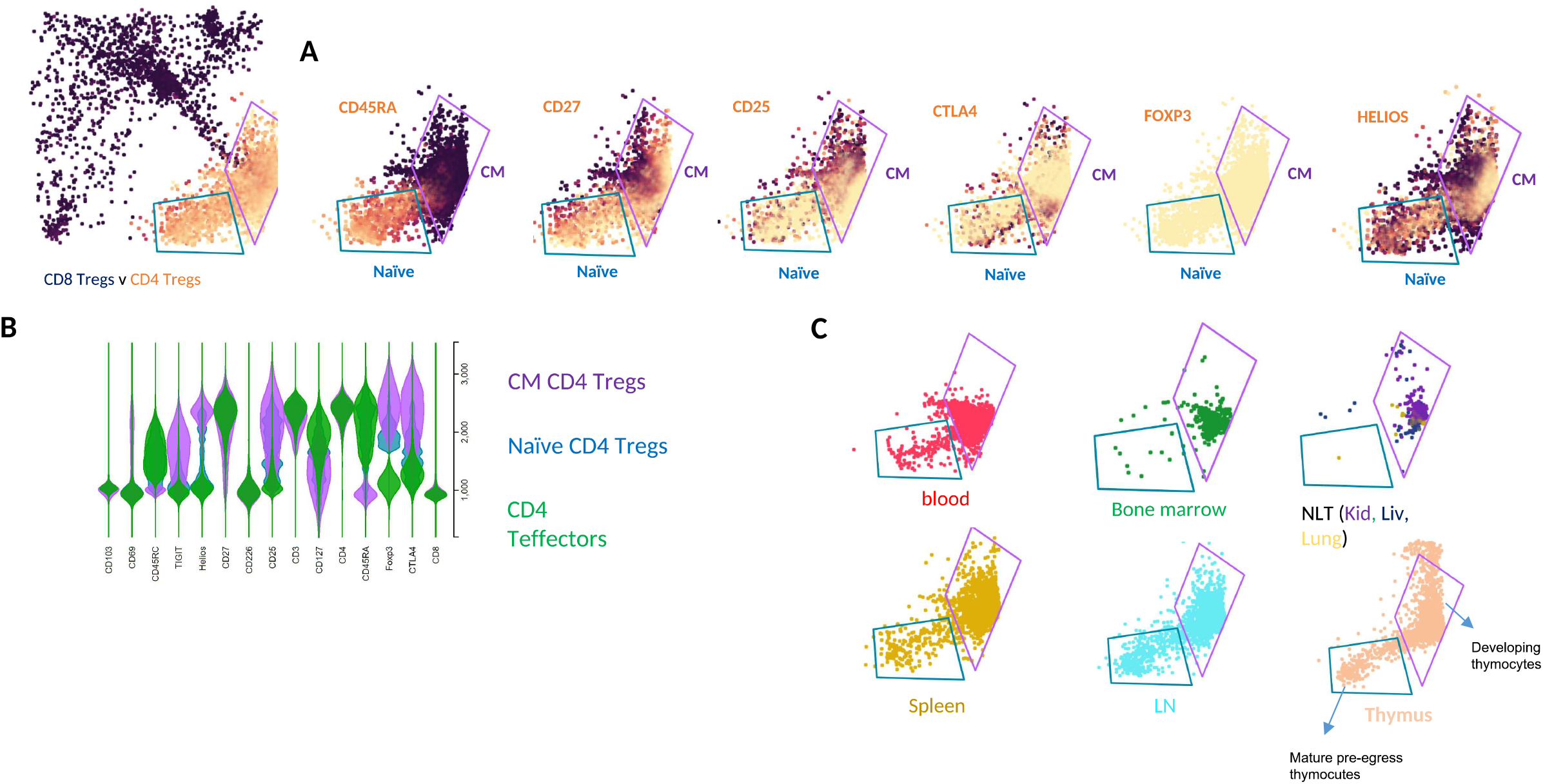
CD4 Treg_FlowAtlas Images. A: CD8 and CD4 Tregs flow cytometry data displayed in FlowAtlas and expression of Treg markers in CD4+ Treg subset. B: summary data of expression levels (all tissues) for CD4+FOXP3+CD127lo Tregs compared with CD4+ Teffector cells. C: distribution of naïve and CM CD4+ Tregs across different tissues. Note: as expected in thymus, the majority of single positive developing thymocytes lack CD45RA expression and therefore fall into the “CM” region whilst mature thymocytes closest to egress gain CD45RA and fall into the “Naïve” region.

**Supplementary Fig 14.**
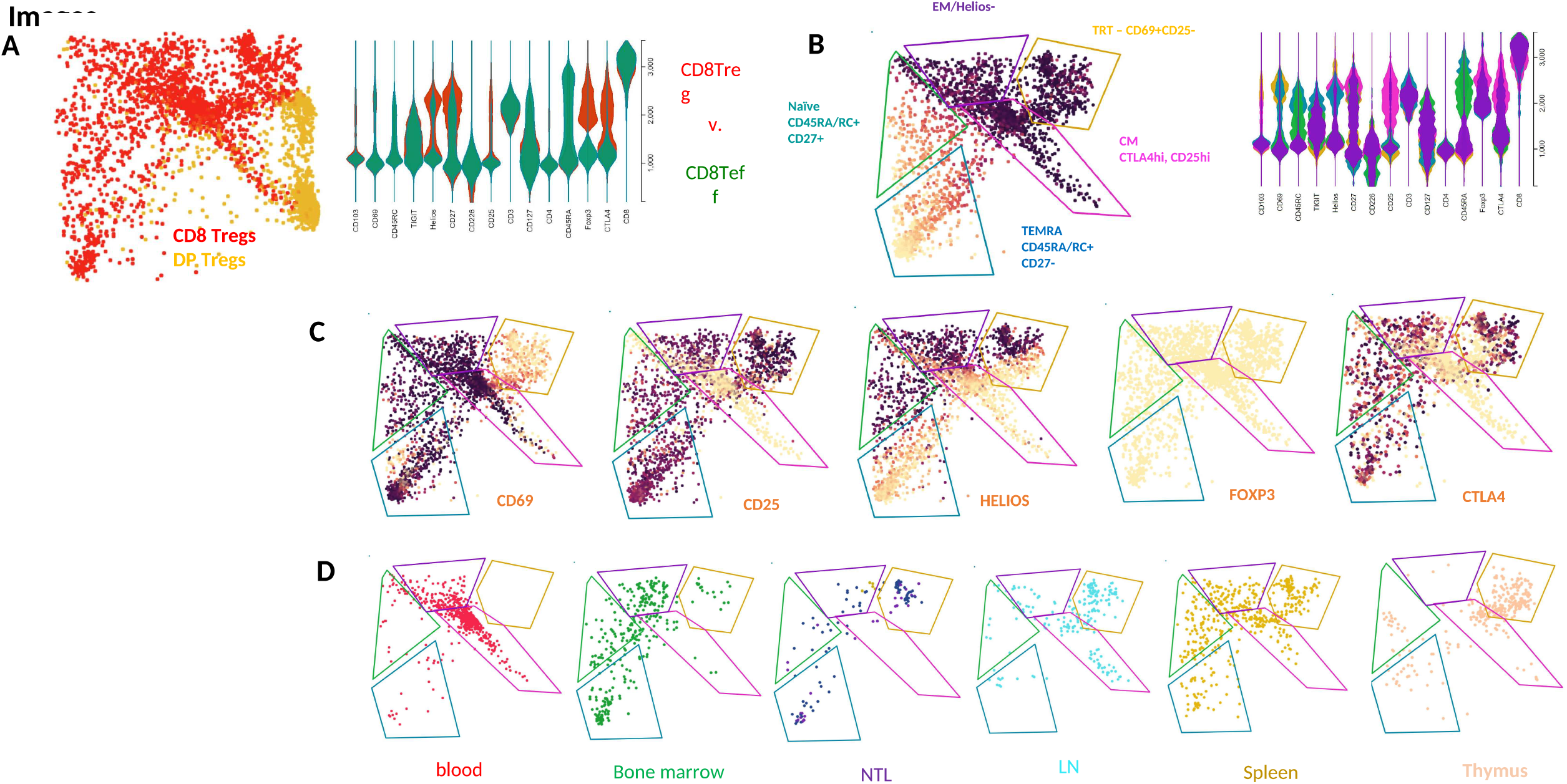
CDS Treg_FlowAtlas Images. A: CD8 and DP Tregs flow cytometry data displayed in FlowAtlas and expression of Treg markers in CDS+FOXP3+CD127lo Treg subset compared with CDS+FOXP3-Teffectors. B: CD8+FOXP3+CD127lo Tregs can be divided into 5 distinct subpopulations. C: Expression of Treg markers within subsets for CD8+ Tregs D: Distribution of CD8+ Treg subsets across tissues

**Supplementary Fig 15.**
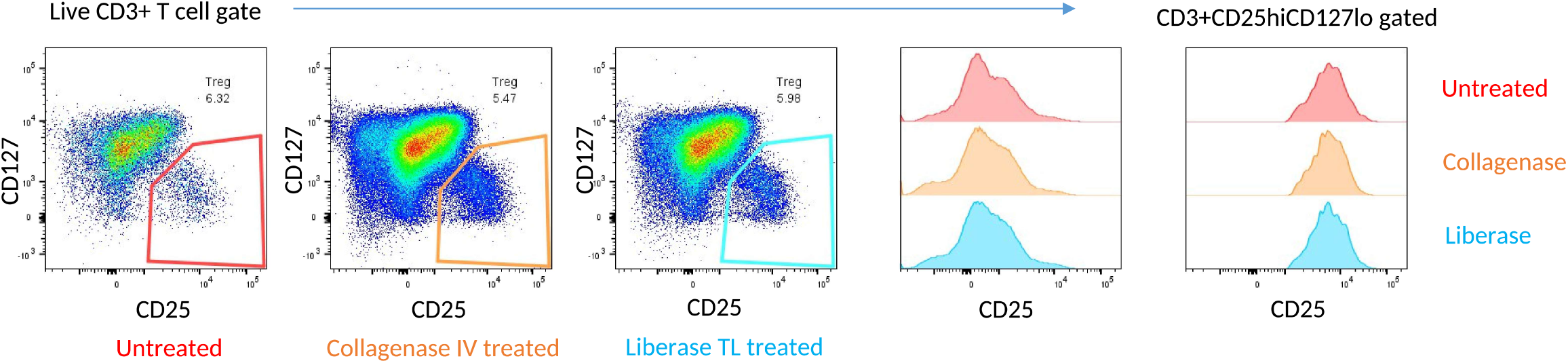
CD25 post enzymatic treatment. Human PBMC untreated (red) or treated with Collagenase IV (orange) or Liberase TL (blue) for 30 minutes at 37°C were washed and stained for Treg markers CD25 and CD127. Representative example gated on Live CD3+ T cells is shown. Histogram overlays show CD25 on live CD3+ T cells (left histogram) and on CD25hiCD127lo gated cells (right histogram).

**Supplementary Fig 16.**
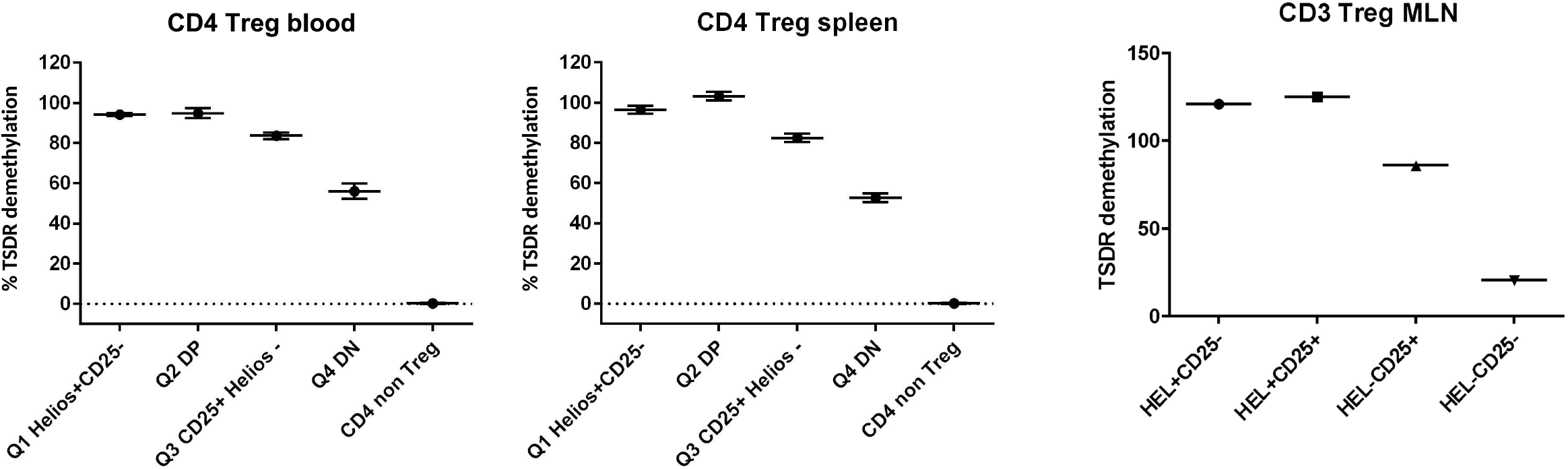
CD4 Treg_FOXP3 TSDR demethylation. TSDR demethylation patterns in FOXP3+CD127lo Tregs on the basis of co-expression of HELIOS and/or surface CD25. Sorted CD4+FOXP3+CD127lo Tregs from human peripheral blood (left plot) and human spleen (middle plot) and in CD3+FOXP3+CD127lo Tregs from human mesenteric Lymph node (MLN, right plot) are shown.

**Supplementary Fig 17.**
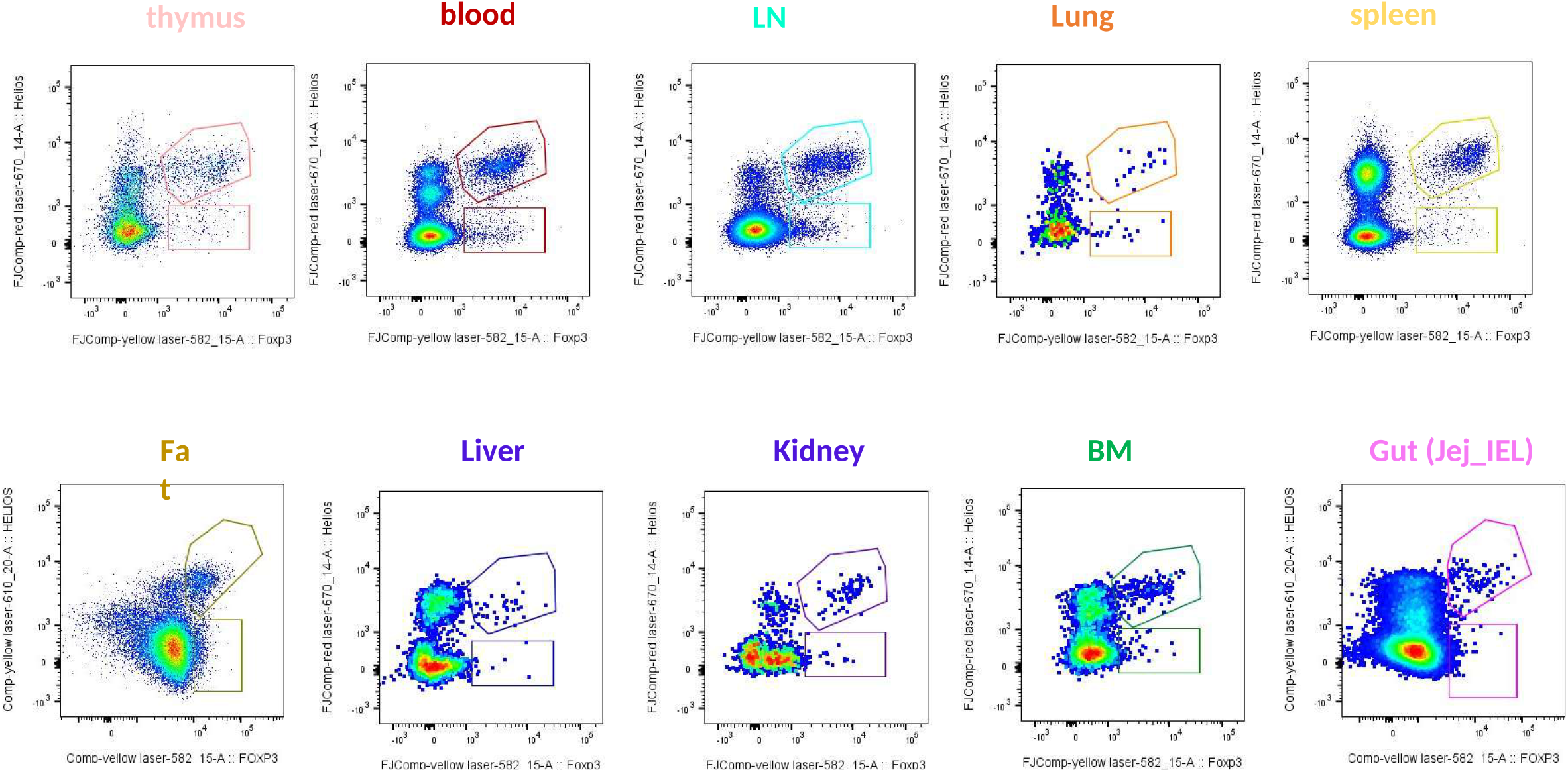
Gating strategy example of 127 vs FOXP3 by tissue.

**Supplementary Fig 18.**
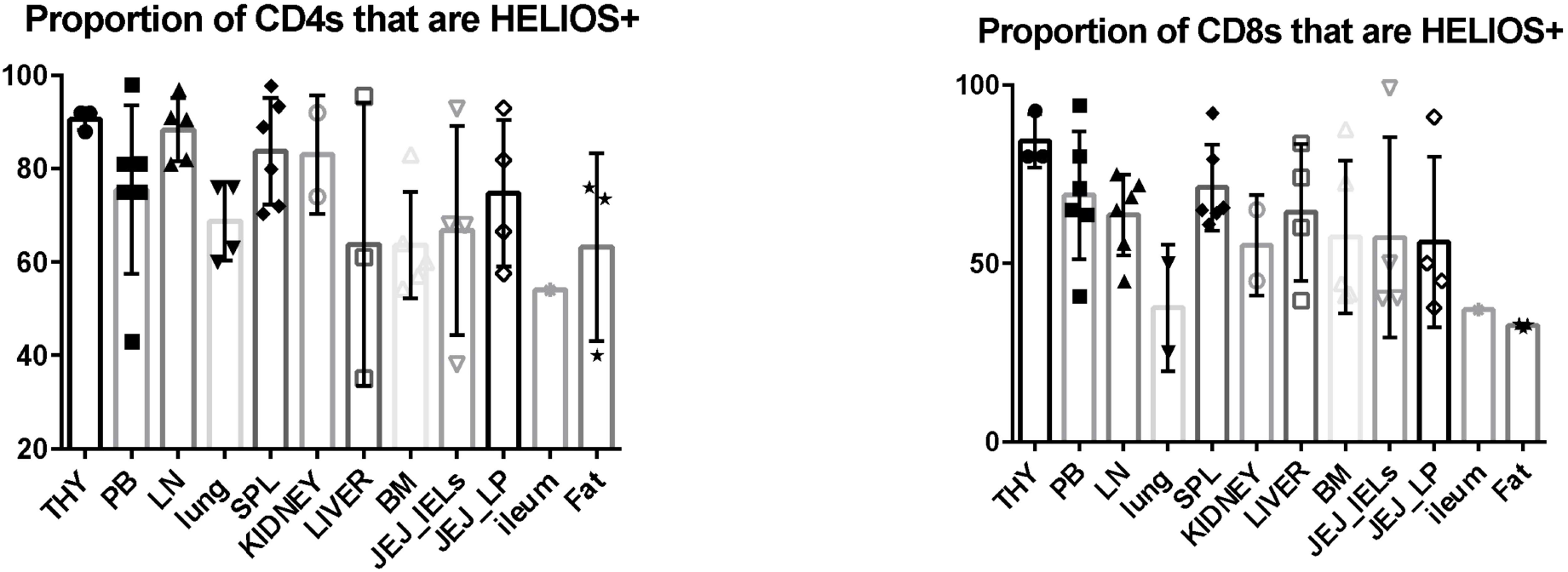
Proportion of Helios+ and Helios− CD4 and CD8 Tregs cells across human tissues.

**Supplementary Figure 19:**
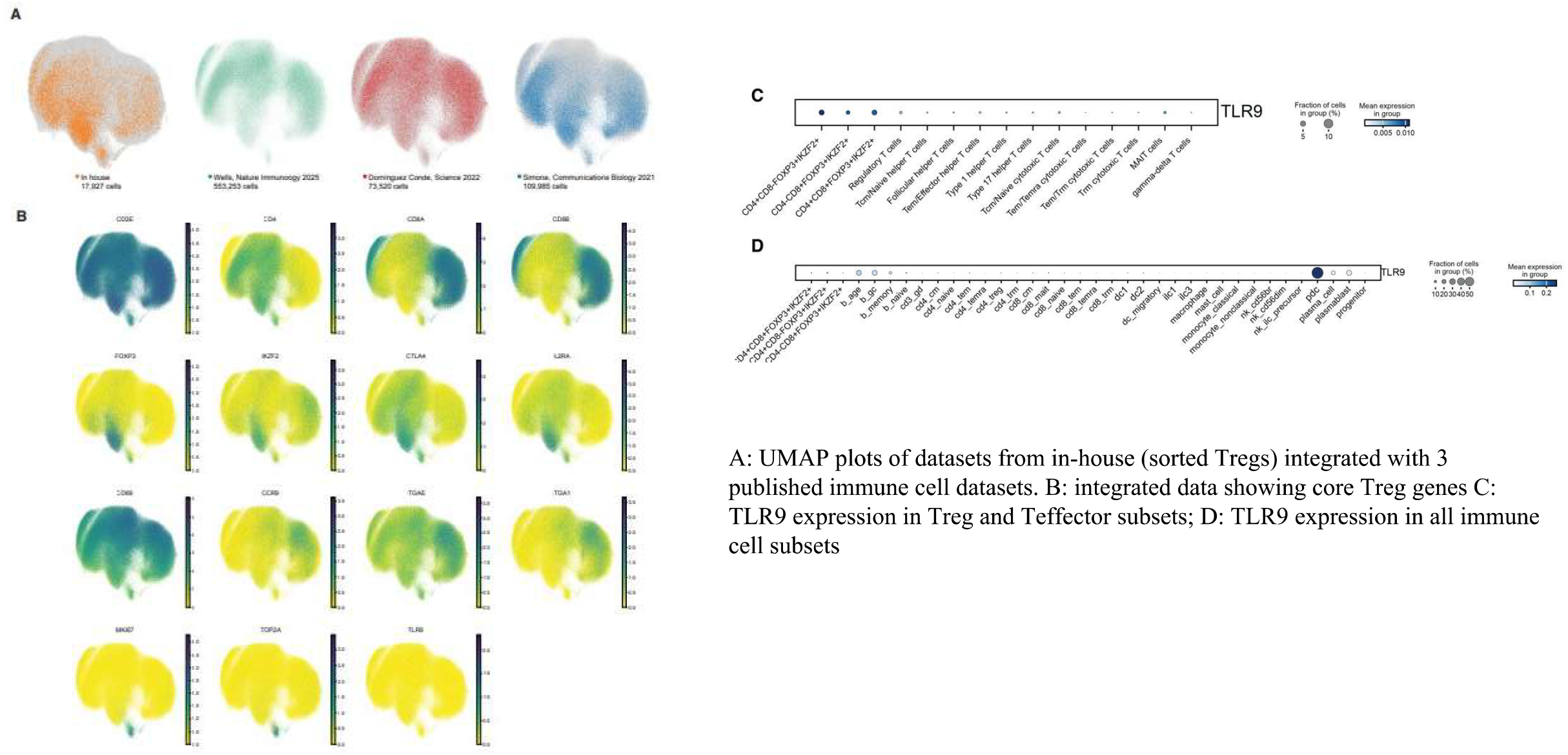
Additional RNAseq single cell data. A: UMAP plots of datasets from in-house (sorted Tregs) integrated with 3 published immune cell datasets. B: integrated data showing core Treg genes C: TLR9 expression in Treg and Teffector subsets; D: TLR9 expression in all immune cell subsets

**Supplementary Figure 20:**
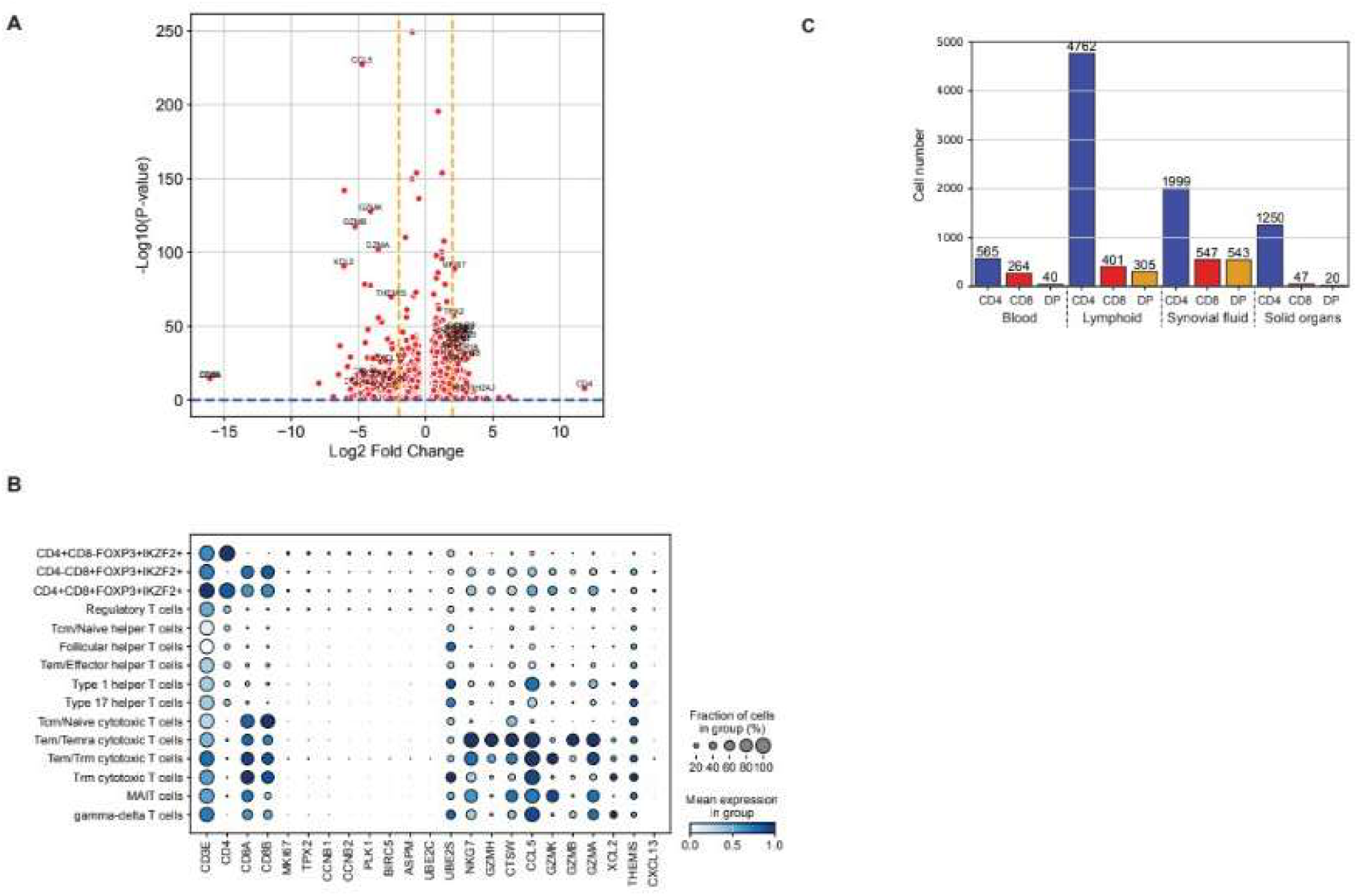
single cell RNA sequencing data for Tregs in tissue datsets. A: volcano plot summarising DE of CD4 and CD8+ Tregs; C: genes upregulated in CD4 vs CD8 Tregs B: gene expression by annotated cell type of integrated data C: Bar chart showing cell numbers across integrated datasets;

**Supplementary Fig 21.**
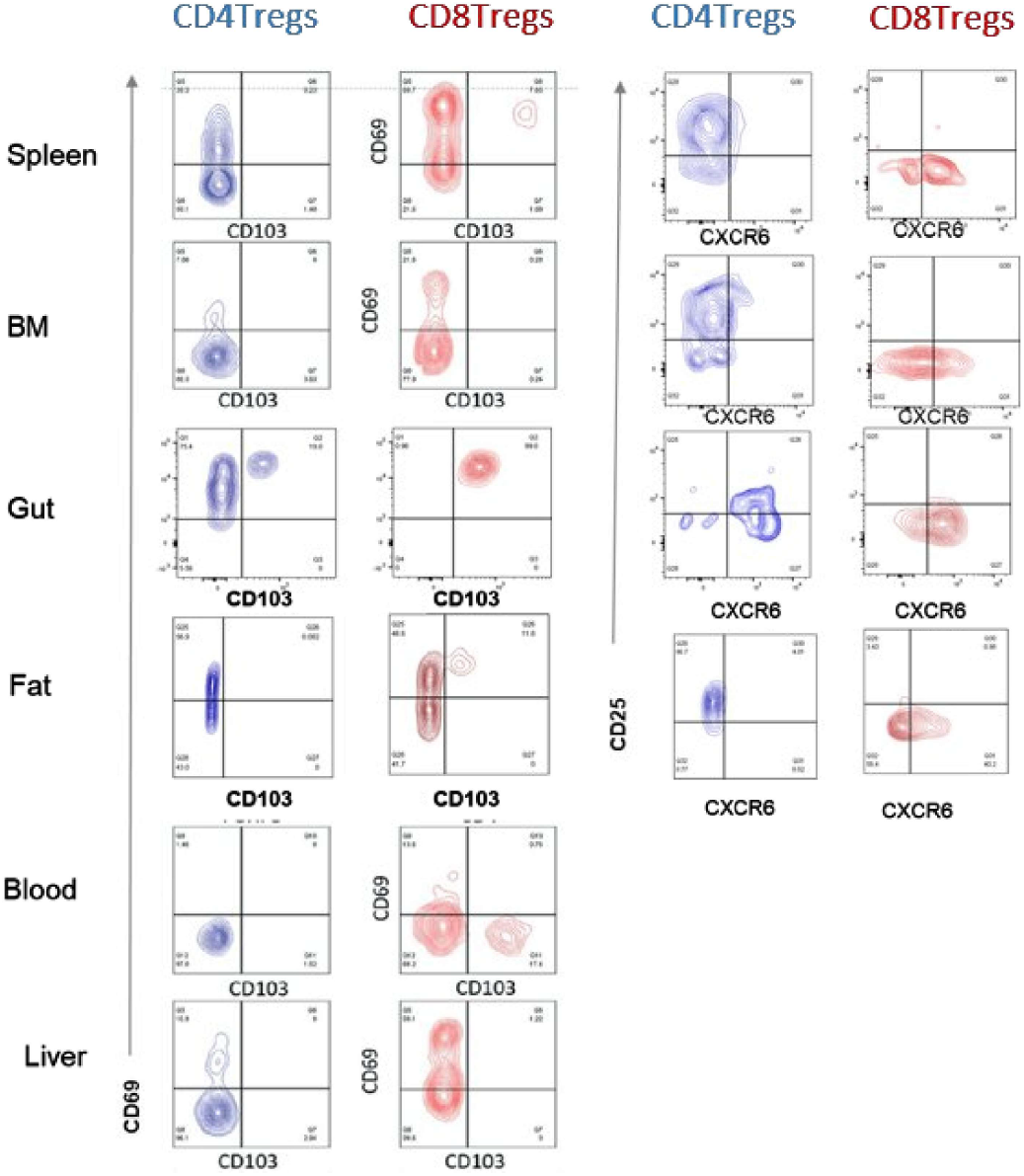
Tissue resident Treg phenotype across human tissues. CD69 vs CD103 expression (left) and CD25 vs CXCR6 expression (right) in representative examples of each human tissue. Gated CD3+HELIOS+FOXP3+CD4+ Tregs (red) and CD3+HELIOS+FOXP3+CD8+ Tregs (blue) are shown

**Supplementary Fig 22.**
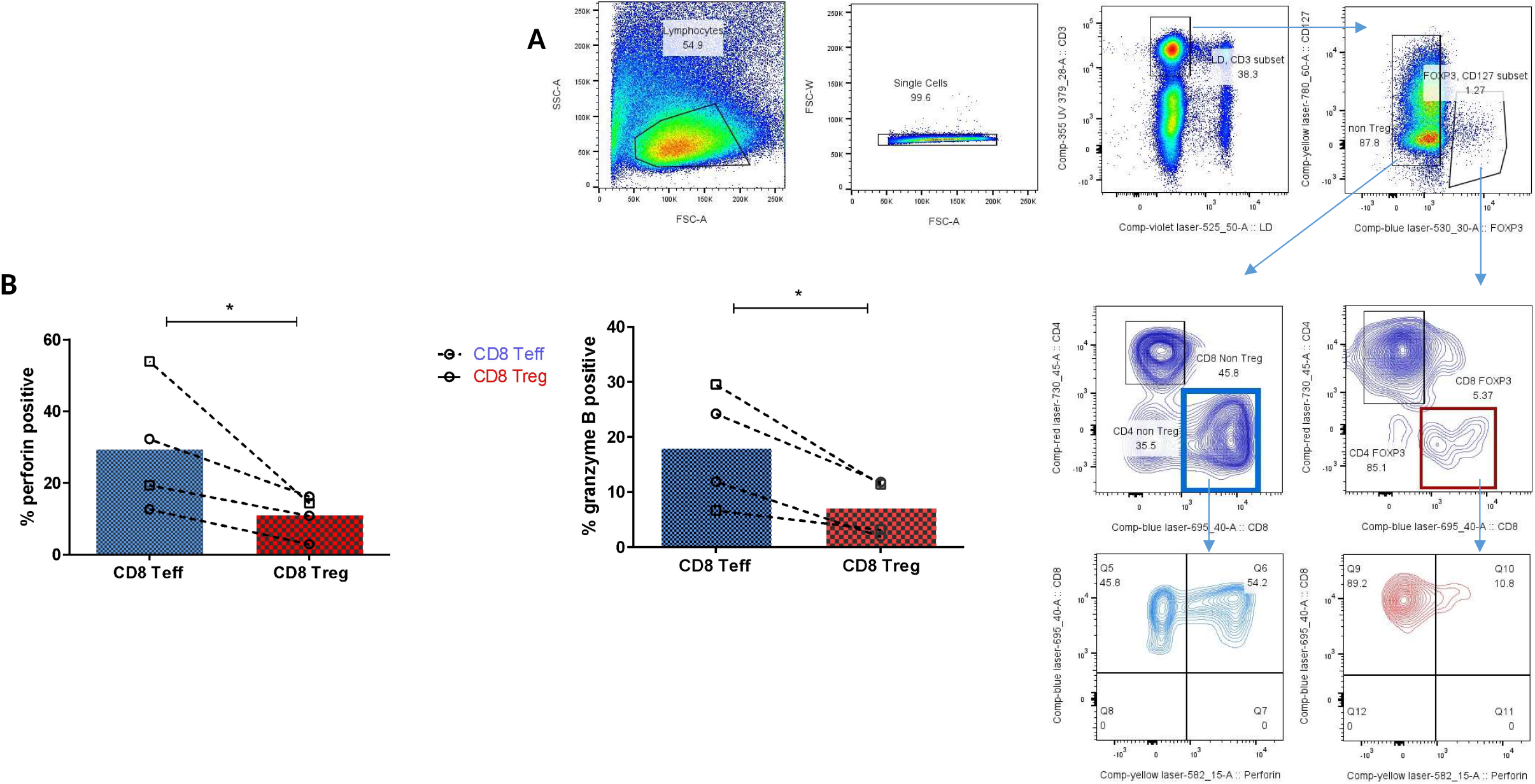
Perforin and granzyme staining In CD8+ Teffectors and CD8+FOXP3+ Tregs. A: representative gating strategy in human spleen of CD8+FOXP3+ Tregs (red) and CD8+FOXP3-T effectors (blue). Perforin and granzyme positivity was measured in each group. B: summary data ofperforin and granzyme positivity in CD8+ T effectors and CD8+ Tregs in 4 donors (2 PBMC - open circles, 2 Spleen MNCs - open squares)

**Supplementary Fig 23.**
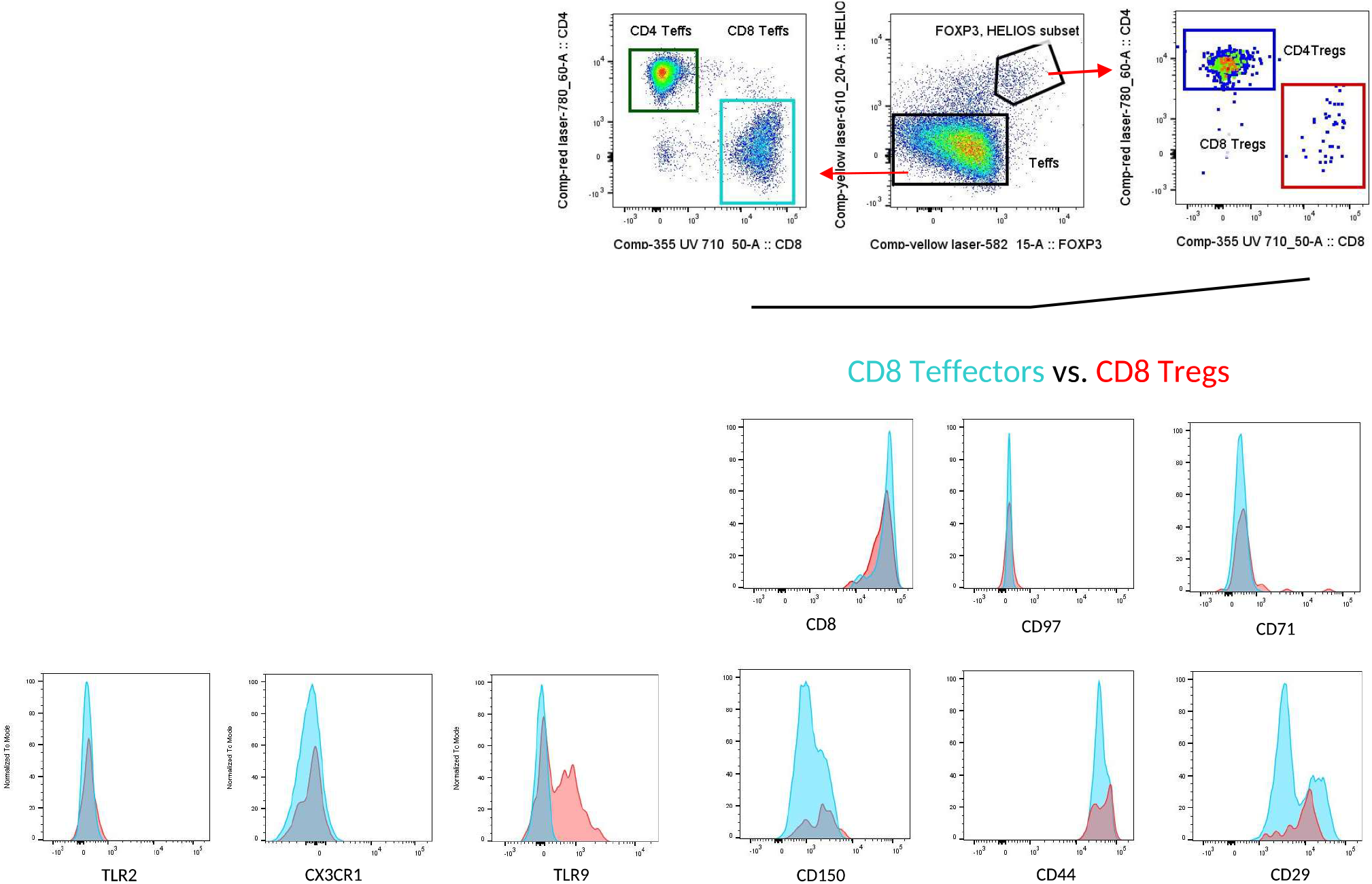
Screening for markers from Nanostring panel on CD8 Tregs vs. effectors. Histogram overlays of expression levels of a number of markers in CD8+ Tregs (red) vs CD8+ Teffector cells (turquoise).

**Supplementary Fig 24.**
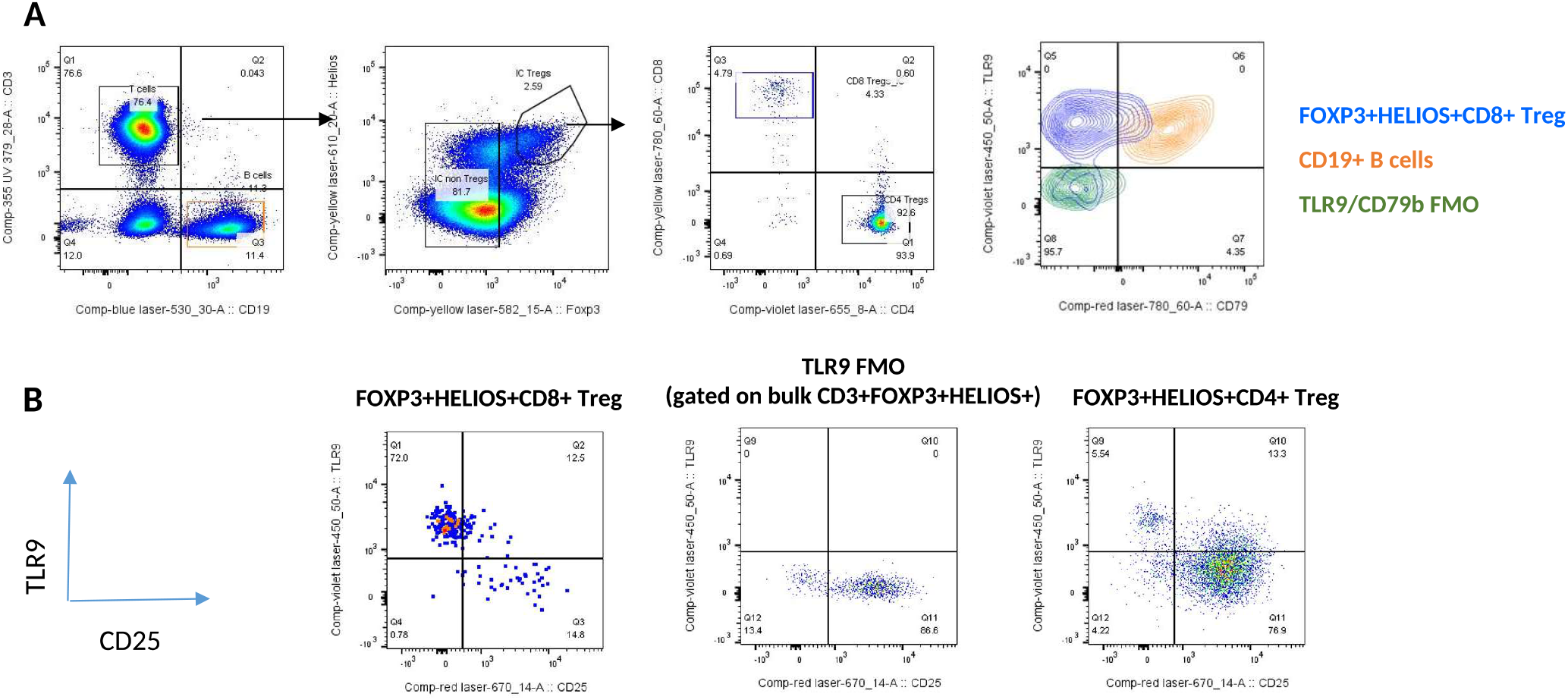
TLR9 staining on human Spleen CD8+ Tregs. A: TLR9 vs CD79B expression in CD19+ B cells (orange) and CD3+FOXP3+HELIOS+CD8+ Tregs (blue) (contour overlay with FMO for TLR9 shown in green). B: TLR9 vs CD25 expression on gated CD3+FOXP3+HELIOS+ CD8+ Tregs (left plot) and CD4+ Tregs (right plot). Data representative of n = 3 human spleen MNC samples. FMO staining for TLR9 is shown in middle plot.

**Supplementary Fig 25.**
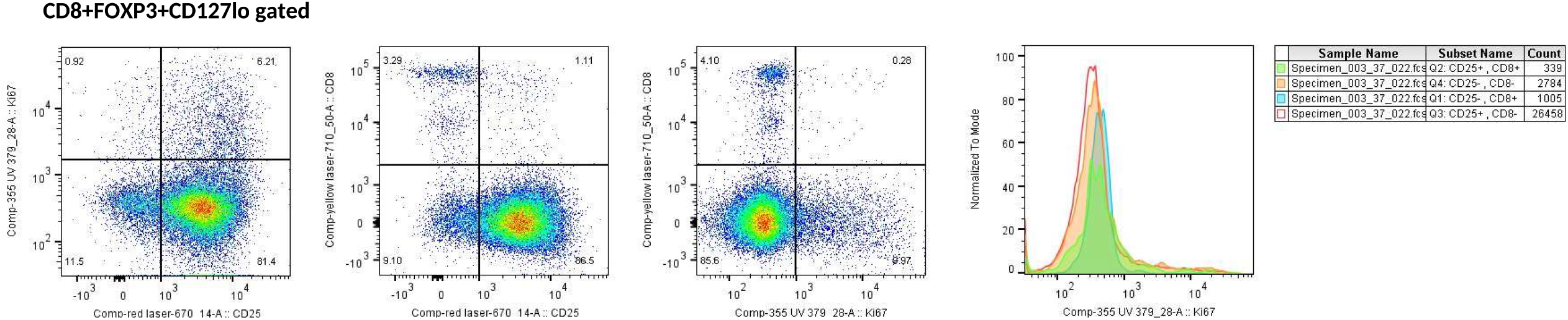
Surface CD25 vs Ki67 staining in CD4 vs CD8 Tregs (representative example) Human CD3+FOXP3+CD127lo gated Tregs in PBMC showing KI67 vs CD25 (left), CD8 vs CD25 (middle) and CD8 vs Ki67 (right). All surface CD25 negative cells in the Treg gate were low for Ki67 and CD8+CD25-cells did not express Ki67 so were not in cell cycle.

**Supplementary Fig 26:**
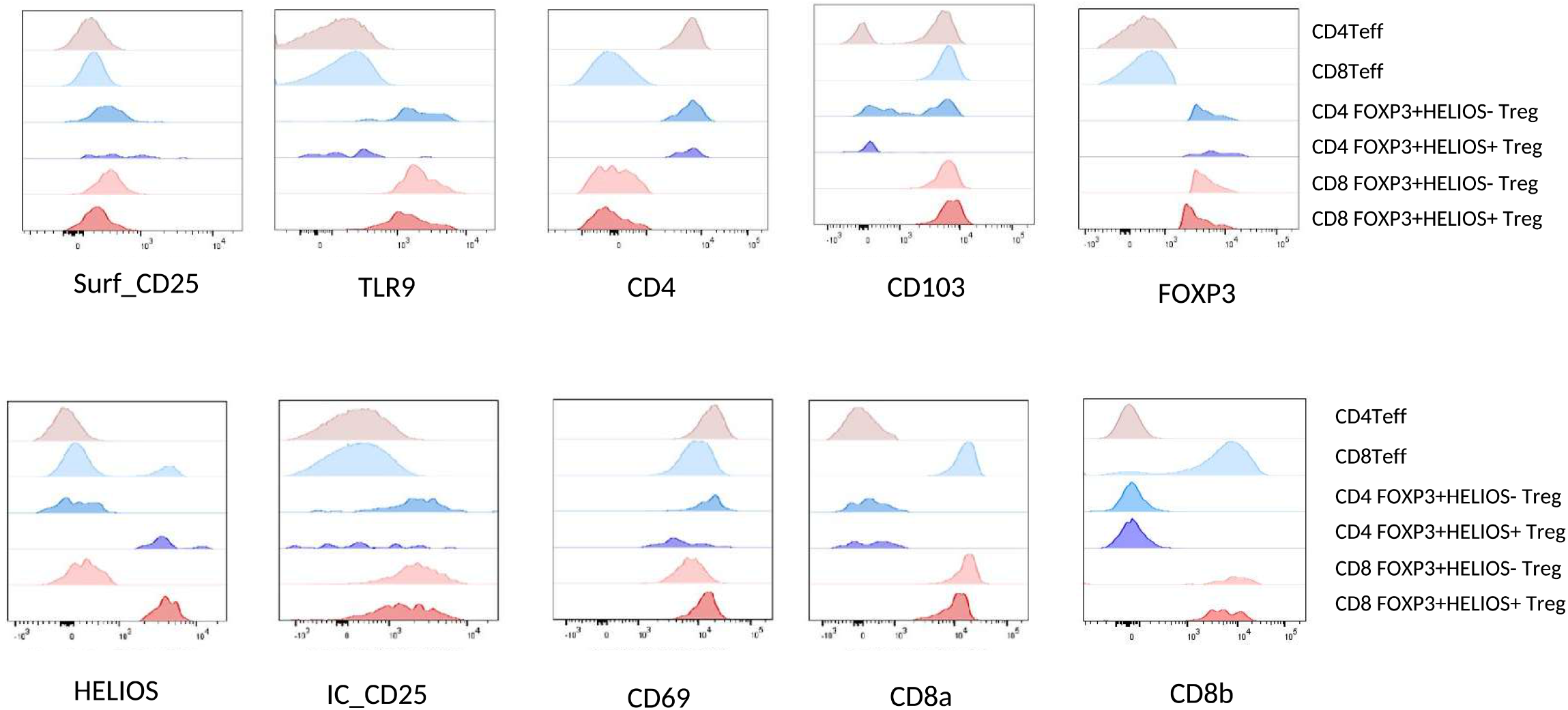
Phenotype of CD4 and CD8+ FOXP3+ HELIOS+ and HELIOS-Tregs in human tissues. Histogram overlays of gated CD4 Teff and CD8 Teff, CD4 and CD8 FOXP3+HELIOS+ Tregs and CD4 and CD8 FOXP3+HELIOS-Tregs in human jejunum IELs (representative example)

**Supplementary Fig 27.**
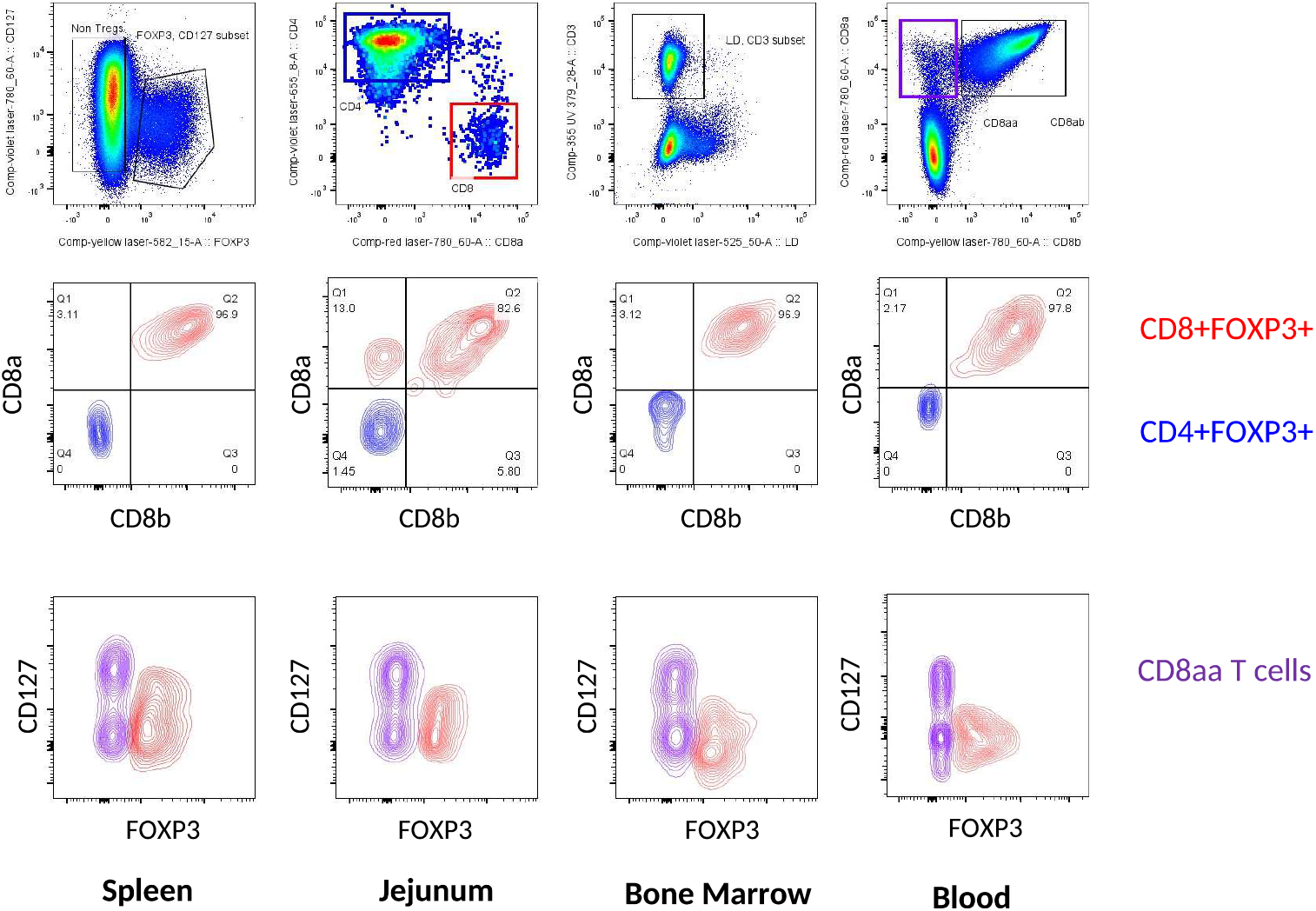
CD8ab staining of CD8+FOXP3+ Tregs. Expression of CD8a and CD8b in CD8+FOXP3+CD127low T cells from Spleen, jejunum, Bone marrow and peripheral blood. Representative plots of n = 4 spleen; n = 3 jejunum; n = 2 bone marrow and n = 2 PB

**Supplementary Fig 28.**
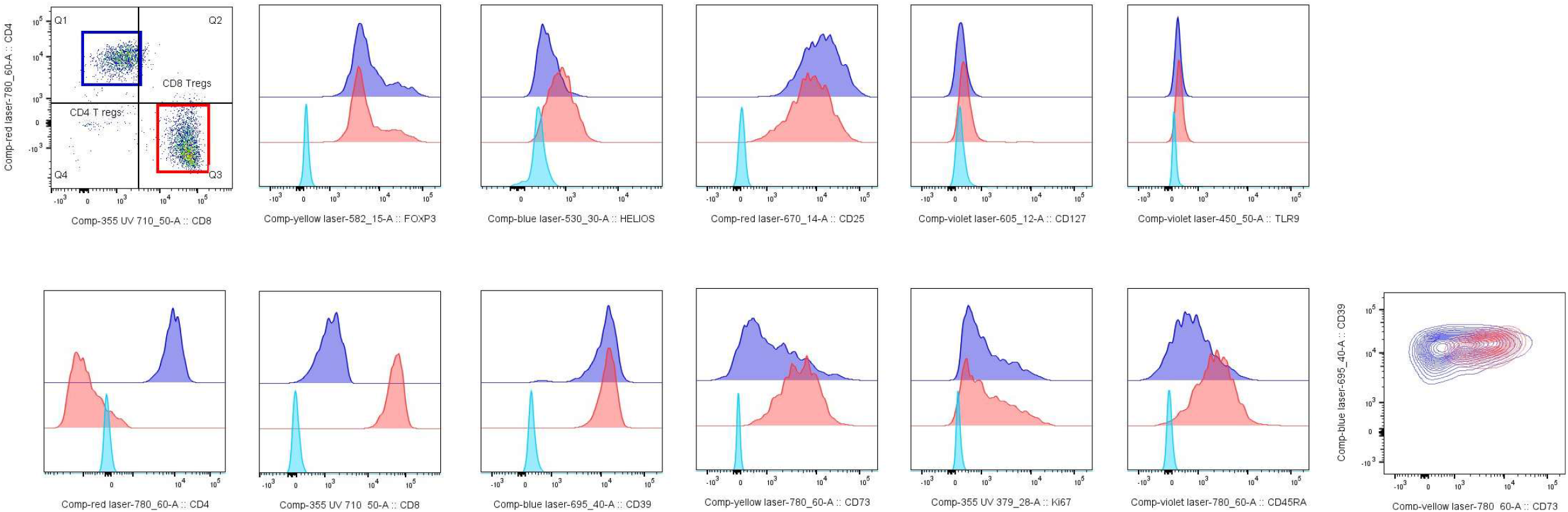
Phenotype of expTregs from spleen. Expression of Treg associated markers in splenic expanded Tregs after 3 rounds of expansion comparing CD4+ expTregs (blue) and CD8+ expTregs (red). Overlaid histograms show gated Live CD3+CD4+ or CD3+CD8+ Tregs in bulk expanded population Overlaid contour plot shows CD39 v CD73 double positivity which was greater in the CD8+ expTregs.

**Supplementary Fig 29.**
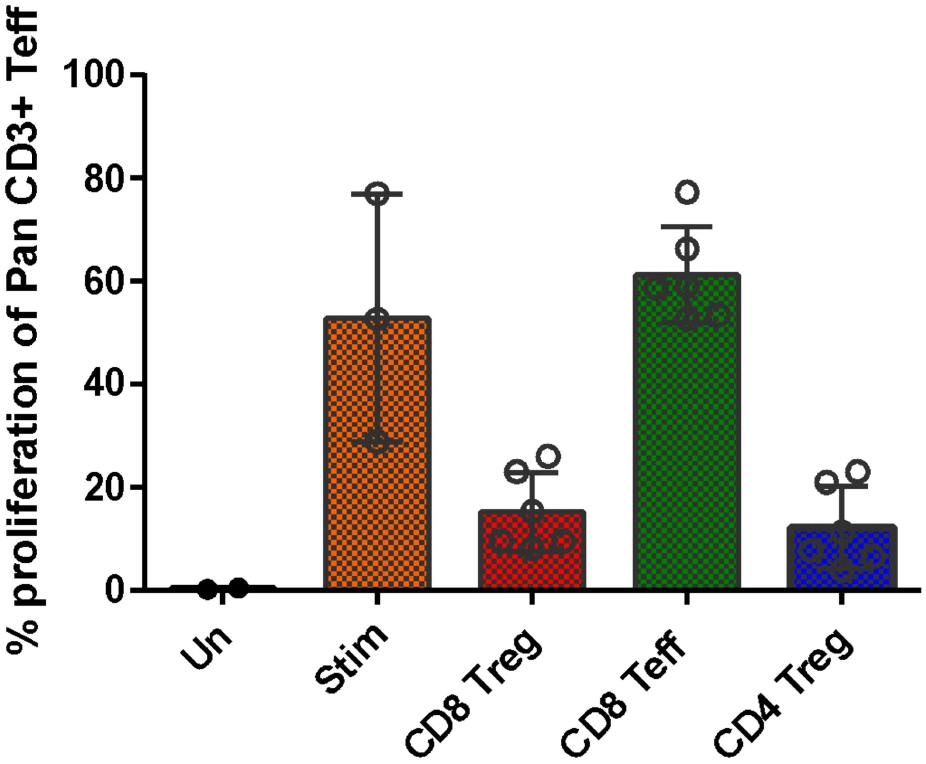
CD8 Teff controlled suppression assay. CD3+ Pan T cells were stimulated using Treg suppression inspector beads (miltenyi biotec) in the absence (orange) and presence of expanded CD8 Tregs (red), CD4 Tregs (blue) or CD8+ Teffectors (green) sorted from the same donors. Duplicates of each co-culture were performed. Donor n = 3.

**Supplementary Fig 30.**
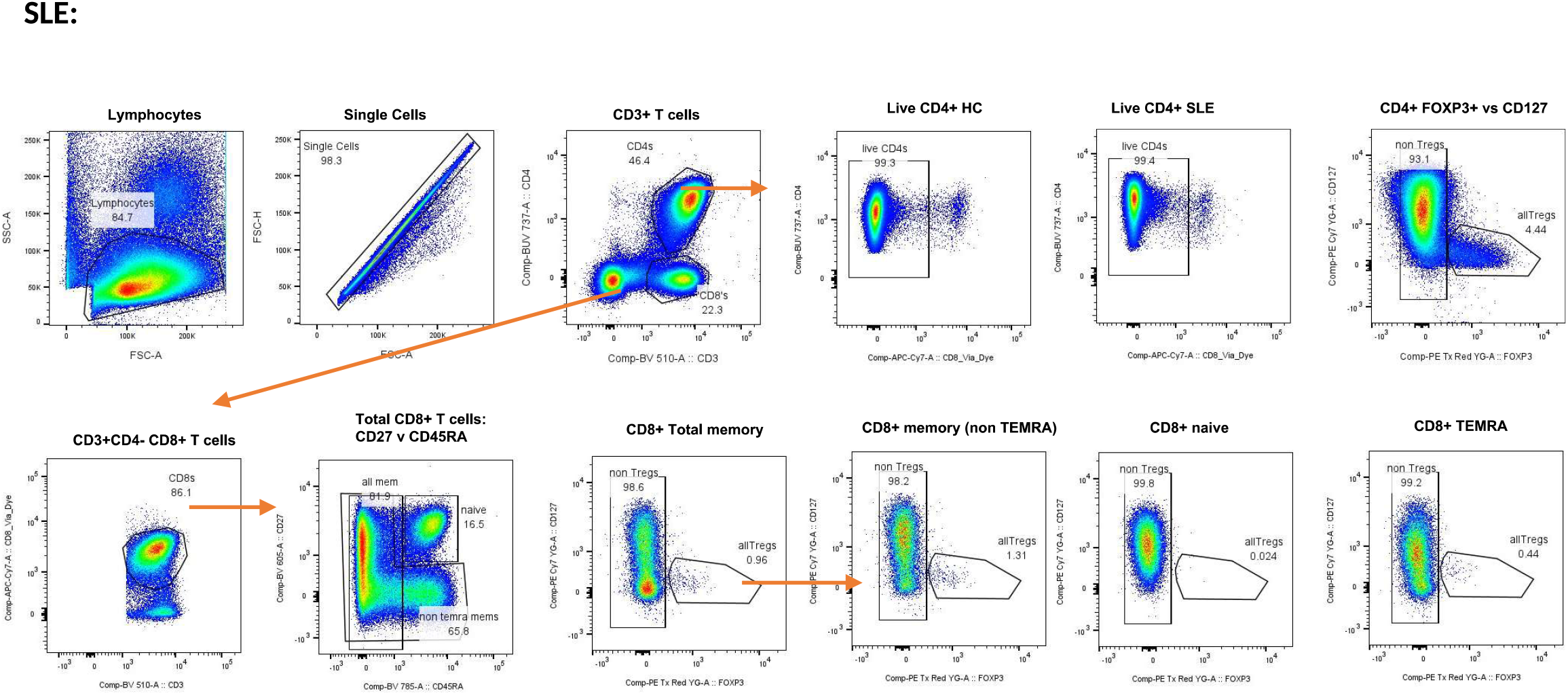
Gating strategy for SLE dataset SLE:

**Supplementary Fig 31.**
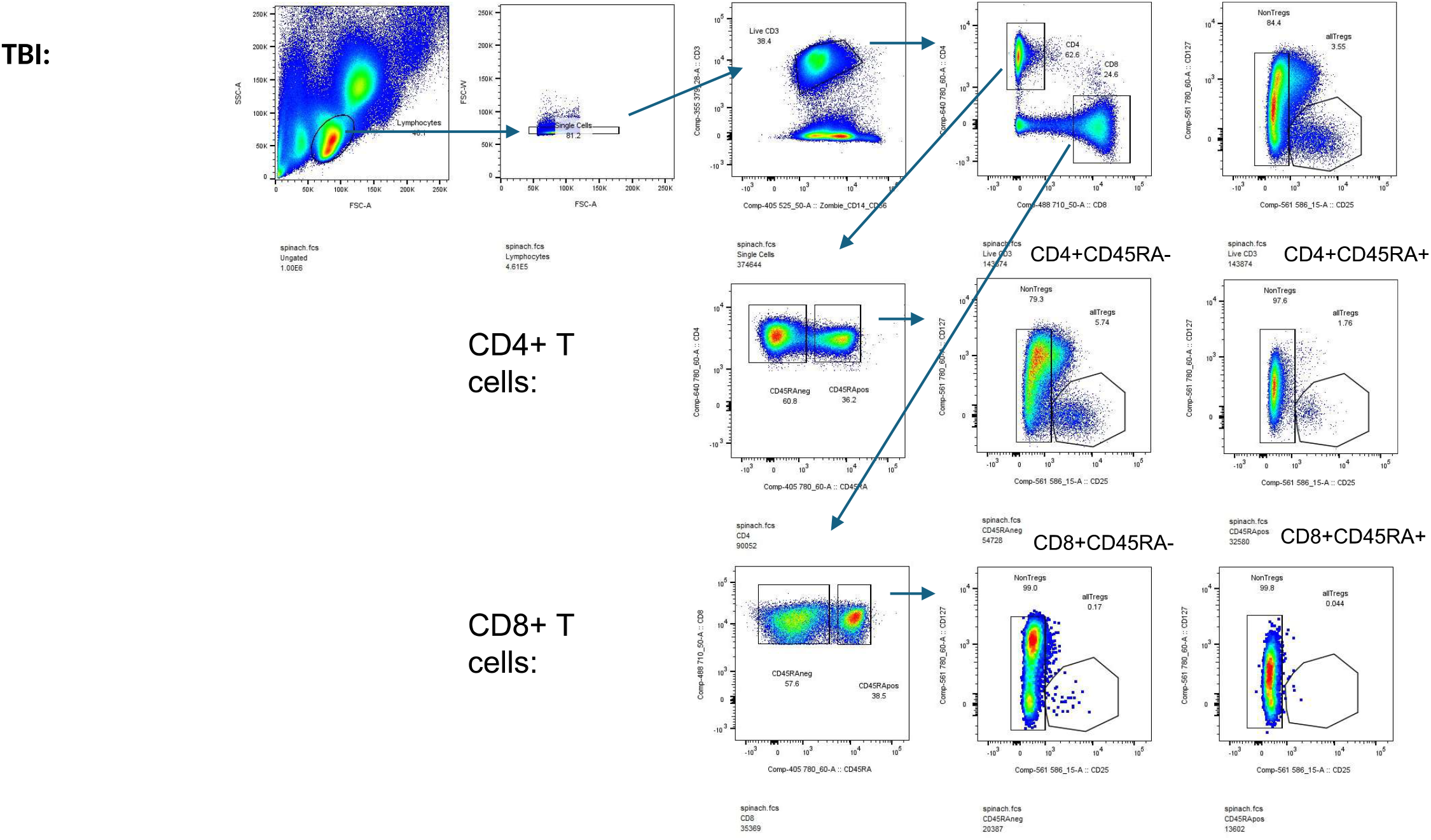
Gating strategy for TBI dataset TBI:

## References

1. J. Pohar, Q. Simon, S. Fillatreau, Antigen-Specificity in the Thymic Development and Peripheral Activity of CD4+FOXP3+ T Regulatory Cells. Front. Immunol. 9 (2018).

2. J. K. Polansky, K. Kretschmer, J. Freyer, S. Floess, A. Garbe, U. Baron, S. Olek, A. Hamann, H. von Boehmer, J. Huehn, DNA methylation controls Foxp3 gene expression. Eur. J. Immunol. 38, 1654–1663 (2008).

3. C. Hennessy, M. Deptula, J. Hester, F. Issa, Barriers to Treg therapy in Europe: From production to regulation. Front. Med. 10, 1090721 (2023).

4. M. Strioga, V. Pasukoniene, D. Characiejus, CD8+ CD28− and CD8+ CD57+ T cells and their role in health and disease. Immunology 134, 17–32 (2011).

5. M. Rifa’i, Y. Kawamoto, I. Nakashima, H. Suzuki, Essential Roles of CD8+CD122+ Regulatory T Cells in the Maintenance of T Cell Homeostasis. J. Exp. Med. 200, 1123–1134 (2004).

6. S. Bezie, D. Meistermann, L. Boucault, S. Kilens, J. Zoppi, E. Autrusseau, A. Donnart, V. Nerriere-Daguin, F. Bellier-Waast, E. Charpentier, F. Duteille, L. David, I. Anegon, C. Guillonneau, Ex Vivo Expanded Human Non-Cytotoxic CD8+CD45RClow/-Tregs Efficiently Delay Skin Graft Rejection and GVHD in Humanized Mice. Front. Immunol. 8 (2018).

7. J. Li, M. Zaslavsky, Y. Su, J. Guo, M. J. Sikora, V. van Unen, A. Christophersen, S.-H. Chiou, L. Chen, J. Li, X. Ji, J. Wilhelmy, A. M. Mcsween, B. A. Palanski, V. V. A. Mallajosyula, N. A. Bracey, G. K. R. Dhondalay, K. Bhamidipati, J. Pai, L. B. Kipp, J. E. Dunn, S. L. Hauser, J. R. Oksenberg, A. T. Satpathy, W. H. Robinson, C. L. Dekker, L. M. Steinmetz, C. Khosla, P. J. Utz, L. M. Sollid, Y.-H. Chien, J. R. Heath, N. Q. Fernandez-Becker, K. C. Nadeau, N. Saligrama, M. M. Davis, KIR+CD8+ T cells suppress pathogenic T cells and are active in autoimmune diseases and COVID-19. Science 0, eabi9591.

8. H.-J. Kim, X. Wang, S. Radfar, T. J. Sproule, D. C. Roopenian, H. Cantor, CD8+ T regulatory cells express the Ly49 Class I MHC receptor and are defective in autoimmune prone B6-Yaa mice. Proc. Natl. Acad. Sci. 108, 2010–2015 (2011).

9. C. Wang, X. Liu, Z. Li, Y. Chai, Y. Jiang, Q. Wang, Y. Ji, Z. Zhu, Y. Wan, Z. Yuan, Z. Chang, M. Zhang, CD8+NKT-like cells regulate the immune response by killing antigen-bearing DCs. Sci. Rep. 5, 14124 (2015).

10. S. Mishra, S. Srinivasan, C. Ma, N. Zhang, CD8+ Regulatory T Cell - A Mystery to Be Revealed. Front. Immunol. 12, 3374 (2021).

11. Y. Kiniwa, Y. Miyahara, H. Y. Wang, W. Peng, G. Peng, T. M. Wheeler, T. C. Thompson, L. J. Old, R.-F. Wang, CD8+ Foxp3+ Regulatory T Cells Mediate lmmunosuppression in Prostate Cancer. Clin. Cancer Res. 13, 6947–6958 (2007).

12. N. Chaput, S. Louafi, A. Bardier, F. Charlotte, J.-C. Vaillant, F. Menegaux, M. Rosenzwajg, F. Lemoine, D. Klatzmann, J. Taieb, Identification of CD8+CD25+Foxp3+ suppressive T cells in colorectal cancer tissue. Gut 58, 520–529 (2009).

13. G. Churlaud, F. Pitoiset, F. Jebbawi, R. Lorenzen, B. Bellier, M. Rosenzwajg, D. Klatzmann, Human and Mouse CD8+CD25+FOXP3+ Regulatory T Cells at Steady State and during lnterleukin-2 Therapy. Front. Immunol. 6 (2015).

14. C. E. Whyte, K. Singh, O. T. Burton, M. Aloulou, L. Kouser, R. V. Veiga, A. Dashwood, H. Okkenhaug, S. Benadda, A. Moudra, O. Bricard, S. Lienart, P. Bielefeld, C. P. Roca, F. J. Naranjo-Galindo, F. Lombard-Vadnais, S. Junius, D. Bending, T. Hochepied, T. Y. F. Halim, S. Schlenner, S. Lesage, J. Dooley, A. Liston, Context-dependent effects of IL-2 rewire immunity into distinct cellular circuits. J. Exp. Med. 219, e20212391 (2022).

15. L.B. Jarvis, M. K. Matyszak, R. C. Duggleby, J.C. Goodall, F. C. Hall, J. S. H. Gaston, Autoreactive human peripheral blood CD8+ T cells with a regulatory phenotype and function. Eur. J. Immunol. 35, 2896–2908 (2005).

16. A. Liston, M. Aloulou, A fresh look at a neglected regulatory lineage: CD8+Foxp3+ Regulatory T cells. Immunol. Lett. 247, 22–26 (2022).

17. D. Simone, F. Penkava, A. Ridley, S. Sansom, M. H. AI-Mossawi, P. Bowness, Single cell analysis of spondyloarthritis regulatory T cells identifies distinct synovial gene expression patterns and clonal fates. Commun. Biol. 4, 1395 (2021).

18. M. S. Vacchio, R. Bosselut, What Happens in the Thymus Does Not Stay in the Thymus: How T Cells Recycle the CD4+ −CD8+ Lineage Commitment Transcriptional Circuitry To Control Their Function. J. Immunol. 196, 4848–4856 (2016).

19. Y. Fujii, M. Okumura, K. lnada, K. Nakahara, H. Matsuda, CD45 isoform expression during T cell development in the thymus. Eur. J. Immunol. 22, 1843–1850 (1992).

20. P.A. Szabo, M. Miron, D. L. Farber, Location, location, location: Tissue resident memory T cells in mice and humans. Sci. Immunol. 4 (2019).

21. B. V. Kumar, W. Ma, M. Miron, T. Granat, R. S. Guyer, D. J. Carpenter, T. Senda, X. Sun, S.-H. Ho, H. Lerner, A. L. Friedman, Y. Shen, D. L. Farber, Human Tissue-Resident Memory T Cells Are Defined by Core Transcriptional and Functional Signatures in Lymphoid and Mucosal Sites. Cell Rep. 20, 2921–2934 (2017).

22. S. Koizumi, H. Ishikawa, Transcriptional Regulation of Differentiation and Functions of Effector T Regulatory Cells. Cells 8, 939 (2019).

23. S. N. Mueller, L. K. Mackay, Tissue-resident memory T cells: local specialists in immune defence. Nat. Rev. Immunol. 16, 79–89 (2016).

24. N. Sugimoto, Y.-J. Liu, DUSP4 Stabilizes FOXP3 Expression In Human Regulatory T Cells. Blood 122, 3473 (2013).

25. M. Delacher, C. D. lmbusch, A. Hotz-Wagenblatt, J.-P. Mallm, K. Bauer, M. Simon, D. Riegel, A. F. Rendeiro, S. Bittner, L. Sanderink, A. Pant, L. Schmidleithner, K. L. Braband, B. Echtenachter, A. Fischer, V. Giunchiglia, P. Hoffmann, M. Edinger, C. Bock, M. Rehli, B. Brors, C. Schmidl, M. Feuerer, Precursors for Nonlymphoid-Tissue Treg Cells Reside in Secondary Lymphoid Organs and Are Programmed by the Transcription Factor BATF. Immunity 52, 295–312.e11 (2020).

26. M. Delacher, M. Simon, L. Sanderink, A. Hotz-Wagenblatt, M. Wuttke, K. Schambeck, L. Schmidleithner, S. Bittner, A. Pant, U. Ritter, T. Hehlgans, D. Riegel, V. Schneider, F. K. Groeber-Becker, A. Eigenberger, C. Gebhard, N. Strieder, A. Fischer, M. Rehli, P. Hoffmann, M. Edinger, T. Strowig, J. Huehn, C. Schmidl, J.M. Werner, L. Prantl, B. Brors, C. D. lmbusch, M. Feuerer, Single-cell chromatin accessibility landscape identifies tissue repair program in human regulatory T cells. Immunity, doi: 10.1016/j.immuni.2021.03.007 (2021).

27. R. Zhang, K. Xu, Y. Shao, Y. Sun, J. Saredy, E. Cutler, T. Yao, M. Liu, L. Liu, C. Drummer IV, Y. Lu, F. Saaoud, D. Ni, J. Wang, Y. Li, R. Li, X. Jiang, H. Wang, X. Yang, Tissue Treg Secretomes and Transcription Factors Shared With Stem Cells Contribute to a Treg Niche to Maintain Treg-Ness With 80% Innate Immune Pathways, and Functions of lmmunosuppression and Tissue Repair. Front. Immunol. 11 (2021).

28. C. Dominguez Conde, C. Xu, L. B. Jarvis, D. B. Rainbow, S. B. Wells, T. Gomes, S. K. Howlett, O. Suchanek, K. Polanski, H. W. King, L. Mamanova, N. Huang, P.A. Szabo, L. Richardson, L. Bolt, E. S. Fasouli, K. T. Mahbubani, M. Prete, L. Tuck, N. Richoz, Z. K. Tuong, L. Campos, H. S. Mousa, E. J. Needham, S. Pritchard, T. Li, R. Elmentaite, J. Park, E. Rahmani, D. Chen, D. K. Menon, O. A. Bayraktar, L. K. James, K. B. Meyer, N. Yosef, M. R. Clatworthy, P.A. Sims, D. L. Farber, K. Saeb-Parsy, J. L. Jones, S. A. Teichmann, Cross-tissue immune cell analysis reveals tissue-specific features in humans. Science 376, eabl5197 (2022).

29. THE TABULA SAPIENS CONSORTIUM, The Tabula Sapiens: A multiple-organ, single-cell transcriptomic atlas of humans. Science 376, eabl4896 (2022).

30. V. Coppard, G. Szep, Z. Georgieva, S. K. Howlett, L.B. Jarvis, D. B. Rainbow, O. Suchanek, E. J. Needham, H. S. Mousa, D. K. Menon, F. Feyertag, K. T. Mahbubani, K. Saeb-Parsy, J. L. Jones, FlowAtlas: an interactive tool for high-dimensional immunophenotyping analysis bridging FlowJo with computational tools in Julia. Front. Immunol. 15 (2024).

31. R. C. Ferreira, H. Z. Simons, W. S. Thompson, D. B. Rainbow, X. Yang, A. J. Cutler, J. Oliveira, X. Castro Dopico, D. J. Smyth, N. Savinykh, M. Mashar, T. J. Vyse, D. B. Dunger, H. Baxendale, A. Chandra, C. Wallace, J. A. Todd, L. S. Wicker, M. L. Pekalski, Cells with Treg-specific FOXP3 demethylation but low CD25 are prevalent in autoimmunity. J. Autoimmun. 84, 75–86 (2017).

32. S. B. Wells, D. B. Rainbow, M. Mark, P. A. Szabo, C. Ergen, A. R. Maceiras, D. P. Caron, E. Rahmani, E. Benuck, V. V. P. Amiri, D. Chen, A. Wagner, S. K. Howlett, L.B. Jarvis, K. L. Ellis, M. Kubota, R. Matsumoto, K. Mahbubani, K. Saeb-Parsy, C. Dominguez-Conde, L. Richardson, C. Xu, S. Li, L. Mamanova, L. Bolt, A. Wilk, S. A. Teichmann, D. L. Farber, P. A. Sims, J. L. Jones, N. Yosef, Multimodal profiling reveals tissue-directed signatures of human immune cells altered with age. bioRxiv [Preprint] (2024). 10.1101/2024.01.03.573877.

33. A. Marcais, C.-A. Coupet, T. Walzer, M. Tomkowiak, R. Ghittoni, J. Marvel, Cell-Autonomous CCL5 Transcription by Memory CD8 T Cells Is Regulated by IL-41. J. Immunol. 177, 4451–4457 (2006).

34. H. Ham, J. B. Hirdler, D. T. Bihnam, Z. Mao, J. K. Gicobi, B. G. Macedo, M. F. Rodriguez-Quevedo, D. F. Schultz, C. Correia, J. Zhong, K. E. Martinez, A. Banuelos, D.S. Ashton, A. B. Lagnado, R. Guo, R. Pessoa, A. Pandey, H. Li, F. Lucien, H. Borges da Silva, H. Dong, D. D. Billadeau, Lysosomal NKG7 restrains mTORC1 activity to promote CD8+ T cell durability and tumor control. Nat. Commun. 16, 1628 (2025).

35. P.A. Szabo, H. M. Levitin, M. Miron, M. E. Snyder, T. Senda, J. Yuan, Y. L. Cheng, E. C. Bush, P. Dogra, P. Thapa, D. L. Farber, P. A. Sims, Single-cell transcriptomics of human T cells reveals tissue and activation signatures in health and disease. Nat. Commun. 10 (2019).

36. A. Baeyens, V. Fang, C. Chen, S. R. Schwab, Exit strategies: S1P signaling and T cell migration. Trends Immunol. 36, 778–787 (2015).

37. Resource, 10X Genomics. https://www.10xgenomics.com/welcome.

38. Y. C. Kim, R. Bhairavabhotla, J. Yoon, A. Golding, A. M. Thornton, D. Q. Tran, E. M. Shevach, Oligodeoxynucleotides stabilize Helios-expressing Foxp3+ human T regulatory cells during in vitro expansion. Blood 119, 2810–2818 (2012).

39. F. Sallusto, D. Lenig, R. Forster, M. Lipp, A. Lanzavecchia, Two subsets of memory T lymphocytes with distinct homing potentials and effector functions. Nature 401, 708–712 (1999).

40. D. L. Woodland, J. E. Kohlmeier, Migration, maintenance and recall of memory T cells in peripheral tissues. Nat. Rev. Immunol. 9, 153–161 (2009).

41. L.B. Jarvis, D. B. Rainbow, V. Coppard, S. K. Howlett, Z. Georgieva, J. L. Davies, H. K. Mullay, J. Hester, T. Ashmore, A. Van Den Bosch, J. T. Grist, A. J. Coles, H. S. Mousa, S. Pluchino, K. T. Mahbubani, J. L. Griffin, K. Saeb-Parsy, F. Issa, L. Peruzzotti-Jametti, L. S. Wicker, J. L. Jones, Therapeutically expanded human regulatory T-cells are super-suppressive due to HIF1A induced expression of CD73. Commun. Biol. 4, 1186 (2021).

42. A. M. Thornton, P. E. Karty, D. Q. Tran, E. A. Wohlfert, P. E. Murray, Y. Belkaid, E. M. Shevach, Expression of Helios, an lkaros transcription factor family member, differentiates thymic-derived from peripherally induced Foxp3+ T regulatory cells. J. Immunol. Baltim. Md 1950184, 3433–3441 (2010).

43. A. M. Thornton, J. Lu, P. E. Karty, Y. C. Kim, C. Martens, P. D. Sun, E. M. Shevach, Helios+ and Helios Treg subpopulations are phenotypically and functionally distinct and express dissimilar TCR repertoires. Eur. J. Immunol. 49, 398–412 (2019).

44. L. Morina, M. E. Jones, C. Oguz, M. J. Kaplan, A. Gangaplara, C. D. Fitzhugh, C. G. Kanakry, E. M. Shevach, M. Buszko, Co-expression of Foxp3 and Helios facilitates the identification of human T regulatory cells in health and disease. Front. Immunol. 14, 1114780 (2023).

45. J. Jacobse, J. Li, E. H. H. M. Rings, J. N. Samsom, J. A. Goettel, Intestinal Regulatory T Cells as Specialized Tissue-Restricted Immune Cells in Intestinal Immune Homeostasis and Disease. Front. Immunol. 12 (2021).

46. J. Choi, B.-R. Kim, B. Akuzum, L. Chang, J.-Y. Lee, H.-K. Kwon, TREGking From Gut to Brain: The Control of Regulatory T Cells Along the Gut-Brain Axis. Front. Immunol. 13 (2022).

47. J.B. Wing, Y. Kitagawa, M. Locci, H. Hume, C. Tay, T. Morita, Y. Kidani, K. Matsuda, T. Inoue, T. Kurosaki, S. Crotty, C. Coban, N. Ohkura, S. Sakaguchi, A distinct subpopulation of CD25-T-follicular regulatory cells localizes in the germinal centers. Proc. Natl. Acad. Sci. U. S. A. 114, E6400–E6409 (2017).

48. A. M. Krieg, TLR9 and DNA “feel” RAGE. Nat. Immunol. 8, 475–477 (2007).

49. J. Tian, A. M. Avalos, S.-Y. Mao, B. Chen, K. Senthil, H. Wu, P. Parroche, S. Drabic, D. Golenbock, C. Sirois, J. Hua, L. L. An, L. Audoly, G. La Rosa, A. Bierhaus, P. Naworth, A. Marshak-Rothstein, M. K. Crow, K. A. Fitzgerald, E. Latz, P.A. Kiener, A. J. Coyle, Toll-like receptor 9-dependent activation by DNA-containing immune complexes is mediated by HMGB1 and RAGE. Nat. Immunol. 8, 487–496 (2007).

50. S. Ivanov, A.-M. Dragoi, X. Wang, C. Dallacosta, J. Louten, G. Musco, G. Sitia, G. S. Yap, Y. Wan, C. A. Biron, M. E. Bianchi, H. Wang, W.-M. Chu, A novel role for HMGB1 in TLR9-mediated inflammatory responses to CpG-DNA. Blood 110, 1970–1981 (2007).

51. J. A. Hall, N. Bouladoux, C. M. Sun, E. A. Wohlfert, R. B. Blank, Q. Zhu, M. E. Grigg, J. A. Berzofsky, Y. Belkaid, Commensal DNA Limits Regulatory T Cell Conversion and Is a Natural Adjuvant of Intestinal Immune Responses. Immunity 29, 637–649 (2008).

52. K. H. G. Mills, TLR9 Turns the Tide on Treg Cells. Immunity 29, 518–520 (2008).

53. Z. Urry, E. Xystrakis, D. F. Richards, J. McDonald, Z. Sattar, D. J. Cousins, C. J. Corrigan, E. Hickman, Z. Brown, C. M. Hawrylowicz, Ligation of TLR9 induced on human IL-10-secreting Tregs by 1a,25-dihydroxyvitamin 03 abrogates regulatory function. J. Clin. Invest. 119, 387–398 (2009).

54. M.A. Alikhan, S. A. Summers, P. Y. Gan, A. J. Chan, M. B. Khouri, J. D. Ooi, J. R. Ghali, D. Odobasic, M. J. Hickey, A. R. Kitching, S. R. Holdsworth, Endogenous Toll-Like Receptor 9 Regulates AKI by Promoting Regulatory T Cell Recruitment. J. Am. Soc. Nephrol. 27, 706–714 (2016).

55. H. Schmitt, J. Ulmschneider, U. Billmeier, M. Vieth, P. Scarozza, S. Sonnewald, S. Reid, I. Atreya, T. Rath, S. Zundler, M. Langheinrich, J. Schuttler, A. Hartmann, T. Winkler, C. Admyre, T. Knittel, C. Dieterich Johansson, A. Zargari, M. F. Neurath, R. Atreya, The TLR9 Agonist Cobitolimod Induces IL10-Producing Wound Healing Macrophages and Regulatory T Cells in Ulcerative Colitis. J. Crohns Colitis 14, 508–524 (2020).

56. R. Atreya, L. Peyrin-Biroulet, A. Klymenko, M. Augustyn, I. Bakulin, D. Slankamenac, P. Miheller, A. Gasbarrini, X. Hebuterne, K. Arnesson, T. Knittel, J. Kowalski, M. F. Neurath, W. J. Sandborn, W. Reinisch, CONDUCT study group, Cobitolimod for moderate-to-severe, left-sided ulcerative colitis (CONDUCT): a phase 2b randomised, double-blind, placebo-controlled, dose-ranging induction trial. Lancet Gastroenterol. Hepatol. 5, 1063–1075 (2020).

57. F. Noyan, K. Zimmermann, M. Hardtke-Wolenski, A. Knoefel, E. Schulde, R. Geffers, M. Hust, J. Huehn, M. Galla, M. Morgan, A. Jokuszies, M. P. Manns, E. Jaeckel, Prevention of Allograft Rejection by Use of Regulatory T Cells With an MHC-Specific Chimeric Antigen Receptor. Am. J. Transplant. Off. J. Am. Soc. Transplant. Am. Soc. Transpl. Surg. 17, 917–930 (2017).

58. S. Bezie, B. Charreau, N. Vimond, J. Lasselin, N. Gerard, V. Nerriere-Daguin, F. Bellier-Waast, F. Duteille, Anegon, C. Guillonneau, Human CD8+ Tregs expressing a MHC-specific CAR display enhanced suppression of human skin rejection and GVHD in NSG mice. Blood Adv. 3, 3522–3538 (2019).

59. D. Rainbow, S. Howlett, L. Jarvis, J. Jones, Multi tissue processing for single cell sequencing of human immune cells. (2021).

60. E. A. Moskalev, M. G. Zavgorodnij, S. P. Majorova, I. A. Vorobjev, P. Jandaghi, I. V. Bure, J. D. Hoheisel, Correction of PCR-bias in quantitative DNA methylation studies by means of cubic polynomial regression. Nucleic Acids Res. 39, e77 (2011).

61. P. M. Warnecke, C. Stirzaker, J. R. Melki, D.S. Millar, C. L. Paul, S. J. Clark, Detection and measurement of PCR bias in quantitative methylation analysis of bisulphite-treated DNA. Nucleic Acids Res. 25, 4422–4426 (1997).

62. J. T. Grist, L. B. Jarvis, Z. Georgieva, S. Thompson, H.K. Sandhu, K. Burling, A. Clarke, S. Jackson, M. Wills, F. A. Gallagher, J. L. Jones, Extracellular Lactate: A Novel Measure of T Cell Proliferation. J. Immunol., doi: 10.4049/jimmunol.1700886 (2017).

63. G. Adigbli, S. Menoret, A. R. Cross, J. Hester, F. Issa, I. Anegon, Humanization of Immunodeficient Animals for the Modeling of Transplantation, Graft Versus Host Disease and Regenerative Medicine. Transplantation Online First (2020).

64. F. A. Wolf, P. Angerer, F. J. Theis, SCANPY: large-scale single-cell gene expression data analysis. Genome Biol. 19, 15 (2018).

65. G. Palla, H. Spitzer, M. Klein, D. Fischer, A. C. Schaar, L.B. Kuemmerle, S. Rybakov, I. L. Ibarra, O. Holmberg, I. Virshup, M. Lotfollahi, S. Richter, F. J. Theis, Squidpy: a scalable framework for spatial omics analysis. Nat. Methods 19, 171–178 (2022).

66. R. Massoni-Badosa, S. Aguilar-Fernandez, J.C. Nieto, P. Soler-Vila, M. Elosua-Bayes, D. Marchese, M. Kulis, A. Vilas-Zornoza, M. M. Buhler, S. Rashmi, C. Alsinet, G. Caratu, C. Moutinho, S. Ruiz, P. Lorden, G. Lunazzi, D. Colomer, G. Frigola, W. Blevins, L. Romero-Rivero, V. Jimenez-Martinez, A. Vidal, J. Mateos-Jaimez, A. Maiques-Diaz, S. Ovejero, J. Moreaux, S. Palomino, D. Gomez-Cabrero, X. Agirre, M. A. Weniger, H. W. King, L. C. Garner, F. Marini, F. J. Cervera-Paz, P. M. Baptista, I. Vilaseca, C. Rosales, S. Ruiz-Gaspa, B. Talks, K. Sidhpura, A. Pascual-Reguant, A. E. Hauser, M. Haniffa, F. Prosper, R. Kuppers, I. G. Gut, E. Campo, J. I. Martin-Subera, H. Heyn, An atlas of cells in the human tonsil. Immunity 57, 379–399.e18 (2024).

67. K. Blighe, kevinblighe/EnhancedVolcano, (2020); https://github.com/kevinblighe/EnhancedVolcano.

68. Renesh Bedre, reneshbedre/bioinfokit: Bioinformatics data analysis and visualization toolkit, Zenodo (2020); 10.5281/zenodo.3841708.

