## Supplementary Table 1 for "Hiding in Plain Sight: CD8+FOXP3+Tregs sequester CD25 and are enriched in human tissues"

| Donor | Donor_code | DCD/DBD | Cause of death | Age bin | Sex | Screened infections (positive) | Tissues acquired |
|---|---|---|---|---|---|---|---|
| 1 | 245C | DCD | ICH | 65-69 | M | EBV+ CMV+ | BM, LIV, LN |
| 2 | 258B | DCD | ICH | 75-79 | M | EBV+ CMV+ | BM, LN, PB, SPL |
| 3 | 262C | DCD | RTA | 25-29 | F | EBV+ | MLN, SPL, THY |
| 4 | 267C | DCD | HBD | 20-24 | M | | SPL, THY |
| 5 | 295B | DCD | ICH | 75-79 | F | EBV+ | SPL, TLN |
| 6 | 390C | DCD | ICH | 65-69 | F | EBV+ CMV+ | BM, LIV, SPL |
| 7 | 403C | DCD | ICH | 50-54 | M | EBV+ CMV+ | BM, SPL |
| 8 | 412C | DCD | ICH | 70-74 | M | EBV+ Toxo+ | BM, ILE, LIV, MLN, PB, SPL, TLN |
| 9 | 423C | DCD | ICH | 60-64 | M | EBV+ | BM, ILE, KID, LIV, MLN, LNG, PB, SPL, TLN |
| 10 | 428C | DCD | ICH | 55-59 | F | EBV+ | BM, SPL |
| 11 | 470BR | DBD | HBD | 18-24 | M | | BM, PB, SPL, THY |
| 12 | 266C (NRP) | DCD | HBD | 70-74 | F | | LNG |
| 13 | 011018_K197 | DNA | DNA | DNA | DNA | SU | KID |
| 14 | 1B18-807/281018_K1 | DBD | DNA | 60-64 | F | SU | KID, LN, SPL |
| 15 | 411C | DCD | HBD | 25-29 | M | EBV+ CMV+ | BM, SPL |
| 16 | 150918K (K193) | DNA | DNA | DNA | DNA | SU | KID |
| 17 | 190918K (K194) | DNA | DNA | DNA | DNA | SU | KID |
| 18 | 704C (NRP) | DCD | ICH | 50-54 | M | | FAT, JEJ_IELs, JEJ_LP, LNG, PB |
| 19 | 759B | DBD | CA | 18-24 | F | EBV+ | BM, FAT, JEJ_IELs, PB, SPL, TLN |
| 20 | 768B | DBD | HBD | 18-24 | M | EBV+ | BM, FAT, JEJ_IELs, JEJ_LPs, LN, LNG, MLN, PB, SPL, THY |
| 21 | 808B | DBD | ICH | 45-49 | M | EBV+ | BM, JEJ_IELs, JEJ_LP, MLN, PB, SPL, TLN |
| 22 | 591C | DCD | ICH | 35-39 | M | CMV+ | BM, MLN, TLN, |
| 23 | 637C | DCD | ICH | 50-54 | M | EBV+ CMV+ | BM, TLN |
| 24 | 538B | DBD | ICH | 25-29 | M | CMV+ | BM, SPL, LN |
| 25 | 640C | DCD | ICH | 70-74 | F | EBV+ CMV+ | BM, SPL, LN |
| 26 | 852C | DCD | RTA | 65-69 | M | EBV+ CMV+ | BM, JEJ_LP, LIV, MLN, SPL, TLN |

| Key | |
|---|---|
| DCD | Donation after Circulatory Death |
| DBD | Dontion after Brain Death |
| SPL | Spleen |
| KID | Kidney |
| BM | Bone marrow |
| LNG | Lung |
| LIV | Liver |
| ILE | ileum |
| JEJ_LP | Jejunum lamina propria prep |
| JEJ_IEL | Jejunum intraepithelia prep |
| MLN | Mesenteric lymph node |
| THY | Thymus |
| TLN | Thoracic lymph node |
| FAT | Fat (abdominal) |
| PB | Peripheral blood |
| CMV | Cytomegalovirus |
| EBV | Epstein Barr Virus |
| Toxo | Toxoplasmosis |
| RTA | road traffic accident |
| DNA | Data not available |
| ICH | Intracranial haemorrhage |
| HBD | Hypoxic brain damage |
| CA | Cardiac arrest |
| SU | Status unknown |
