## Supplementary Table 2 for "Hiding in Plain Sight: CD8+FOXP3+Tregs sequester CD25 and are enriched in human tissues"

| Supplementary Table_2 | | | | | | | | | | | | |
|---|---|---|---|---|---|---|---|---|---|---|---|---|
| **Corresponding figures** | | | Fig 1 | Fig 1 | Fig2 | Fig 2 | Fig FLOWATLAS & Fig3/4 | Fig 3/4 | Fig 3/4 | Fig 3/4 | | Fig 3/4 |
| **Laser** | | **ORGAN/panel no.** | **BLOOD_1** | **BLOOD_2** | **THYMUS** | **CORD BLOOD** | **TISSUE 1** | **TISSUE 2** | **TISSUE 3** | **TISSUE 4** | **TISSUE 5** | **TISSUE 6** |
| 355 | 379/28 | BUV395 | Ki67* (B56) | Ki67* (B56) | CD103 (Ber-ACT8) | CD103 (Ber-ACT8) | CD103 (Ber-ACT8) | CD3 (UCHT1) | CD103 (Ber-ACT8) | CD103 (Ber-ACT8) | CD103 (Ber-ACT8) | CD103 (Ber-ACT8) |
| | 515/30 | Zuv (515/30) | | | | | | zombie LD | | | | |
| | 560/40 | BUV563 | | | | | | CD8 (RPA-T8) | | | | |
| | 670/30 | BUV661 | | | | | | | | | | |
| | 740/35 | BUV737 | CD49d (9F10) | CD49d (9F10) | CD69 (FN50) | CD8 (SK1) | CD69 (FN50) | CD69 (FN50) | CD69 (FN50) | CD69 (FN50) | CD69 (FN50) | CD69 (FN50) |
| | 820/60 | BUV805 | | | | | | CD4 (RPA-T4) | | | | |
| 405 | 450/50 | BV421 | PD-1 (MIH4) | CD39 (Tu66) | CD25 (2 clones:2A3/MA2.1) | | CD25 (2 clones:2A3/MA2.1) | CD103 (Ber-ACT8) | TLR9 (S16013D) | TLR9 (S16013D) | TLR9 (S16013D) | TLR9 (S16013D) |
| | 515/20 | BV510 | CD31 (WM59) | CD31 (WM59) | CD45RA (HI100) | Zombie aqua | Zombie aqua | HLA-DR (G46-6) | Zombie aqua | Zombie aqua | Zombie aqua | LD Aqua/56 dump |
| | 585/15 | BV570 | CD3 (UCHT1) | CD3 (UCHT1) | | | CD3 (UCHT1) | | | | | |
| | 605/40 | BV605 | CD27 (L128) | CD45RA (HI100) | | | CD127 (HIL-7R-M21) | CCR4 (L291H4) | | | | |
| | 670/30 | BV650 | | | CD4 (SK3) | CD4 (SK3) | CD4 (SK3) | CCR6 (11A9) | CD3 (UCHT1) | CD3 (UCHT1) | CD3 (UCHT1) | CD3 (UCHT1) |
| | 710/40 | BV711 | CD15s (FH6) | CD15s (FH6) | | | | PD-1 (MIH4) | | | | |
| | 780/60 | BV786 | CD45RA (HI100) | CD73 (AD2) | CD5 SB780 (UCHT2) | CD45RA (HI100) | CD45RA (HI100) | CD45RA (HI100) | CD45RA (HI100) | CD127 (eBioRDR5) | CD127 (eBioRDR5) | CD45RA (HI100) |
| 488 | 525/50 | FITC/BB515 | HELIOS (22F6) | HELIOS (22F6) | FOXP3 (PCH1101/259D) | CD25 (2 clones:2A3/MA2.1) | CD45RC (MT2) | CCR10 (1B5) | CD25 x 2 (2A3/MA2.1) | CD25 x 2 (2A3/MA2.1) | CD25 x 2 (2A3/MA2.1) | CD25 x 2 (2A3/MA2.1) |
| | 715/30 | BB700/PerCPeFl710 | TIGIT (MBSA43) | TIGIT (MBSA43) | TCRab (IP26) | TCRab (IP26) | TIGIT (MBSA43) | CXCR3 (G025H7) | CD8 (RPA-T8) | CD8 (RPA-T8) | CD8 (RPA-T8) | CD8 (RPA-T8) |
| 561 | 585/15 | PE | CD226 (11A8) | CD226 (11A8) | ThPOK (11H11A14) | FOXP3 (PCH1101/259D) | FOXP3 (PCH1101/259D) | FOXP3 (PCH1101/259D) | FOXP3 (PCH1101/259D) | FOXP3 (PCH1101/259D) | FOXP3 (PCH1101/259D) | FOXP3 (PCH1101/259D) |
| | 610/20 | PE-Dazzle | FOXP3 (2 clones: PCH1101/259D) | FOXP3 (2 clones: PCH1101/259D) | HELIOS (22F6) | HELIOS (22F6) | CTLA4 (BN13) | HELIOS (22F6) | HELIOS (22F6) | HELIOS (22F6) | HELIOS (22F6) | HELIOS (22F6) |
| | 670/30 | PEcy5 | CTLA-4 (BN13) | CTLA-4 (BN13) | | | | | | | | |
| | 710/50 | PEcy5.5 | CD8 (RPA-T8) | CD8 (RPA-T8) | | | | | | | | |
| | 780/60 | PEcy7 | CD127 (ebioRDR5) | CD127 (ebioRDR5) | CD8 (SK1) | CD127 (ebioRDR5) | CD8 (SK1) | CD127 (ebioRDR5) | CD27 PEvio770 (323) | CXCR6 (K041E5) | CXCR6 (K041E5) | CD127 (ebioRDR5) |
| 640 | 670/30 | APC | CD25 (2 clones:2A3/MA2.1) | CD25 (2 clones:2A3/MA2.1) | RUNX3 (527327) | | HELIOS (22F6) | CD25 (2 clones:2A3/MA2.1) | CD25 x 2 (2A3/MA2.1) | CD25 x 2 (2A3/MA2.1) | | CD27 (M-T271) |
| | 730/35 | APCR700 | CD4 (RPA-T4) | CD4 (RPA-T4) | | | CD27 (M-T271) | CXCR5 (RF8B2) | | CD4R718 (SK3) | CD4R718 (SK3) | |
| | 780/60 | APCcy7/Fire750 | Zombie NIR | Zombie NIR | Zombie NIR | CD127 (A019D5) | CD226 (11A8) | CCR7 (G043H7) | CD127 (A019D5) | KIR x2 (DX27) & (DX9) | KIR x2 (DX27) & (DX9) | CD4 (RPA-T4) |
| | | **sample type** | **PBMC** | **PBMC** | **Thymus MNCs** | **Cord blood MNCs** | **Tissue MNCs** | **Tissue MNCs** | **Tissue MNCs** | **Tissue MNCs** | **Tissue MNCs** | **Tissue MNCs** |

Target (clone name in brackets)

Intracellular stain marked in red

**Tissue MNCs Analysis:**

| | |
|---|---|
| TISSUE 1 | Cross sectional 46 tissues from 17 donors used for poprortions of CD9 vs CD4 Tregs in tissues and FLOWATLAS analysis |
| TISSUE 2 | Cross sectional 22 tissues from 5 donors: used for proporitions of CD8 vs CD4 Tregs in tissues. |
| TISSUE 3-6 | Individual donors matched tissues, used for proportions of CD8 vs CD4 Tregs and for Tissue Treg phenotyping |
